# Unlocking the venom vault: Museum venomics reveals an untapped biochemical archive in natural history collections

**DOI:** 10.64898/2026.05.22.727068

**Authors:** Damien Esquerré, J. Scott Keogh, Daniel Dashevsky, Joseph Boileau, Adam Carroll, Nathan Dunstan, Alexander S. Mikheyev

## Abstract

Venoms are powerful weapons that shape ecological interactions across the animal kingdom. They also have high medical importance, causing thousands of human fatalities annually, while offering a rich resource for drug discovery. Despite this, considerable logistical, safety and ethical challenges mean only a fraction of the world’s venoms have been profiled quantitatively. Natural history collections offer opportunities to greatly expand our knowledge of venom systems. We used quantitative proteomic mass spectrometry on preserved venom glands, spanning 0–57 years in age, from 37 venomous snake species (32 elapids and 5 viperids), alongside fresh venom samples from most of the same taxa. Preserved glands and fresh venoms from the same species showed strong concordance in venom composition across multiple metrics. Critically, specimen age did not degrade data quality, indicating that decades-old material yields reliable quantitative profiles, opening the possibility for vast quantities of existing museum specimens to be used. When we integrate our data with published profiles, we confirm known elapid vs viper broad diversity patterns while revealing substantial Australasian elapid venom diversity. Our findings demonstrate the potential of natural history collections as a vast, largely untapped biochemical archive for high-throughput “museum venomics,” enabling evolutionary and temporal analyses of venom diversity at unprecedented scale.

## Introduction

Venoms are among the most diverse and powerful weapons to have evolved in the history of life, playing critical ecological roles in defence and predation ^1–4^. Venoms are typically complex cocktails of components that originate via gene duplication from endogenous proteins, which are subsequently weaponised ^5–7^. Released from the stabilising selection on housekeeping proteins, venoms display extremely fast rates of evolution ^8^. Rapid evolution, combined with over 100 independent origins over the last 600 million years ^9,10^, has resulted in a plethora of toxins that incapacitate victims in countless ways, from disabling blood clotting to blocking synaptic transmission ^10,11^. The specificity and potency that makes toxins lethal also makes them valuable for drug discovery, and many are already proving useful to treat conditions such as cancer, arthritis, diabetes, obesity, and pain ^12–15^. At the same time, snakebite remains a major medical issue, causing >100,000 deaths and hundreds of thousands of permanent disabilities annually ^16–18^.

Despite the evolutionary and medical relevance of venoms, the sheer diversity of their component toxins, and the challenges and risks associated with obtaining venom from live animals, means that quantitative profiles exist for only a small fraction of venomous species ^19^. Having comprehensive knowledge of the biochemical diversity found in chemical arsenals across the tree of life has life-saving implications ^18^. Knowing the relative abundance of the different toxins found in dangerous animals can provide guidance for available species-specific anti-venom administration. Novel venom diversity can be mined for bioactive compounds with translational value such as therapeutics and pesticides ^20–22^. The rapid evolution of venoms, and their role as the primary mechanism of prey acquisition and defence, make them a powerful system for addressing key questions in evolutionary biology in a comparative framework ^2,11,23,24^. However, a major roadblock to testing macro-evolutionary hypotheses is the lack of comprehensive quantitative datasets on venom diversity. Comparative phylogenetic analyses require thorough and balanced sampling across a phylogeny, but venom research has historically been biased towards medically significant species ^19^, resulting in datasets too limited and skewed to support robust inference ^25^.

Even for charismatic and medically important groups such as snakes, we still lack quantitative data from most species. Partly this is because the tools to generate quantitative proteomes have been lacking until the turn of the millennium. Over the last 20 years, modern proteomics and transcriptomics have transformed venomics, with robust workflows for identification and quantification ^26–29^. Most studies rely on bottom-up proteomics, where proteins are enzymatically digested into peptides that are separated and identified by Liquid Chromatography-Tandem Mass Spectrometry (LC–MS/MS), with peptide spectra matched to protein sequence databases ^30,31^. Relative toxin abundances can then be estimated using label-free quantification methods based on peptide signal intensities, allowing researchers to generate quantitative venom profiles without the need for expensive and labour intensive isotopic or chemical labelling. These approaches are increasingly becoming more accessible and high-throughput ^29^.

While the proteomic technology and analytical methods to boost our understanding of venom diversity are in place, the main limitation remains obtaining a comprehensive set of fresh samples. To date, venom studies have relied almost exclusively on live animals, but this imposes important logistical, legal, safety and ethical limitations. For clades that are species-rich, span multiple countries with differing regulations, inhabit remote and inaccessible habitats, or are secretive in nature, robust sampling could translate into lifetimes of fieldwork. Natural history collections offer a potential solution. Museums worldwide hold millions of specimens collected over more than two centuries, often representing entire clades within single or a handful of institutions. They are already invaluable for taxonomy, morphology, and ecology ^32^, and dissections of preserved specimens have yielded unparalleled contributions to our knowledge on diet and reproductive biology of some groups ^33^. Crucially, recent advances have demonstrated that genomic data, including whole genomes and even epigenomic signatures, can be recovered from formalin-fixed tissues, making it now possible to advance our understanding of phylogenetics, epigenetics and adaptation on even extinct populations ^34– 37^. To date there have been no studies attempting to extract biochemical data from the venom glands of preserved specimens, leaving a potentially game-changing resource untapped.

Here, we investigate the potential of natural history collections to study venom diversity using one of the most iconic and diverse venomous radiations: the Hydrophiinae elapid snakes. This relatively young clade colonised Australasia around 25 million years ago and has since diversified into one of the most remarkable vertebrate groups on Earth ^38–40^. They dominate the Australian snake fauna, ranging from conspicuous large-bodied predators such as taipans (*Oxyuranus*), black snakes (*Pseudechis*) and brown snakes (*Pseudonaja*), to fossorial specialists that feed exclusively on blind snakes (*Vermicella*) or reptile eggs (*Brachyurophis*). Most remarkably, the radiation produced the two most diverse marine reptile groups alive today, the sea kraits (*Laticauda*) and the true sea snakes (*Hydrophis* clade), with the latter displaying an explosive diversification since its origin around 7 million years ago ^40,41^.

Coupled with this ecological diversification, these front-fanged snakes have also evolved an extraordinary variety of toxins and venom phenotypes ^19,42–44^. Individual species can express more than 100 distinct toxins spanning up to 20 protein families ^31,45^. The two dominant toxin families found in the hydrophiine arsenal are three-fingered toxins (3FTx) and phospholipases A_2_ (PLA_2_). 3FTxs are proteins that mostly act as presynaptic neurotoxins, binding to acetylcholine receptors, blocking nerve-muscle communication and inducing a deadly paralysis ^43^. PLA_2_s are one of the most widely distributed toxin families in snakes, with independent recruitment events in elapids and vipers ^24^. They have evolved a striking functional diversity, ranging from myotoxins or cytotoxins that destroy tissue by dissolving cell membranes to pre-synaptic neurotoxins that prevent neurotransmitter release. Notably, hydrophiines have evolved multi-meric PLA_2_s that are among the most potent neurotoxins known and play an important role on the notorious lethality of taipans and brown snakes ^46^. A summary of the diversity and functions of snake toxin families is provided in Table S1.

Hydrophiines are an ideal system to test key hypothesis on the evolution of venoms and to discover toxinological diversity, but quantitative data remain sparse: relative toxin-family abundances are known for only 20 of the ∼203 species (<10%). In this study, we apply liquid chromatography– tandem mass spectrometry (LC–MS/MS) to test whether reliable quantitative venom proteomes can be obtained from formalin-fixed, ethanol-stored venom glands spanning six decades of museum collections. We assess whether specimen age influences data quality, and we combine our results with published datasets to demonstrate the power of museum venomics for investigating patterns of venom diversity. More broadly, our study establishes a proof of concept for museum proteomics, showing that preserved specimens can be leveraged to unlock biochemical traits across clades, time periods, and geographic ranges that would otherwise remain inaccessible.

## Results

### Preserved venom glands yield reliable proteomes

From 64 formalin-fixed venom glands spanning 37 hydrophiine species, five viperid species, and up to 57 years of storage, we generated quantitative proteomes by LC–MS/MS and compared them with lyophilised milked venoms from 25 matched species. As expected, milked venom samples from live animals contained a higher proportion of venom proteins than preserved gland samples, which contained a lot of structural and non-venom proteins (mean 93.5% vs 71.2%; Fig. S1). Visualisation of all samples and published proteomes as pie charts (Appendix 1) highlights extensive interspecific and intraspecific variation in venom profiles.

To quantify gland–venom similarity we applied three complementary metrics: linear-model R^2^, Spearman rank-correlation R^2^, and beta-regression R^2^. Preserved glands correlated strongly with milked venoms from the same species across all three metrics (Appendix 1): LM R^2^ = 0.92 (p < 0.005), Spearman R^2^ = 0.76 (p = 0.02), and GLM beta regression R^2^ = 0.314 (p < 0.005). These correlations were substantially higher than comparisons between congeneric species (LM R^2^ = 0.82, p = 0.02; Spearman R^2^ = 0.67, p = 0.02; GLM beta regression R^2^ = 0.23, p = 0.004) and much higher than between genera (LM R^2^ = 0.50, p = 0.4; Spearman R^2^ = 0.30, p = 0.3; GLM beta regression R^2^ = 0.14, p = 0.11). The comparison of the formalin-fixed gland and milked venom from a single captive *Pseudechis australis* give an indication of intra-individual variation using these methods and showed almost identical profiles (LM R^2^ = 1.0, p < 0.005; Spearman R^2^ = 0.94, p < 0.005; GLM beta regression R^2^ = 0.39, p < 0.005), dominated by PLA_2_ with minor contributions from 3FTx, LAAO, and NGF.

### Specimen age does not compromise venom data quality

Specimen age had no effect on the similarity between preserved glands and fresh venoms (Fig. 2). Across species-level comparisons, correlation strength showed no significant relationship with age using LM (R^2^ = 0.016, F_1_,_38_ = 0.62, p = 0.43), Spearman correlation (R^2^ = 0.09, F_1_,_38_ = 3.76, p = 0.06), or GLM beta regression (R^2^ = 0.10, F_1_,_38_ = 3.99, p = 0.053). Interestingly, both LM and beta regression showed a weak positive trend with age. Species-specific analyses also revealed no systematic relationship between age and gland–venom similarity (Fig. 2).

**Figure 1.**
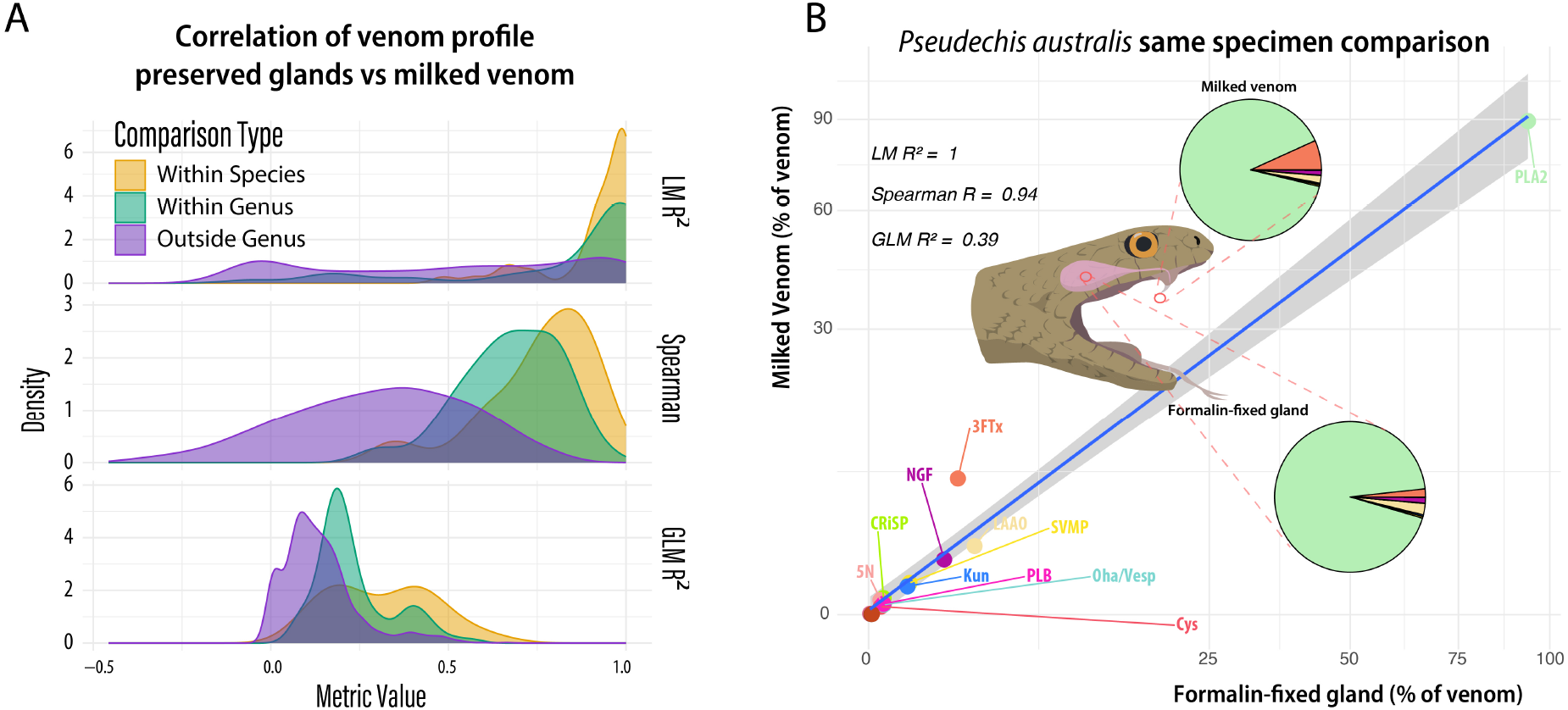
Preserved venom glands yield proteomes concordant with fresh milked venom. (A) Kernel density plots of R^2^ values from three metrics (linear model, Spearman rank correlation, and beta regression) comparing preserved glands to fresh venom at the same-species, same-genus, and different-genus levels. All three metrics show greater concordance within species. (B) Proteomic profiles from a formalin-fixed gland and fresh milked venom from the same individual mulga snake (*Pseudechis australis*) show near-identical composition, confirming that formalin fixation does not introduce systematic artifacts.

**Figure 2.**
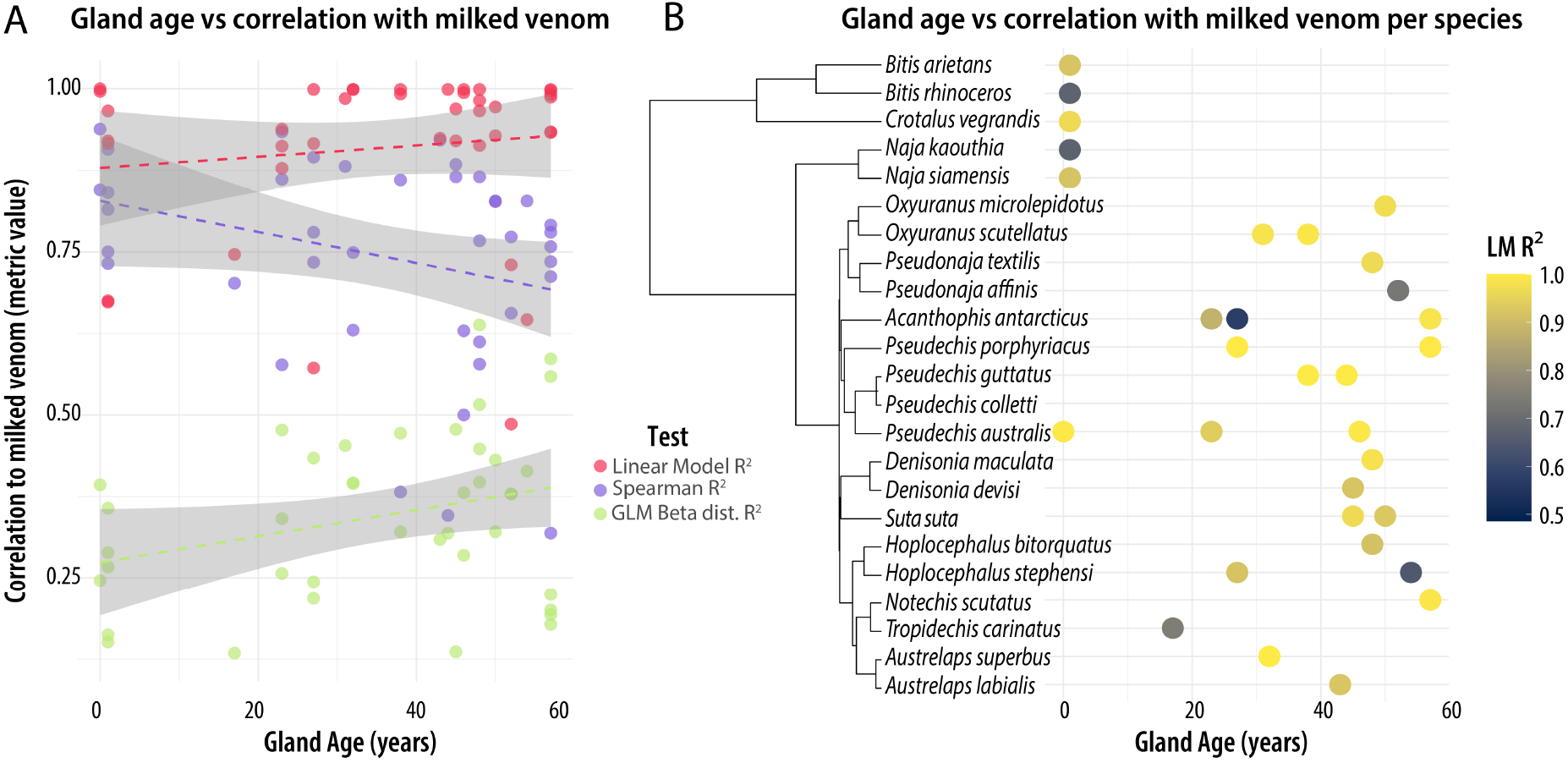
Specimen age does not affect proteomic data quality. (A) Regression of specimen age versus gland–venom correlation for each of the three metrics; none shows a significant relationship. (B) For each species, gland age on the x-axis and points coloured by linear model R^2^ of gland versus venom, illustrating that natural within-species variation exceeds any effect of preservation across specimens stored for more than half a century.

### Museum venomics extends known patterns of venom diversity

To place our preserved-gland data in the broader context of snake venom diversity, we combined our proteomes with an updated database of previously published snake proteomes covering Elapidae and most Viperidae. PCA of toxin-family abundances yielded consistent results with both raw and log(x+1)-transformed data and thus we only report the latter (Fig. 3). PC1 (48.9% of variance) was positively loaded on 3FTx (0.67) and negatively on SVMPs (– 0.462), SVSPs (–0.416), and CTLs (–0.28), clearly separating vipers (high SVMP/SVSP/CTL, little to no 3FTx) from elapids. PC2 (12.2%) was strongly associated with PLA_2_ (0.835, positive) and negatively with 3FTx (–0.327) and SVMP (–0.267), broadly separating hydrophiines (PLA_2_-rich) from Afro-Asian elapids (3FTx-rich), with sea kraits and sea snakes occupying intermediate space. However, Australo-Papuan elapids display a broad diversity with species displaying different phenotypes, ranging between PLA_2_ and 3FTx dominated venoms, including a variety of other toxin families such as Kunitz serine protease inhibitors, CTLs and CRiSPs. Both preserved gland and milked venom samples clustered with their respective clades, demonstrating that preserved material reliably matches known venom diversity patterns.

**Figure 3.**
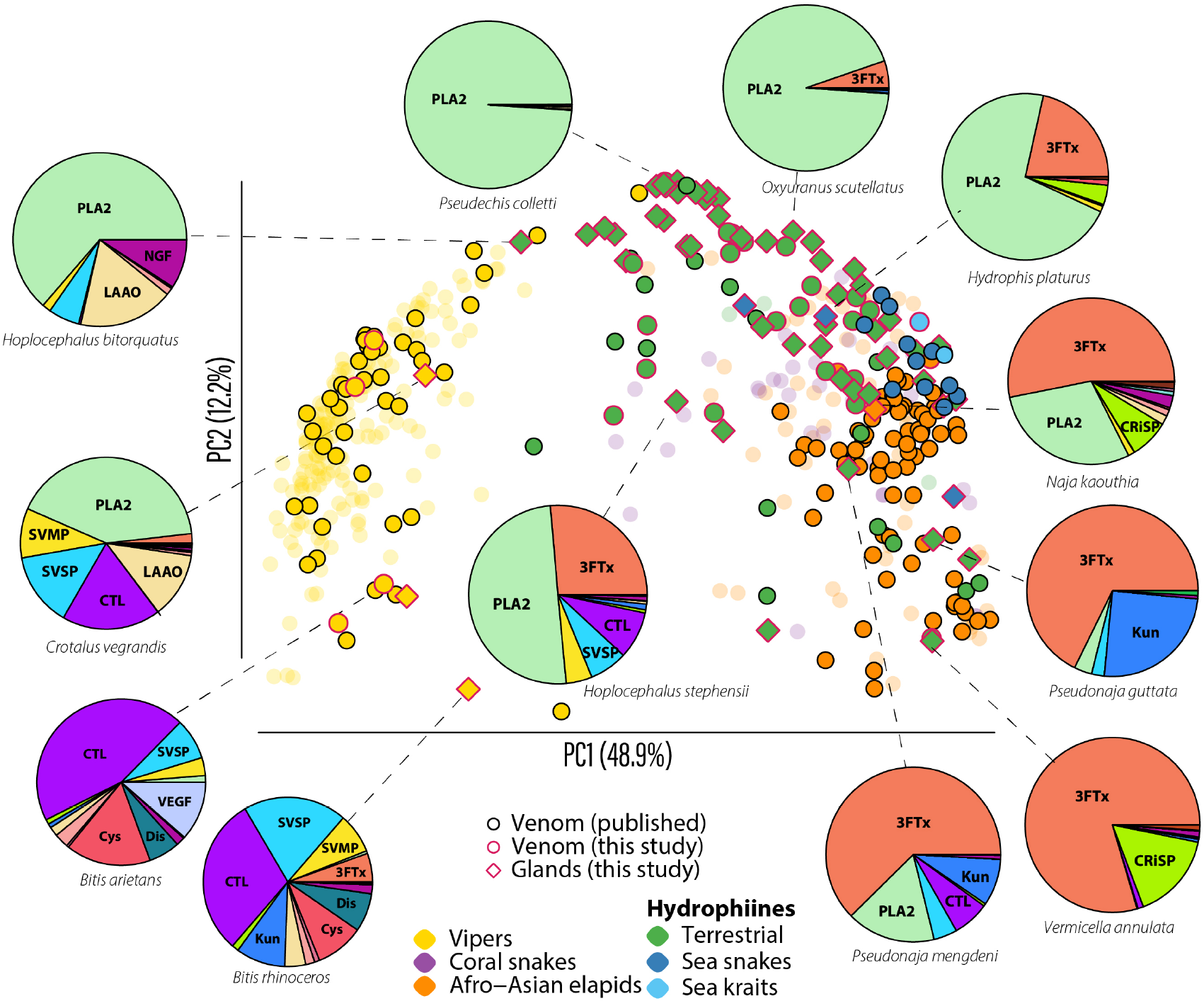
Museum venomics recovers known venom diversity patterns. Principal components analysis of log-transformed toxin-family abundances from this study (magenta outlines) and published proteomes. PC1 separates vipers (hemotoxin-dominated) from elapids (neurotoxin-dominated); PC2 separates PLA_2_-rich hydrophiines from 3FTx-rich Afro-Asian elapids. Preserved gland and milked venom samples cluster with their respective clades, confirming that museum material accurately recovers established diversity patterns. Faded points represent genera not included in our study.

## Discussion

Our study demonstrates that the millions of specimens stored in natural history collections across the globe offer a novel and unique opportunity to expand our knowledge on venom diversity. We show venom glands in preserved specimens yield equivalent proteomes to those from fresh milked venom, and importantly that the age of the specimen has no detectable effect on data quality. This opens a new incredibly rich source of biochemical data that can revolutionise how we study venom composition which can inform studies ranging from fundamental evolutionary biology to translational medicine and drug discovery.

### The potential of museum venomics

Natural history collections across the globe contain specimens collected over the last two centuries, which have only increased in scientific value as various human activities have caused global declines in biodiversity ^47^. Recent advances have shown that we can extract high quality genetic data from these specimens, including full organelle and nuclear genomes ^35,48–51^ and even epigenetic signatures from chromatin architecture ^35^. Proteomics has been applied to identify archaeological or palaeontological samples ^52–54^, or to analyse formalin-fixed patient tissue samples for biomedical research ^55^. Yet, proteomes have remained an overlooked dimension of diversity in natural history specimens ^56^. Our results provide evidence that venom proteomes can be recovered from museum specimens, yielding consistent toxin-family profiles even from glands fixed over half a century ago. The lack of an effect of age is especially encouraging, highlighting that proteins in venom remain detectable for as long as we tested. This is consistent with venoms being stored for decades retaining some biological activity ^57^. Importantly, for bottom-up proteomics, where proteins are digested into peptides, loss of tertiary structure and denaturation of proteins over time should not result in important changes in our ability to detect their presence in historic biological tissue. This age-independence significantly boosts the potential of historical specimens in proteomic studies.

Correlations between formalin-fixed glands and fresh venoms were remarkably high, with only a few exceptions best explained by natural biological variation rather than artefacts of preservation or methodology. Snake venoms are particularly variable phenotypes, particularly across geographic ranges and life-stages ^58–62^, but research into within population variation is conspicuously lacking. Importantly, identical proteomes within species should not be expected, thus divergent proteomes between some glands and venoms from different specimens are not concerning. In line with this, our samples of common death adders (*Acanthophis antarcticus*) from different localities contained the same dominant toxin families but in differing proportions (Appendix 1). Conversely, the comparison of preserved gland and fresh venom from the same *Pseudechis australis* individual produced nearly identical profiles, reinforcing the lack of artifacts introduced by formalin fixation and ethanol preservation.

We used three complementary statistical approaches to assess concordance between preserved glands and fresh venoms. Linear regressions (LM) quantify the strength of direct proportionality in toxin abundances, while Spearman rank correlations capture whether the relative ordering of toxin families is preserved even if their absolute values differ. Beta regressions reduce the influence of highly abundant toxins and give greater weight to low-abundance components. LM R^2^ values were consistently high, which is particularly relevant for evolutionary and ecological applications since dominant toxins are also likely to have the greatest functional importance ^24^. In contrast, beta regression R^2^ values are strongly shaped by small differences among minor toxins that can span orders of magnitude and are thus expected to be lower. Nevertheless, all three approaches converged on the same result: preserved glands correlate much more closely with fresh venoms of the same species than with those of congeners or unrelated taxa, underscoring the robustness of our findings.

We demonstrate how relatively easy it is to expand our knowledge on venom diversity using natural history collections. In a single LC–MS/MS experiment, we more than doubled the number of Australian snakes with quantitative venom profiles. Achieving this by relying on material from live specimens would have been extremely challenging. Finding all these species in the wild could take many years and tens of thousands of dollars as many of them are highly secretive, live in remote areas and many are amongst the most dangerous animals on Earth ^62– 65^. On the other hand, in a single museum collection a researcher could gather samples covering entire clades or continents in just a few days. With collaboration across major institutions, we envision the possibility of generating venom proteomes for most of the world’s venomous snake species within the next few years. Importantly, historical specimens also provide unique opportunities to study temporal dynamics, including decadal shifts in venom composition, akin to studies already conducted on other phenotypic dimensions ^32^.

Natural history collection specimens are a precious resource that often represents diversity that has since disappeared from nature, and destructive sampling should be carefully considered. However, destructive sampling does not necessarily result in total loss of the specimen and often only relies on taking a small sample, with precious information being obtained by doing so ^66^. Our approach requires removal of only one venom gland, leaving the skin, skeleton, and the second gland intact. Thus, specimens remain suitable for morphological studies, micro-CT scanning, and further venom analyses. This study adds to the expanding utility museum collections have for molecular research, and to the recognition that small, strategic destructive samples can yield disproportionately valuable data ^36,66^.

Comprehensive knowledge of venom diversity could have major life-saving translational benefits. Fatalities and permanent disabilities produced by snakebite have drastically reduced with access to anti-venoms in developed countries, but across many poorly resourced tropical regions snake envenomation remains a serious threat ^17^. Antibodies found in antivenoms are highly specific to the toxins they neutralise ^67,68^. While venoms are chemically diverse ^68^, the most serious pathological effects of venom are mediated by a handful of toxin families such as PLA2s, 3FTX, SVMPs and SVSPs ^69^. Since it is impractical and prohibitively costly to produce specific antivenoms for each of the hundreds of venomous snakes across the globe, one recent strategy for the development of next-generation snakebite treatments has been the development of broadly neutralizing antibodies focused on these major toxin families ^70–73^. An understanding of the full toxinological diversity of snakes is crucial for application of most suitable anti-venoms and antibodies.

### Patterns of snake venom diversity

When our data is integrated with published datasets, our results align with known venom diversity patterns across snakes. Vipers and elapids separate well, with the former relying heavily on hemotoxins such as SVMPs, SVSPs and CTLs and the latter leaning more towards neurotoxins like 3FTx and Kunitz peptides. Australo-Papuan and Afro-Asian elapids have broadly distinct venom strategies, relying mostly on PLA_2_s and 3FTxs, respectively. However, Australo-Papuan elapids have evolved a massive diversity of venom strategies that cover the whole spectrum, with species having PLA_2_s or 3FTxs dominated venoms and a wide diversity in between, highlighting that there are multiple effective cocktails that snakes can evolve to incapacitate prey. We confirm previous known patterns, like streamlined venoms of sea snakes and sea kraits featuring comparatively fewer toxin families ^74,75^, the notable reliance of Australian black snakes (*Pseudechis*) on PLA_2_^76,77^, the important fraction of SVMPs in *Hoplocephalus* ^78^, and the wildly diverse venoms of brown snakes (*Pseudonaja*) ^79^. Interestingly, our data also supports a previous study that found expression of 3FTx in African *Bitis* vipers ^80^, traditionally believed to be exclusive to elapid and colubrid snakes.

### Next steps and limitations

This approach is not limited to snakes or even venoms, as proteomes from most tissue types and taxa should be suitable for LC–MS/MS. However, challenges will arise when performing bottom-up proteomics in poorly characterised groups lacking thorough reference databases, where coupling museum proteomics with transcriptomics or top-down proteomics will be essential ^31,81^. This approach works well with snake venoms, because there have been enough genomic, transcriptomic, and proteomic studies to provide a robust reference database of the major toxin families which enabled us to characterize the specific composition of the samples in our studies. Nonetheless, exploration of snake venom diversity in lesser studied lineages would still be helpful by increasing the representation of rarer and novel toxins in the reference databases.

We also advise some degree of caution when comparing and integrating proteomic data across different studies, as there is a considerable variation in how proteins are identified, and especially how they are quantified ^26^. There is utility in interrogating these venom databases to establish broad patterns in venom evolution ^24,82^, but this can be much improved by using consistent pipelines. The taxonomic and geographic breadth museum venomics can leverage offers a promising pathway to have standardised and consistent datasets.

Quantitative proteomics is a fast-evolving field, and there are many recent technological advancements that can be incorporated to refine or expand museum proteomics. High-throughput venomics could upscale the volume of samples we can process ^29^, and multi-enzymatic limited digestion (rather than just trypsin digestion) of proteins can also increase the number of toxins we can identify ^83^. Absolute quantification and top-down venomics of intact toxins also provides potential for increased precision of identification and quantification of toxins found in venom glands ^26,84^. Finally, incorporating older specimens spanning the last two centuries could validate the exploration of proteomes across deeper time scales and profile venoms of long-gone populations and species.

We emphasise that museum proteomics complements, rather than replaces, studies of fresh venom, which remain critical for understanding gene expression, pharmacology, and toxicity. Nevertheless, further advances in museum genomics alongside refining museum proteomics could further advance how venoms and other proteomes are studied. Our results establish museum proteomics as a powerful new approach for evolutionary biology, ecology, and biomedicine. By unlocking the biochemical data preserved in collections worldwide, we can vastly expand the scale and scope of venom research and, more broadly, gain access to molecular phenotypes that would otherwise remain beyond reach.

## Methods

### Sample collection

We dissected one venom gland per specimen from 64 preserved snakes housed in the Australian National Wildlife Collection (CSIRO, Canberra), the Australian National University zoological collection and the South Australia Museum. These specimens represented 37 species in 16 genera of elapids, primarily from the Hydrophiinae, and five viperids (Table SX). Most had been initially fixed in formalin and subsequently stored in 70– 80% ethanol at room temperature for up to 57 years, as is typical of museum samples.

For comparison with preserved material, we acquired lyophilised milked venom from 25 of the same species through Venom Supplies Pty Ltd (Tanunda, South Australia). To directly test how preserved glands reflect fresh venom while controlling for geographic, and ontogenetic venom variation ^58,62^, we also milked an adult mulga snake (*Pseudechis australis*) from the Venom Supplies collection and then euthanised it two weeks later. Its venom glands were dissected, fixed in 10% formalin, and stored for three months in 80% ethanol, providing a same-individual validation when comparing its milked venom and preserved gland.

To contextualise our results, we assembled an updated database of published snake venom proteomes using ^19^ and ^13^ as starting points. This dataset incorporates all available Elapidae and most Viperidae proteomes published to date.

### Proteomics

Venom gland tissue samples were processed using a modified filter-aided sample preparation (FASP) protocol for formalin-fixed samples {Wiśniewski.2013gh}. Tissues preserved in ethanol were drained, weighed, and transferred to low protein-bind microcentrifuge tubes (≤100 mg tissue) containing a stainless-steel ball bearing, snap-frozen in liquid nitrogen, and homogenised in a Qiagen TissueLyzer (25 Hz, 2 min) with pre-chilled adapters. Samples were maintained frozen until lysis buffer (0.1 M Tris-HCl, pH 8.0, 0.1 M DTT, 2% SDS; prepared according to Sigma-Aldrich buffer guidelines) was added. Lysis buffer was added to ground venom gland samples at 10 µL/mg tissue. For lyophilised venom samples 250 µL of lysis buffer was added. Tubes were vortexed, heated at 99 °C with agitation (600 rpm, 1 h), and clarified by centrifugation (16,000 g, 18 °C, 10 min). Protein concentrations were determined using a reducing agent-compatible micro-BCA assay after dilution in 2% SDS.

For digestion, clarified lysates (50–100 µg protein) were combined with UA buffer (8 M urea, 0.1 M Tris-HCl, pH 8.5) containing 0.1 M DTT in 30 kDa MWCO ultrafiltration units, incubated at 37 °C for 30 min, and centrifuged (14,000 g) to <10 µL. Samples were sequentially washed with UA buffer, alkylated in 0.05 M iodoacetamide (IAA) in UA buffer (protected from light, 20 min), washed again with UA, then exchanged into digestion buffer (0.05 M Tris-HCl, pH 8.5). Proteins were digested with Trypsin/LysC (1:50 enzyme-to-protein ratio) in digestion buffer (40 µL) at 37 °C for 18 h in an Eppendorf ThermoMixer, followed by peptide recovery with additional digestion buffer and MilliQ water washes. Peptides were desalted using SDB-RPS STAGE tips with a Spin96 device as described in ^85^, dried, and resuspended in 0.1% formic acid for LC–MS/MS analysis.

Peptides equivalent to 1 µg of protein digest were analysed via data-dependent MS/MS on a Thermo Orbitrap Fusion ETD mass spectrometer coupled to a Dionex UltiMate 3000 RSLCnano LC system via a Nanospray Flex nano-ESI ion source (Thermo Part No. ES072). Samples were injected by the autosampler and initially trapped onto a PepMap100 C18 µ-Precolumn (5 µm particle size, 100 Å pore size; Thermo Scientific; Part No. 160454) with a 5 µL/min flow of 5% (v/v) acetonitrile in water for 3 min. The trap was then placed in-line with the analytical column, which was an in-house packed fused-silica capillary (∼20 cm × 75 µm ID × 360 µm OD; Polymicro Technologies; Part No. 1068150019) fitted with a chemical frit (Next Advance; Part No. FRIT-KIT) and packed with ReproSil-Pur 120 C18-AQ resin (1.9 µm; Dr Maisch GmbH; Part No. r119.aq.0001) using a Next Advance PC77 Pressure Injection Cell. The column temperature was maintained at 45 °C with an in-house fabricated oven ^86^.

Chromatographic separation was performed at 350 nL/min with the following gradient: 5% B (0–6 min), 12.5% B (9 min), 40% B (85 min), 80% B (86–99 min), 5% B (100–120 min); where Solvent A = water + 0.1% formic acid, and Solvent B = acetonitrile + 0.1% formic acid. Mass spectrometer settings were as follows: Application mode = Peptide; positive-ion mode ESI, static spray voltage 2.4 kV; ion transfer tube 305 °C; Orbitrap survey scans at 60,000 resolution, scan range 350–2000 m/z, RF lens = 60%, maximum injection time = 35 ms, intensity threshold = 5.0E3, charge states 2–6. Dynamic exclusion was 30 s (±10 ppm). MS/MS scans: isolation window 0.7 m/z, HCD collision energy = 30%, resolution = 15,000, maximum injection time = 22 ms. Duty cycle limited to 2 s, total method duration 120 min.

### Bioinformatics

Raw LC–MS/MS files were analysed in MaxQuant v2.0.3.0 (Max Planck Institute of Biochemistry) ^87^ using the Andromeda search engine against the UniProt database filtered by taxon=Serpentes (snakes). Searches assumed Trypsin/P specificity with up to two missed cleavages, carbamidomethylation of cysteine as a fixed modification, and oxidation of methionine and acetylation of protein N-termini as variable modifications. Precursor and fragment mass tolerances were set to 4.5 ppm and 20 ppm, respectively. Identifications were filtered at a 1% false discovery rate (FDR) at the peptide-spectrum match, protein, and site levels.

Protein quantification was performed using intensity-based absolute quantification (iBAQ) ^88^. This label-free method estimates protein abundance by dividing the summed peptide intensities by the number of theoretically observable peptides for each protein, providing relative measures that are comparable across proteins within a sample. Across all samples, >7,600 proteins were matched, of which 1,641 were classified as venom toxins, which is a consequence of using gland tissue. Each toxin was manually assigned to one of 24 protein families (Table SX2).

Data handling was performed using custom Python scripts to combine results into a single file. Subsequent analyses were conducted in R v4.4.2. Relative abundance of each toxin family in each sample was estimated as the sum of iBAQ values for all proteins in that family, divided by the sum of iBAQ values across all proteins identified in that sample.

To assess sample quality, we first compared the proportion of proteins known to occur in venoms across milked venom and preserved gland samples. We removed nine gland samples that had remarkably low amounts of venom proteins (under 15% of the total protein abundance) and were resulting in spurious profiles. For the remaining samples, all non-venom proteins were removed prior to downstream analyses.

### Statistical analyses

To evaluate how preserved venom gland samples compared to fresh venom, we performed pairwise statistical comparisons of toxin family proportions between each preserved gland sample and the corresponding milked venom from the same species. Because different correlation methods capture different aspects of similarity, we used three complementary approaches. First, we used a Linear regression (LM) to test for linear relationships in relative toxin abundances between glands and venoms. Second, we used a Spearman rank correlation to assess whether the rank order of toxin families was preserved, even if relative proportions differed. Third we used a Generalised linear model (GLM) with beta regression: to model proportional data on a log-odds scale, which reduces the influence of large values and gives more weight to minor toxin families. In summary, the LM evaluates abundance vs. scarcity, the Spearman correlation tests whether the relative order of toxin abundances is consistent, and the beta regression captures differences in order of magnitude across toxin families.

To verify that correlations were not simply due to general similarities among snake venoms, we also conducted control comparisons between different species within the same genus, and between species from different genera. We used the R functions *lm, cor*.*test* and *betareg* ^89^ to perform the LM, Spearman correlation and Beta Regression, respectively.

To test whether specimen age influenced the accuracy of preserved-gland profiles, we fitted linear models of gland–venom correlation (using all three metrics) against specimen age for each within-species comparison.

Finally, to visualise how well our data integrated with published venom diversity, we combined our dataset with the compiled venom proteome database. We performed a principal components analysis (PCA) both on raw proportions and on log(x+1)-transformed data to reduce the dominance of highly abundant toxin families, using the R function prcomp. We plotted samples coloured by major species groups: vipers (Viperidae), coral snakes (basal Elapidae: *Calliophis, Sinomicrurus, Micrurus*), Afro-Asian elapids (*Aspidelaps, Dendroaspis, Ophiophagus, Walterinnesia, Hemachatus, Naja, Bungarus*), sea kraits (*Laticauda*), sea snakes or marine hydrophiines (*Hydrophis, Aipysurus*), and terrestrial hydrophiines.

## Acknowledgements

Timothee Bonnet for advice on statistical analyses. Timothy Jackson, Bryan Fry and Theo Tasoulis for enlightening conversations when designing the project. We thank Leo Joseph, Tonya Haff and Chris Wilson for allowing access to the specimens under their care in the Australian National Wildlife Collection. DE thanks the Australian Research Council for the Discovery Early Career Research Award (DE230100003) and the Australian Academy of Sciences for the J.G. Russell Award for funding this research.

## Author contributions

Conceptualization: DE, JSK, ASM

Methodology: DE, JB, AC, DD, ND

Investigation: DE, JB, AC

Visualization: DE

Funding acquisition: DE, ASM

Project administration: DE

Supervision: DE, JSK, ASM

Writing – original draft: DE

Writing – review & editing: DE, JSK, DD, JB, ASM

## Data, code, and materials availability

All data and code are available on https://github.com/DEsquerre/MuseumVenomics.

