## Supplementary Materials for "Unlocking the venom vault: Museum venomics reveals an untapped biochemical archive in natural history collections"

---

**This file includes:**

Fig. S1
Table S1. Specimen information
Table S2. Toxin family descriptions
Appendix S1. Individual venom profiles (pie charts) — separate PDF file

### Figures

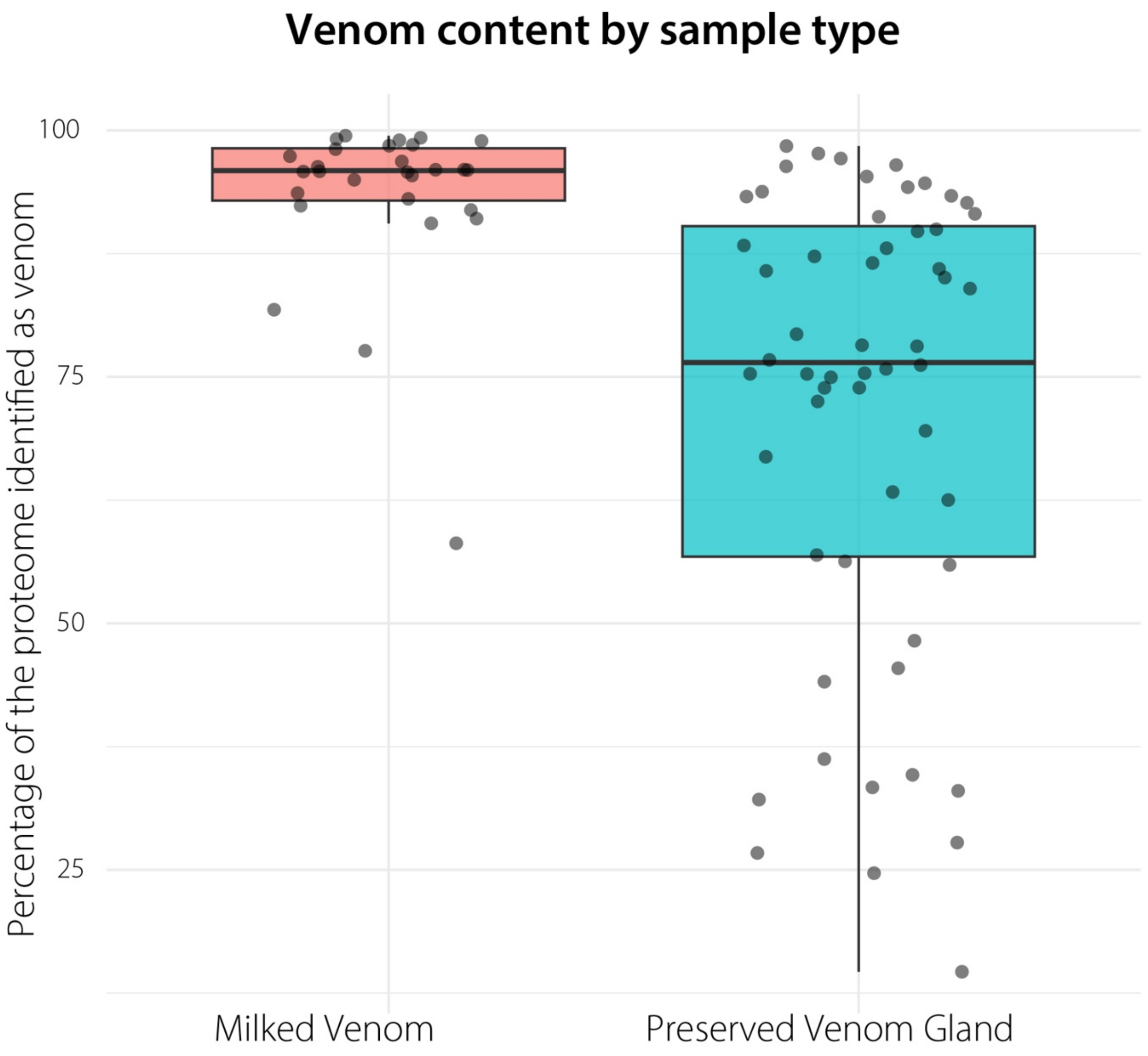


**Fig. S1.** Preserved glands contain a higher proportion of non-venom proteins than pure milked venom. Bar plot showing the percentage of total protein abundance attributable to proteins known to occur in venoms, in milked venom samples and preserved venom gland samples. Milked venoms showed a mean of 93.5% venom proteins vs 71.2% in preserved glands, reflecting the presence of structural and housekeeping proteins in gland tissue. Nine samples with <15% venom proteins were excluded from downstream analyses as outliers.

### Tables

**Table S2.** Summary of venom toxin protein families included in analyses, with abbreviations, nomenclature, role in venom composition, primary action, and brief description. Toxin families in the "Primary" category comprise the dominant components of venom in at least one species; "Minor" families are typically present at <5% total abundance.

| Abbreviation | Full name | Role in venom | Action | Description |
|---|---|---|---|---|
| 3FTx | Three-finger toxin | Primary | Neurotoxic, cytotoxic | Small non-enzymatic proteins common in elapid and colubrid venoms. Many are alpha-neurotoxins binding nicotinic acetylcholine receptors leading to paralysis. Some interfere with coagulation or platelet aggregation; others have cytotoxic effects by disrupting membranes via ion pores. |
| CRiSP | Cysteine-rich secretory protein | Primary | Ion channel blocking | Ancient reptile toxins that block calcium or potassium channels, affecting smooth muscle contraction, though most are uncharacterised. Found as minor components in front-fanged lineages but can make up more of the composition in other taxa. |
| CTL | C-type lectin | Primary | Coagulation modulation | Carbohydrate-binding lectins and snaclecs that bind platelet and coagulation factor targets, causing haemostatic disruption. Abundant in vipers, rare in elapids. |
| Kun | Kunitz-type protease inhibitor | Primary | Protease inhibition, ion channel blocking | Small peptides that either inhibit serine proteases (e.g., trypsin, chymotrypsin) to disrupt haemostasis or, in neurotoxic forms, block voltage-gated $K^+$ and $Ca^{2+}$ channels. Found mainly in elapids; dendrotoxins in mambas are examples. |
| LAAO | L-amino acid oxidase | Primary | Cytotoxic | Oxidatively deaminates L-amino acids, producing hydrogen peroxide, α-ketoacids, and ammonia; causes apoptosis, haemolysis, haemorrhage, and has antimicrobial effects. |
| PLA2 | Phospholipase A2 | Primary | Membrane disruption | Among the most widespread venom toxins. Hydrolyse phospholipids at the sn-2 position, generating lysophospholipids and fatty acids. Functions range from presynaptic neurotoxins (blocking neurotransmitter release) to myotoxins (inducing muscle necrosis) and anticoagulants. Exist in catalytic and non-catalytic forms, often oligomeric (e.g., taipoxin). |
| SVMP | Snake venom metalloproteinase | Primary | Haemorrhage, tissue damage | Zinc-dependent proteases that degrade extracellular matrix and coagulation factors. Major toxins in viper venoms causing haemorrhage, blistering, tissue necrosis, and consumptive coagulopathy. Categorised into PI–PIII classes. Present in smaller amounts in some elapids. |
| SVSP | Snake venom serine protease | Primary | Coagulation effects | Serine proteases that carry out a diverse range of coagulopathic functions by interacting with the kallikrein–kinin system or clotting cascade. Even procoagulant forms may lead to net anticoagulation when clots formed are unstable and quickly degraded (consumptive coagulopathy). |
| 5N | 5'-nucleotidase | Minor | Nucleotide breakdown | Widespread minor venom component that degrades nucleotides. Some evidence for inhibition of platelet aggregation. Through release of purines, may act as a spreading factor or hypotensive toxin. |
| AChE | Acetylcholinesterase | Minor | Neurotransmitter breakdown | Hydrolyses acetylcholine, disrupting nerve signal transmission at neuromuscular junctions; precise function in snake venoms is not entirely clear. |

| Abbreviation | Full name | Role in venom | Action | Description |
|---|---|---|---|---|
| AmPep | Aminopeptidase | Minor | Protein degradation | Exopeptidase that removes N-terminal amino acids from peptides and proteins, possibly aiding digestion of prey tissue. |
| Cys | Cystatin | Minor | Protease inhibition | Cysteine protease inhibitors commonly found as minor venom components, highly conserved. Unclear whether they play a role in venom preservation or as a supportive toxin. |
| Dis | Disintegrin | Minor | Anti-platelet | Small peptides inhibiting integrins, preventing platelet aggregation, cell adhesion, and migration; mostly found in viper venoms. |
| Hyal | Hyaluronidase | Minor | Tissue penetration | Degrades hyaluronic acid in connective tissue and interstitium, increasing spread of other venom components; can reduce swelling. |
| NGF | Nerve growth factor | Minor | Unclear | Common minor component of venoms; toxic activity remains unclear. May protect venom proteins from degradation by inhibiting metalloproteases or act as a spreading factor. |
| NP | Natriuretic peptide | Minor | Vasodilation | Hormones affecting the cardiovascular system; in venom, may lower blood pressure and affect prey circulation. |
| Oha-Vesp | Ohanin/Vespryn | Minor | Immobilisation | Non-enzymatic proteins that may impair prey mobility; minor component in some elapid venoms. |
| Other | Other toxins | Minor | Variable | Proteins not falling into main venom families or of unknown action. |
| PDE | Phosphodiesterase | Minor | Nucleic acid breakdown | Minor component of many snake venoms. Some evidence for DNA degradation and platelet inhibition; through release of purines, may act as a spreading factor or hypotensive toxin. |
| PLA2 inhibitor | Phospholipase A2 inhibitor | Minor | Blocks PLA2 activity | Endogenous snake proteins that bind venom PLA2s and prevent their activity. Likely evolved to protect snakes from autotoxic effects of their own venom. |
| PLB | Phospholipase B | Minor | Membrane degradation | Hydrolyses both sn-1 and sn-2 positions of phospholipids. Contributes to prey tissue degradation and enhances spread of other toxins. |
| PLC | Phospholipase C | Minor | Signal disruption | Hydrolyses phosphatidylinositol to generate second messengers, disrupting cell signalling. |
| Venom factor | Cobra venom factor | Minor | Complement activation | Complement-activating proteins found in cobras and close relatives. Forms stable convertases that hyperactivate and deplete host complement, suppressing immune response and promoting venom spread. |
| Venom peroxiredoxin | Venom peroxiredoxin | Minor | Antioxidant enzyme | Peroxide-reducing enzymes that detoxify reactive oxygen species. Role in venom remains unclear. |
| Waprin | Whey acidic protein motif protease inhibitor | Minor | Antimicrobial, protease inhibition | Small proteins containing a WAP domain. Possess antimicrobial activity against bacteria and fungi and can inhibit certain proteases. Role in venom remains unclear. |

**Table S1.** Specimen information for all samples included in this study. ANWC = Australian National Wildlife Collection (CSIRO, Canberra); ANU = Australian National University zoological collection. Preserved gland samples from ANWC and ANU were initially fixed in formalin and stored in 70–80% ethanol. SVL = snout–vent length. Genus and species names in italics.

| DEVC # | Specimen # | Origin | Sample type | Family | Genus | Species | Date collected | Country | State/Province | Locality | Sex | SVL | Age (yr) |
|---|---|---|---|---|---|---|---|---|---|---|---|---|---|
| DEVC 001 | PP043 | Venom Supplies | Milked venom | Elapidae | *Pseudechis* | *porphyriacus* | — | Australia | South Australia | Adelaide/Barossa region | — | — | — |
| DEVC 002 | AASA58 | Venom Supplies | Milked venom | Elapidae | *Acanthophis* | *antarcticus* | — | Australia | South Australia | South Australian locality | — | — | — |
| DEVC 003 | NSSA A0066 | Venom Supplies | Milked venom | Elapidae | *Notechis* | *scutatus* | — | Australia | South Australia | SE South Australia | — | — | — |
| DEVC 004 | — | ANU | Preserved gland | Elapidae | *Pseudechis* | *porphyriacus* | 1965 | Australia | No locality | — | — | — | — |
| DEVC 006 | — | ANU | Preserved gland | Elapidae | *Notechis* | *scutatus* | 1965 | Australia | No locality | — | — | — | — |
| DEVC 008 | — | ANU | Preserved gland | Elapidae | *Acanthophis* | *antarcticus* | 1965 | Australia | No locality | — | — | — | — |
| DEVC 010 | — | ANU | Preserved gland | Elapidae | *Pseudechis* | *porphyriacus* | 1965 | Australia | No locality | — | — | — | — |
| DEVC 012 | — | ANU | Preserved gland | Elapidae | *Notechis* | *scutatus* | 1965 | Australia | No locality | — | — | — | — |
| DEVC 014 | — | ANU | Preserved gland | Elapidae | *Acanthophis* | *antarcticus* | 1965 | Australia | No locality | — | — | — | — |
| DEVC 017 | BA1 | Venom Supplies | Gland (frozen/EtOH, no formalin) | Viperidae | *Bitis* | *arietans* | — | Captive bred | — | Zoo stock | Male | 1080 mm | — |
| DEVC 018 | MJF001 | Venom Supplies | Gland (frozen/EtOH, no formalin) | Elapidae | *Naja* | *siamensis* | — | Captive bred | — | Zoo stock | Female | 1430 mm | — |
| DEVC 019 | NK70 | Venom Supplies | Gland (frozen/EtOH, no formalin) | Elapidae | *Naja* | *kaouthia* | — | Captive bred | — | Zoo stock | Male | 882 mm | — |
| DEVC 020 | — | Venom Supplies | Gland (frozen/EtOH, no formalin) | Viperidae | *Bitis* | *rhinoceros* | — | Captive bred | — | Zoo stock | — | 919 mm | — |
| DEVC 021 | CV1 | Venom Supplies | Gland (frozen/EtOH, no formalin) | Viperidae | *Crotalus* | *vegrandis* | — | Captive bred | — | Zoo stock | Male | 935 mm | — |
| DEVC 022 | — | Venom Supplies | Milked venom | Elapidae | *Laticauda* | *colubrina* | — | Indonesia | Bali | — | — | — | — |
| DEVC 023 | — | Venom Supplies | Milked venom | Elapidae | *Hoplocephalus* | *stephensi* | — | Australia | Queensland | — | — | — | — |
| DEVC 024 | — | Venom Supplies | Milked venom | Viperidae | *Agkistrodon* | *bilineatus* | — | Captive bred | — | Zoo stock | — | — | — |
| DEVC 025 | — | Venom Supplies | Milked venom | Viperidae | *Bitis* | *rhinoceros* | — | Captive bred | — | Zoo stock | — | — | — |
| DEVC 026 | — | Venom Supplies | Milked venom | Elapidae | *Naja* | *kaouthia* | — | Captive bred | — | Zoo stock | — | — | — |
| DEVC 027 | — | Venom Supplies | Milked venom | Elapidae | *Pseudechis* | *guttatus* | — | Australia | Queensland | — | — | — | — |

| DEVC # | Specimen # | Origin | Sample type | Family | Genus | Species | Date collected | Country | State/Province | Locality | Sex | SVL | Age (yr) |
|---|---|---|---|---|---|---|---|---|---|---|---|---|---|
| DEVC 028 | — | Venom Supplies | Milked venom | Viperidae | *Bitis* | *arietans* | — | Captive bred | — | Zoo stock | — | — | — |
| DEVC 029 | — | Venom Supplies | Milked venom | Elapidae | *Austrelaps* | *superbus* | — | Australia | South Australia | — | — | — | — |
| DEVC 030 | — | Venom Supplies | Milked venom | Elapidae | *Pseudonaja* | *inframacula* | — | Australia | South Australia | — | — | — | — |
| DEVC 031 | — | Venom Supplies | Milked venom | Elapidae | *Oxyuranus* | *scutellatus* | — | Australia | Queensland | — | — | — | — |
| DEVC 032 | CV0003 | Venom Supplies | Milked venom | Viperidae | *Crotalus* | *vegrandis* | — | Captive bred | — | Zoo stock | — | — | — |
| DEVC 033 | — | Venom Supplies | Milked venom | Elapidae | *Pseudonaja* | *affinis* | — | Australia | Western Australia | — | — | — | — |
| DEVC 034 | — | Venom Supplies | Milked venom | Elapidae | *Pseudechis* | *colletti* | — | Australia | Queensland | — | — | — | — |
| DEVC 035 | CB492 | Venom Supplies | Milked venom | Elapidae | *Pseudonaja* | *textilis* | — | Australia | South Australia | — | — | — | — |
| DEVC 036 | — | Venom Supplies | Milked venom | Elapidae | *Hoplocephalus* | *bitorquatus* | — | Australia | Queensland | — | — | — | — |
| DEVC 037 | — | Venom Supplies | Milked venom | Elapidae | *Echiopsis* | *curta* | — | Australia | Western Australia | — | — | — | — |
| DEVC 038 | — | Venom Supplies | Milked venom | Elapidae | *Pseudechis* | *australis* | — | Australia | Western Australia | — | — | — | — |
| DEVC 039 | — | Venom Supplies | Milked venom | Elapidae | *Oxyuranus* | *microlepidotus* | — | Australia | South Australia | — | — | — | — |
| DEVC 040 | — | Venom Supplies | Milked venom | Elapidae | *Naja* | *siamensis* | — | Captive bred | — | Zoo stock | — | — | — |
| DEVC 041 | — | Venom Supplies | Milked venom | Elapidae | *Denisonia* | *devisi* | — | Australia | Queensland | — | — | — | — |
| DEVC 042 | — | Venom Supplies | Milked venom | Elapidae | *Austrelaps* | *labialis* | — | Australia | Queensland | — | — | — | — |
| DEVC 043 | — | Venom Supplies | Milked venom | Elapidae | *Tropidechis* | *carinatus* | — | Captive bred | — | Zoo stock | — | — | — |
| DEVC 044 | — | Venom Supplies | Milked venom | Elapidae | *Denisonia* | *maculata* | — | Australia | Queensland | — | — | — | — |
| DEVC 045 | — | Venom Supplies | Milked venom | Elapidae | *Suta* | *suta* | — | Australia | Queensland | — | — | — | — |
| DEVC 046 | R02840 | ANWC | Preserved gland | Elapidae | *Pseudonaja* | *textilis* | 1974-11-12 | Australia | ACT | Aranda, Canberra | Female | 1280 mm | — |
| DEVC 050 | R00152 | ANWC | Preserved gland | Elapidae | *Pseudonaja* | *affinis* | 1970-10-31 | Australia | Western Australia | 9 Miles E Of N Bannister | — | 1640 mm | — |
| DEVC 052 | R01968 | ANWC | Preserved gland | Elapidae | *Pseudonaja* | *affinis* | 1970-08-01 | Australia | South Australia | 25 Miles E Of Ceduna | — | head & tail | — |
| DEVC 056 | R00801 | ANWC | Preserved gland | Elapidae | *Pseudonaja* | *guttatus* | 1972-11-01 | Australia | Queensland | Lorna Downs Station | — | 930 mm | — |
| DEVC 058 | R03095 | ANWC | Preserved gland | Elapidae | *Pseudonaja* | *mengdeni* | — | Australia | South Australia | Moolawatana Homestead | Male | 1100 mm | — |
| DEVC 060 | R03096 | ANWC | Preserved gland | Elapidae | *Pseudonaja* | *mengdeni* | — | Australia | South Australia | 20 km E Of Erudina Station | Male | 1090 mm | — |

| DEVC # | Specimen # | Origin | Sample type | Family | Genus | Species | Date collected | Country | State/Province | Locality | Sex | SVL | Age (yr) |
|---|---|---|---|---|---|---|---|---|---|---|---|---|---|
| DEVC 062 | R06089 | ANWC | Preserved gland | Elapidae | *Acanthophis* | *antarcticus* | 1999-02-10 | Australia | NSW | Wapengo | Female | 850 mm | — |
| DEVC 064 | R05679 | ANWC | Preserved gland | Elapidae | *Acanthophis* | *antarcticus* | 1995-01-17 | Australia | NSW | Murwillumbah/Kyogle Road | — | 500 mm | — |
| DEVC 066 | R10857 | ANWC | Preserved gland | Elapidae | *Acanthophis* | *laevis* | 2015-04-06 | Papua New Guinea | Western Province | Morehead | — | 550 mm | — |
| DEVC 068 | R01054 | ANWC | Preserved gland | Elapidae | *Acanthophis* | *laevis* | 1971-07-14 | Indonesia | West Papua | Cape Steenboom | — | 515 mm | — |
| DEVC 070 | R05272 | ANWC | Preserved gland | Elapidae | *Acanthophis* | *praelongus* | 1990-08-13 | Australia | Queensland | Cape York Peninsula | — | 360 mm | — |
| DEVC 072 | R3016 | ANWC | Preserved gland | Elapidae | *Aipysurus* | *laevis* | 1979-12-17 | Australia | Coral Sea Islands | Saumarez Reef | — | 940 mm | — |
| DEVC 075 | R03110 | ANWC | Preserved gland | Elapidae | *Austrelaps* | *labialis* | — | Australia | South Australia | Kangaroo Island | Female | 570 mm | — |
| DEVC 077 | R02819 | ANWC | Preserved gland | Elapidae | *Austrelaps* | *labialis* | 1979-05-01 | Australia | South Australia | Kangaroo Island | Male | 620 mm | — |
| DEVC 079 | R01162 | ANWC | Preserved gland | Elapidae | *Austrelaps* | *ramsayi* | 1976-01-18 | Australia | NSW | Peppers Ck, Numeralla | — | 810 mm | — |
| DEVC 081 | R05174 | ANWC | Preserved gland | Elapidae | *Austrelaps* | *ramsayi* | 1989-12-07 | Australia | Tasmania | Flinders Island | — | 1070 mm | — |
| DEVC 083 | R05179 | ANWC | Preserved gland | Elapidae | *Austrelaps* | *superbus* | 1990-02-26 | Australia | Tasmania | Flinders Island | — | 970 mm | — |
| DEVC 085 | R05508 | ANWC | Preserved gland | Elapidae | *Austrelaps* | *superbus* | 1990-03-02 | Australia | Tasmania | Flinders Island | — | 1000 mm | — |
| DEVC 089 | R01674 | ANWC | Preserved gland | Elapidae | *Denisonia* | *devisi* | 1977-08-11 | Australia | NSW | Macquarie Marshes | — | 440 mm | — |
| DEVC 091 | R01427 | ANWC | Preserved gland | Elapidae | *Denisonia* | *maculata* | 1974-01-01 | Australia | Queensland | Yeppen, S Of Rockhampton | — | 340 mm | — |
| DEVC 093 | R02710 | ANWC | Preserved gland | Elapidae | *Drysdalia* | *coronoides* | 1968-12-07 | Australia | NSW | Mt Gingera, Brindabella Range | — | 345 mm | — |
| DEVC 099 | R02027 | ANWC | Preserved gland | Elapidae | *Hoplocephalus* | *bitorquatus* | 1974-01-01 | Australia | Queensland | Rockhampton | — | 450 mm | — |
| DEVC 101 | R02882 | ANWC | Preserved gland | Elapidae | *Hoplocephalus* | *bitorquatus* | 1974-02-01 | Australia | Queensland | Yeppen Yeppen Lagoon | — | 520 mm | — |
| DEVC 103 | R00147 | ANWC | Preserved gland | Elapidae | *Hoplocephalus* | *stephensi* | 1968-02-11 | Australia | Queensland | Brisbane | — | 945 mm | — |
| DEVC 105 | R05736 | ANWC | Preserved gland | Elapidae | *Hoplocephalus* | *stephensi* | 1995-01-26 | Australia | NSW | Whian Whian State Forest | — | 585 mm | — |
| DEVC 107 | R05923 | ANWC | Preserved gland | Elapidae | *Hydrophis* | *platurus* | 1997-01-12 | Australia | NSW | Bengello Beach, Moruya | — | 650 mm | — |
| DEVC 109 | R05592 | ANWC | Preserved gland | Elapidae | *Hydrophis* | *platurus* | 1992 | Australia | ACT | Jervis Bay | — | 931 mm | — |
| DEVC 111 | R00581 | ANWC | Preserved gland | Elapidae | *Oxyuranus* | *microlepidotus* | 1972-08-03 | Australia | Queensland | Lorna Downs Homestead | Male | 1490 mm | — |
| DEVC 113 | R04739 | ANWC | Preserved gland | Elapidae | *Oxyuranus* | *scutellatus* | 1984-08-01 | Australia | Queensland | Gordonvale | — | 1950 mm | — |
| DEVC 115 | R05467 | ANWC | Preserved gland | Elapidae | *Oxyuranus* | *scutellatus* | 1991-08-28 | Australia | Queensland | Shoalwater Bay | Male | 1840 mm | — |

| DEVC # | Specimen # | Origin | Sample type | Family | Genus | Species | Date collected | Country | State/Province | Locality | Sex | SVL | Age (yr) |
|---|---|---|---|---|---|---|---|---|---|---|---|---|---|
| DEVC 117 | R01201 | ANWC | Preserved gland | Elapidae | *Pseudechis* | *australis* | 1976-01-27 | Australia | Northern Territory | Mcarthur River | Male | 2000 mm | — |
| DEVC 119 | R06173 | ANWC | Preserved gland | Elapidae | *Pseudechis* | *australis* | 1999-11-12 | Australia | Queensland | Thargomindah–Cunnamulla | Male | 1890 mm | — |
| DEVC 121 | R02070 | ANWC | Preserved gland | Elapidae | *Pseudechis* | *colletti* | — | Australia | Queensland | Hughenden | — | 610 mm | — |
| DEVC 123 | R10999 | ANWC | Preserved gland | Elapidae | *Pseudechis* | *rossignolii* | 2015-03-26 | Papua New Guinea | — | Morehead–Mibini Road | — | 380 mm | — |
| DEVC 125 | R01899 | ANWC | Preserved gland | Elapidae | *Pseudechis* | *guttatus* | 1978-02-22 | Australia | NSW | Teasdale Dam, NE Of Warren | — | 1010 mm | — |
| DEVC 127 | R04630 | ANWC | Preserved gland | Elapidae | *Pseudechis* | *guttatus* | 1984-10-14 | Australia | NSW | 23 Km E Of Caragabal | — | 1015 mm | — |
| DEVC 129 | R05784 | ANWC | Preserved gland | Elapidae | *Pseudechis* | *porphyriacus* | 1995-04-29 | Australia | NSW | Nullum State Forest, Tweed Valley | — | 1380 mm | — |
| DEVC 131 | R01521 | ANWC | Preserved gland | Elapidae | *Suta* | *suta* | 1977-02-09 | Australia | NSW | Woorandara Station, W Of Booligal | — | 370 mm | — |
| DEVC 133 | R00920 | ANWC | Preserved gland | Elapidae | *Suta* | *suta* | 1972-07-28 | Australia | Northern Territory | Brunette Downs, Barkly Tablelands | — | 610 mm | — |
| DEVC 135 | R06918 | ANWC | Preserved gland | Elapidae | *Tropidechis* | *carinatus* | 2005-11-17 | Australia | Queensland | Mt Lewis Via Julatten | — | 950 mm | — |
| DEVC 137 | R03109 | ANWC | Preserved gland | Elapidae | *Tropidechis* | *carinatus* | — | Australia | Queensland | Springbrook | Male | 680 mm | — |
| DEVC 139 | R06769 | ANWC | Preserved gland | Elapidae | *Vermicella* | *annulata* | 2002-11-09 | Australia | Queensland | SE Of Maleny | — | 680 mm | — |
| DEVC 141 | R06675 | ANWC | Preserved gland | Elapidae | *Vermicella* | *annulata* | 2000-12-24 | Australia | Queensland | NW Of Maleny | — | 600 mm | — |
| DEVC 143 | R00732 | ANWC | Preserved gland | Elapidae | *Vermicella* | *multifasciata* | 1972-03-04 | Australia | Northern Territory | Adelaide River | — | 485 mm | — |
| DEVC 145 | PA132 | Venom Supplies | Milked venom (same-individual validation) | Elapidae | *Pseudechis* | *australis* | 2022-09-01 | Australia | Captive bred | Venom Supplies | — | — | — |
| DEVC 146 | PA134 | Venom Supplies | Preserved gland (same-individual validation) | Elapidae | *Pseudechis* | *australis* | 2022-09-01 | Australia | Captive bred | Venom Supplies | — | — | — |

### Appendix S1

Individual venom proteomic profiles for all samples included in this study and in the published venom proteome database, shown as pie charts with toxin family proportions. Samples are ordered alphabetically by genus and species name. Each pie chart is labelled with the DEVC collection number (for museum and validation samples) or source reference (for published proteomes), and the year of collection where known. Pie chart colour scheme follows the toxin family abbreviations in Table S2.

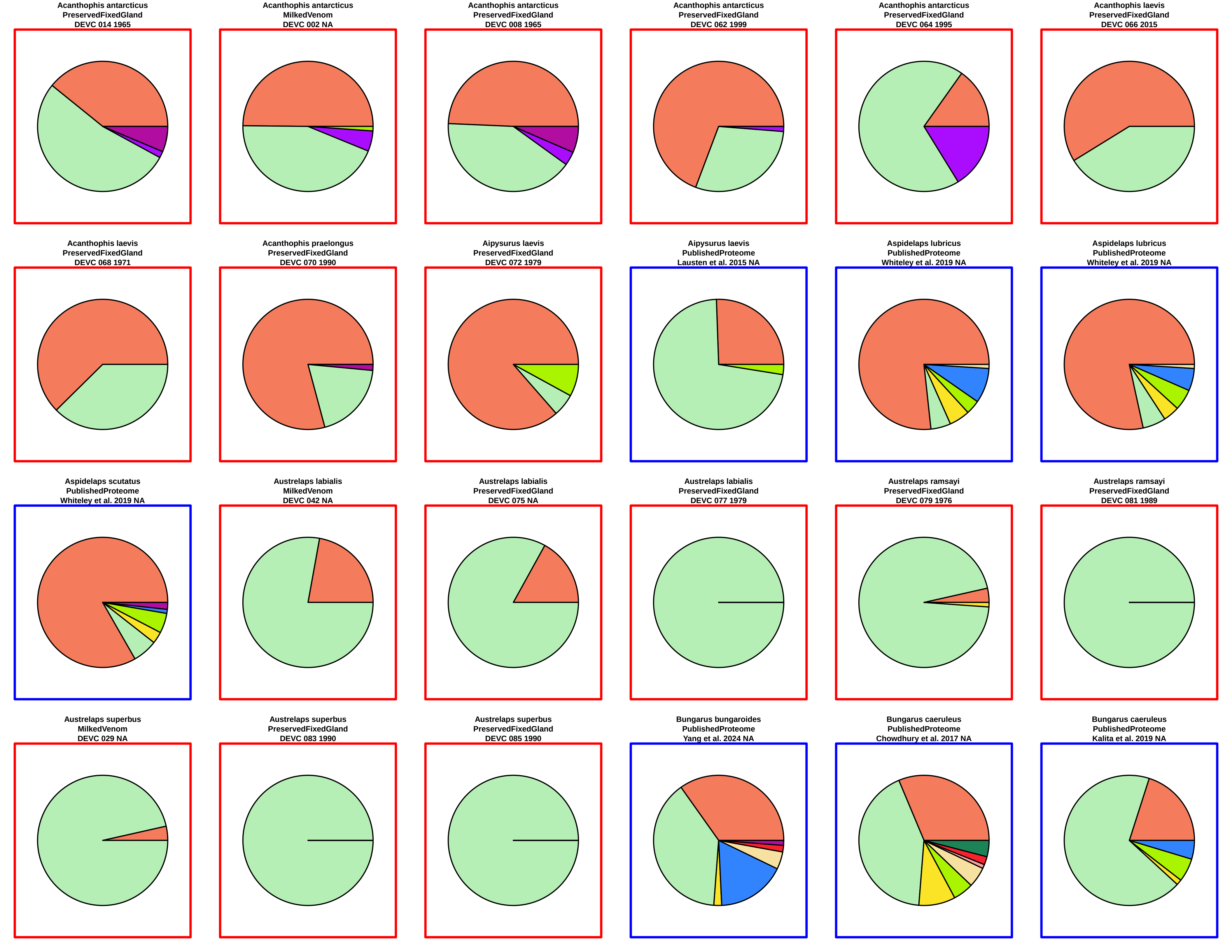
Acanthophis antarcticus
PreservedFixedGland
DEVC 014 1965
Acanthophis antarcticus
MilkedVenom
DEVC 002 NA
Acanthophis antarcticus
PreservedFixedGland
DEVC 008 1965
Acanthophis antarcticus
PreservedFixedGland
DEVC 062 1999
Acanthophis antarcticus
PreservedFixedGland
DEVC 064 1995
Acanthophis laevis
PreservedFixedGland
DEVC 066 2015
Acanthophis laevis
PreservedFixedGland
DEVC 068 1971
Acanthophis praelongus
PreservedFixedGland
DEVC 070 1990
Aipysurus laevis
PreservedFixedGland
DEVC 072 1979
Aipysurus laevis
PublishedProteome
Lausten et al. 2015 NA
Aspidelaps lubricus
PublishedProteome
Whiteley et al. 2019 NA
Aspidelaps lubricus
PublishedProteome
Whiteley et al. 2019 NA
Aspidelaps scutatus
PublishedProteome
Whiteley et al. 2019 NA
Austrelaps labialis
MilkedVenom
DEVC 042 NA
Austrelaps labialis
PreservedFixedGland
DEVC 075 NA
Austrelaps labialis
PreservedFixedGland
DEVC 077 1979
Austrelaps ramsayi
PreservedFixedGland
DEVC 079 1976
Austrelaps ramsayi
PreservedFixedGland
DEVC 081 1989
Austrelaps superbus
MilkedVenom
DEVC 029 NA
Austrelaps superbus
PreservedFixedGland
DEVC 083 1990
Austrelaps superbus
PreservedFixedGland
DEVC 085 1990
Bungarus bungaroides
PublishedProteome
Yang et al. 2024 NA
Bungarus caeruleus
PublishedProteome
Chowdhury et al. 2017 NA
Bungarus caeruleus
PublishedProteome
Kalita et al. 2019 NA

Bungarus caeruleus
PublishedProteome
Patra et al. 2019 NA

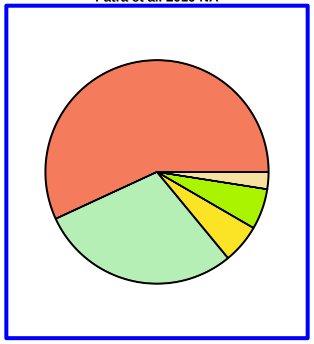

Bungarus caeruleus
PublishedProteome
Rashmi et al. 2024 NA

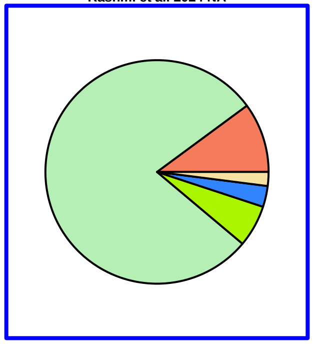

Bungarus caeruleus
PublishedProteome
Rashmi et al. 2024 NA

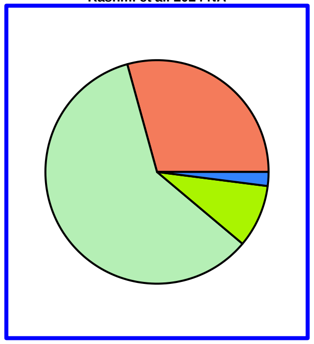

Bungarus caeruleus
PublishedProteome
Rashmi et al. 2024 NA

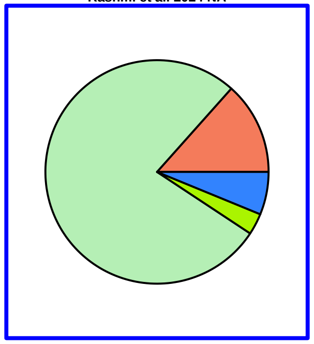

Bungarus caeruleus
PublishedProteome
Rashmi et al. 2024 NA

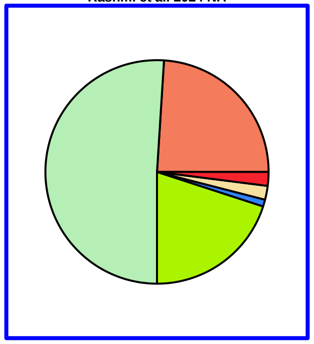

Bungarus caeruleus
PublishedProteome
Rashmi et al. 2024 NA

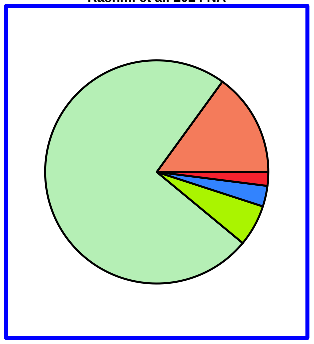

Bungarus caeruleus
PublishedProteome
Senji Laxme et al. 2019 NA

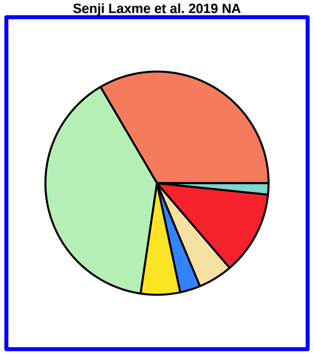

Bungarus caeruleus
PublishedProteome
Sunagar et al. 2021 NA

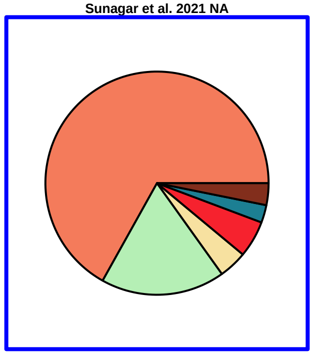

Bungarus caeruleus
PublishedProteome
Sunagar et al. 2021 NA

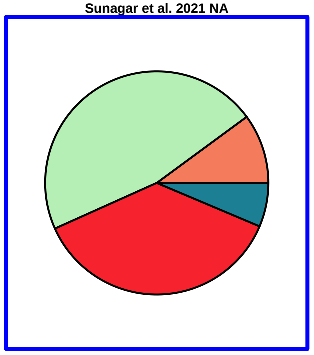

Bungarus candidus
PublishedProteome
Rusmili et al. 2014 NA

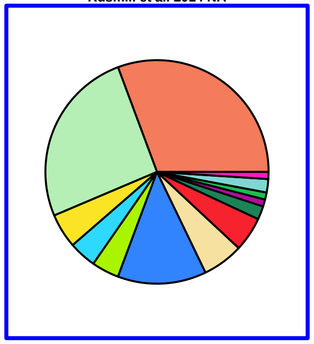

Bungarus fasciatus
PublishedProteome
Hia et al. 2020 NA

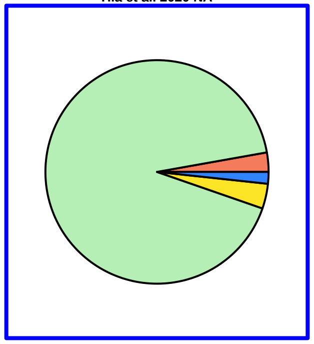

Bungarus fasciatus
PublishedProteome
Hia et al. 2020 NA

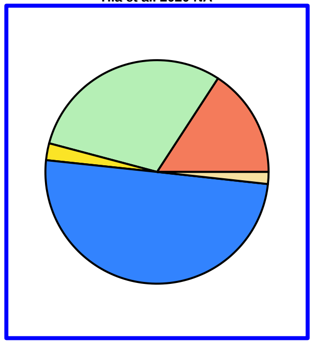

Bungarus fasciatus
PublishedProteome
Hia et al. 2020 NA

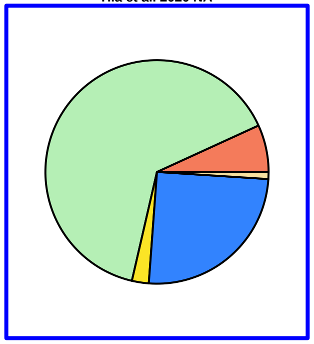

Bungarus fasciatus
PublishedProteome
Hia et al. 2020 NA

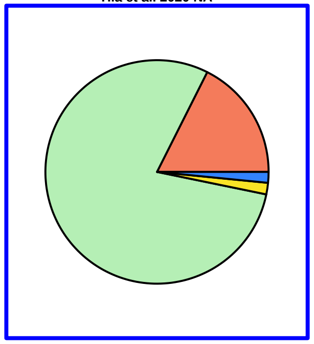

Bungarus fasciatus
PublishedProteome
Hia et al. 2020 NA

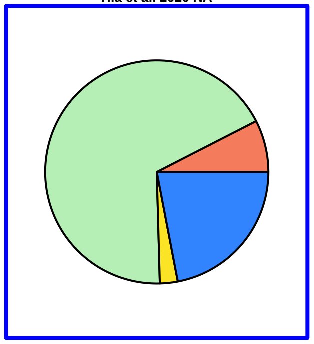

Bungarus fasciatus
PublishedProteome
Rusmili et al. 2016 NA

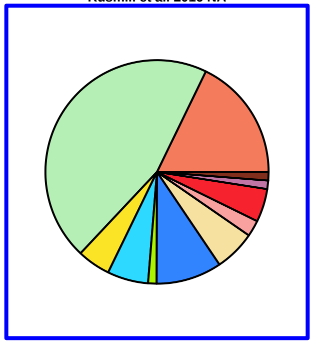

Bungarus fasciatus
PublishedProteome
Senji Laxme et al. 2019 NA

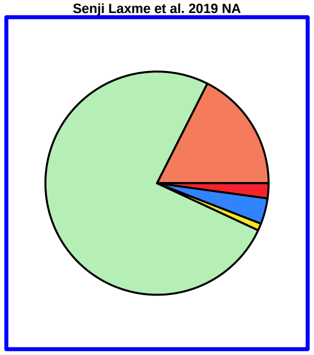

Bungarus fasciatus
PublishedProteome
Ziganshin et al. 2015 NA

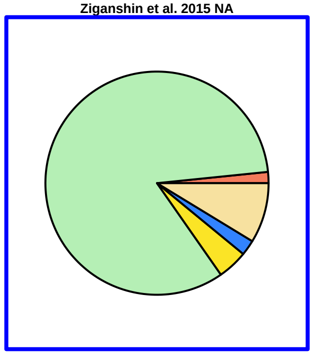

Bungarus flaviceps
PublishedProteome
Chapeaurouge et al. 2018. NA

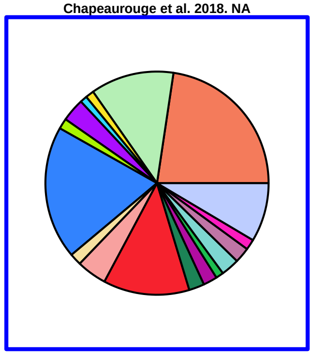

Bungarus flaviceps
PublishedProteome
Tan et al. 2022 NA

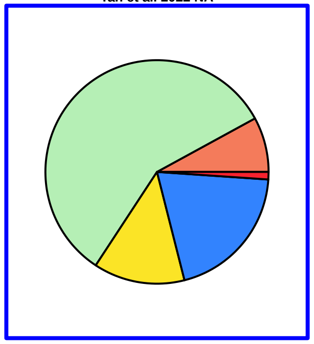

Bungarus multicinctus
PublishedProteome
Oh et al. 2021 NA

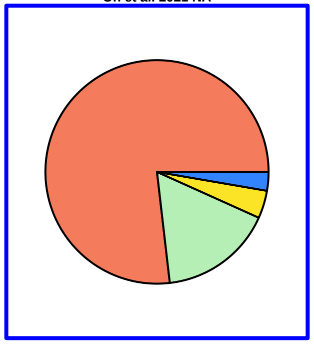

Bungarus multicinctus
PublishedProteome
Oh et al. 2021 NA

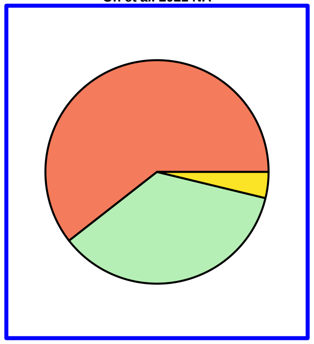

Bungarus multicinctus
PublishedProteome
Qin et al. 2023 NA

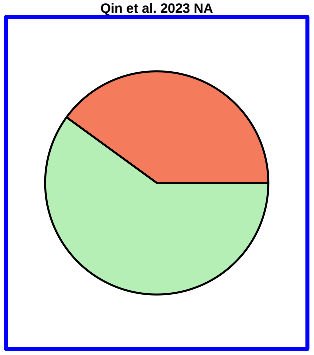

Bungarus multicinctus
PublishedProteome
Shan et al. 2016 NA

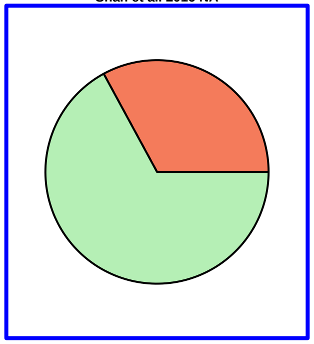

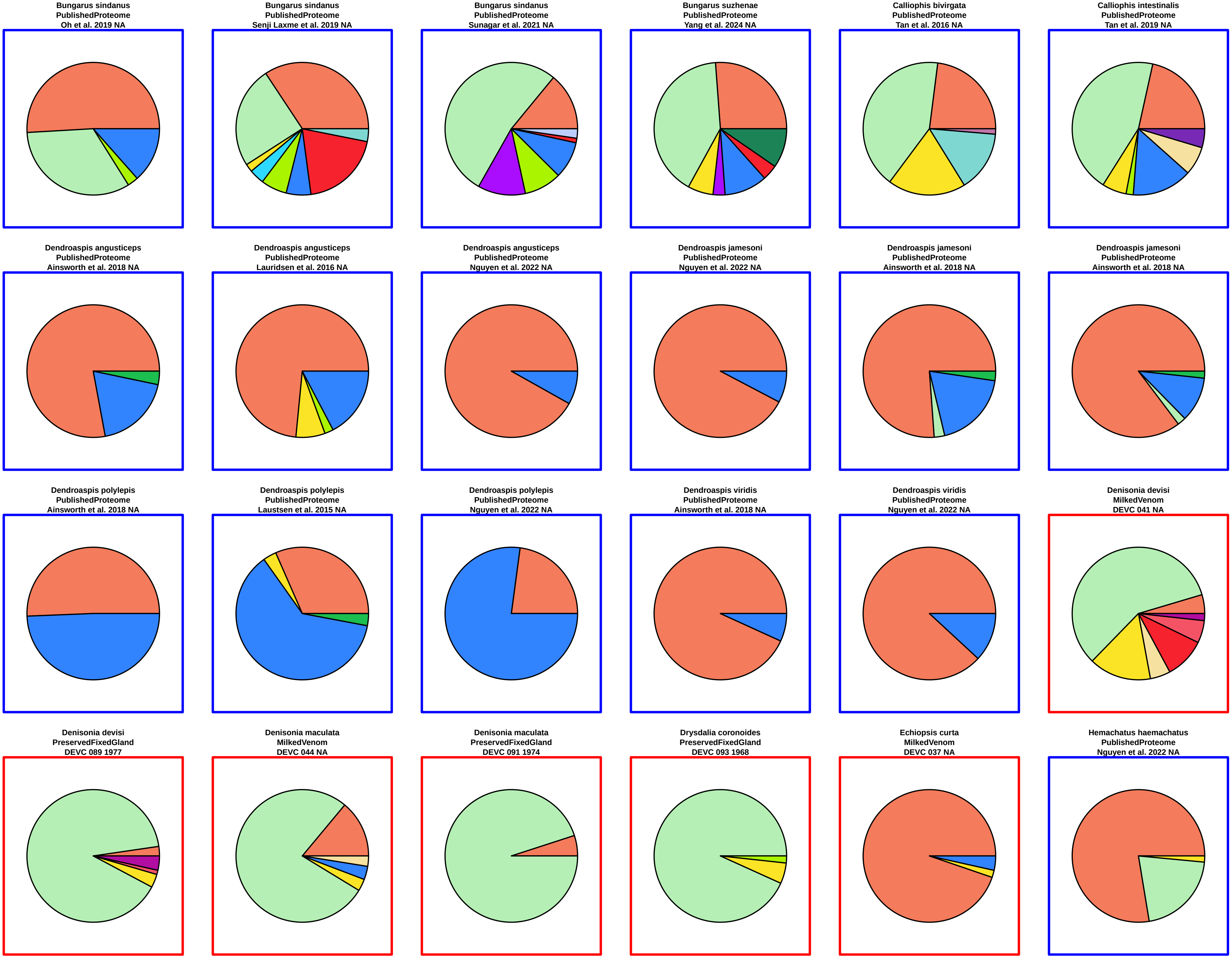

Bungarus sindanus
PublishedProteome
Oh et al. 2019 NA
Bungarus sindanus
PublishedProteome
Senji Laxme et al. 2019 NA
Bungarus sindanus
PublishedProteome
Sunagar et al. 2021 NA
Bungarus suzhenae
PublishedProteome
Yang et al. 2024 NA
Calliophis bivirgata
PublishedProteome
Tan et al. 2016 NA
Calliophis intestinalis
PublishedProteome
Tan et al. 2019 NA
Dendroaspis angusticeps
PublishedProteome
Ainsworth et al. 2018 NA
Dendroaspis angusticeps
PublishedProteome
Lauridsen et al. 2016 NA
Dendroaspis angusticeps
PublishedProteome
Nguyen et al. 2022 NA
Dendroaspis jamesoni
PublishedProteome
Nguyen et al. 2022 NA
Dendroaspis jamesoni
PublishedProteome
Ainsworth et al. 2018 NA
Dendroaspis jamesoni
PublishedProteome
Ainsworth et al. 2018 NA
Dendroaspis polylepis
PublishedProteome
Ainsworth et al. 2018 NA
Dendroaspis polylepis
PublishedProteome
Laustsen et al. 2015 NA
Dendroaspis polylepis
PublishedProteome
Nguyen et al. 2022 NA
Dendroaspis viridis
PublishedProteome
Ainsworth et al. 2018 NA
Dendroaspis viridis
PublishedProteome
Nguyen et al. 2022 NA
Denisonia devisi
MilkedVenom
DEVC 041 NA
Denisonia devisi
PreservedFixedGland
DEVC 089 1977
Denisonia maculata
MilkedVenom
DEVC 044 NA
Denisonia maculata
PreservedFixedGland
DEVC 091 1974
Drysdalia coronoides
PreservedFixedGland
DEVC 093 1968
Echiopsis curta
MilkedVenom
DEVC 037 NA
Hemachatus haemachatus
PublishedProteome
Nguyen et al. 2022 NA

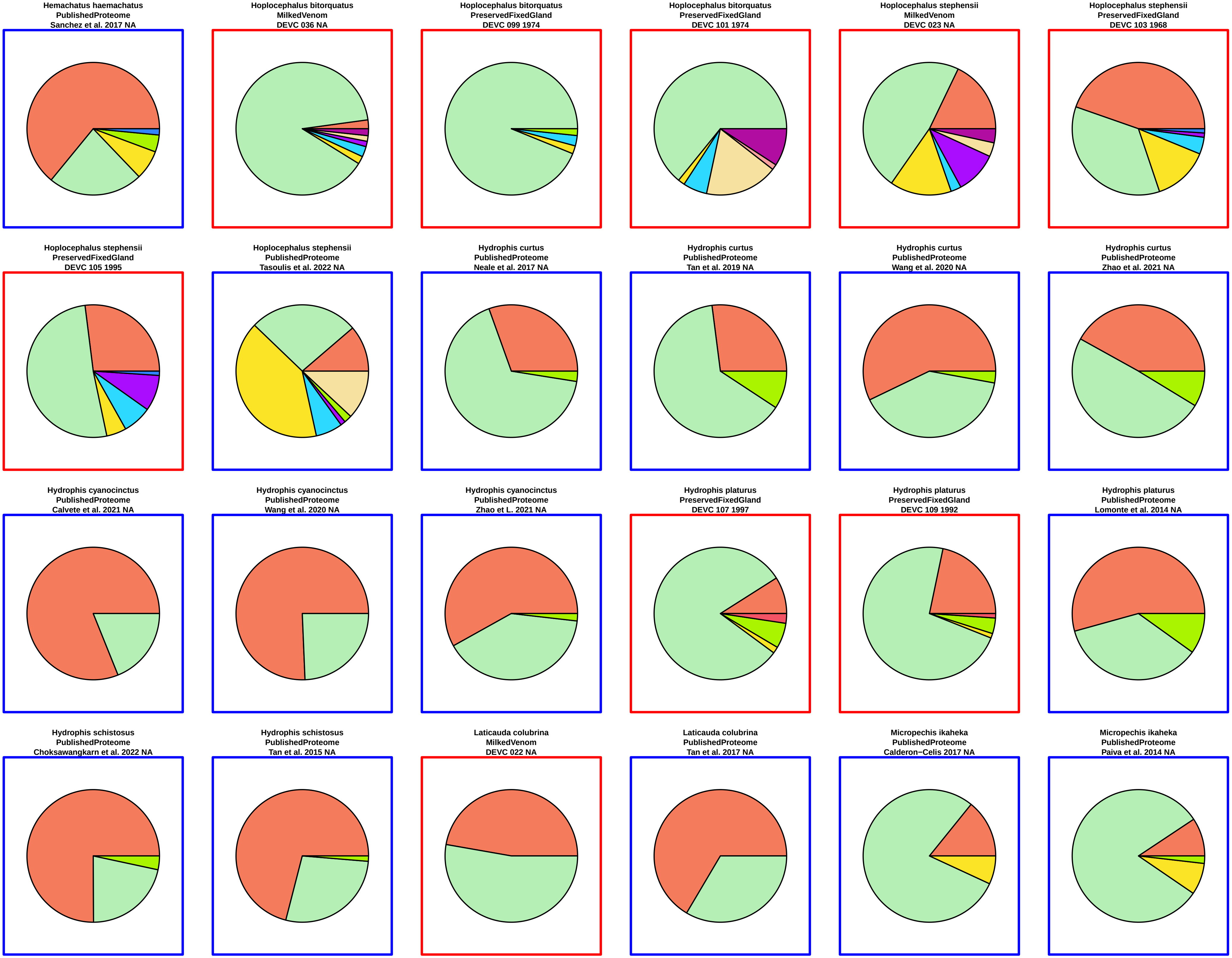
Hemachatus haemachatus
PublishedProteome
Sanchez et al. 2017 NA
Hoplocephalus bitorquatus
MilkedVenom
DEVC 036 NA
Hoplocephalus bitorquatus
PreservedFixedGland
DEVC 099 1974
Hoplocephalus bitorquatus
PreservedFixedGland
DEVC 101 1974
Hoplocephalus stephensii
MilkedVenom
DEVC 023 NA
Hoplocephalus stephensii
PreservedFixedGland
DEVC 103 1968
Hoplocephalus stephensii
PreservedFixedGland
DEVC 105 1995
Hoplocephalus stephensii
PublishedProteome
Tasoulis et al. 2022 NA
Hydrophis curtus
PublishedProteome
Neale et al. 2017 NA
Hydrophis curtus
PublishedProteome
Tan et al. 2019 NA
Hydrophis curtus
PublishedProteome
Wang et al. 2020 NA
Hydrophis curtus
PublishedProteome
Zhao et al. 2021 NA
Hydrophis cyanocinctus
PublishedProteome
Calvete et al. 2021 NA
Hydrophis cyanocinctus
PublishedProteome
Wang et al. 2020 NA
Hydrophis cyanocinctus
PublishedProteome
Zhao et L. 2021 NA
Hydrophis platurus
PreservedFixedGland
DEVC 107 1997
Hydrophis platurus
PreservedFixedGland
DEVC 109 1992
Hydrophis platurus
PublishedProteome
Lomonte et al. 2014 NA
Hydrophis schistosus
PublishedProteome
Choksawangkarn et al. 2022 NA
Hydrophis schistosus
PublishedProteome
Tan et al. 2015 NA
Laticauda colubrina
MilkedVenom
DEVC 022 NA
Laticauda colubrina
PublishedProteome
Tan et al. 2017 NA
Micropechis ikaheka
PublishedProteome
Calderon−Celis 2017 NA
Micropechis ikaheka
PublishedProteome
Paiva et al. 2014 NA

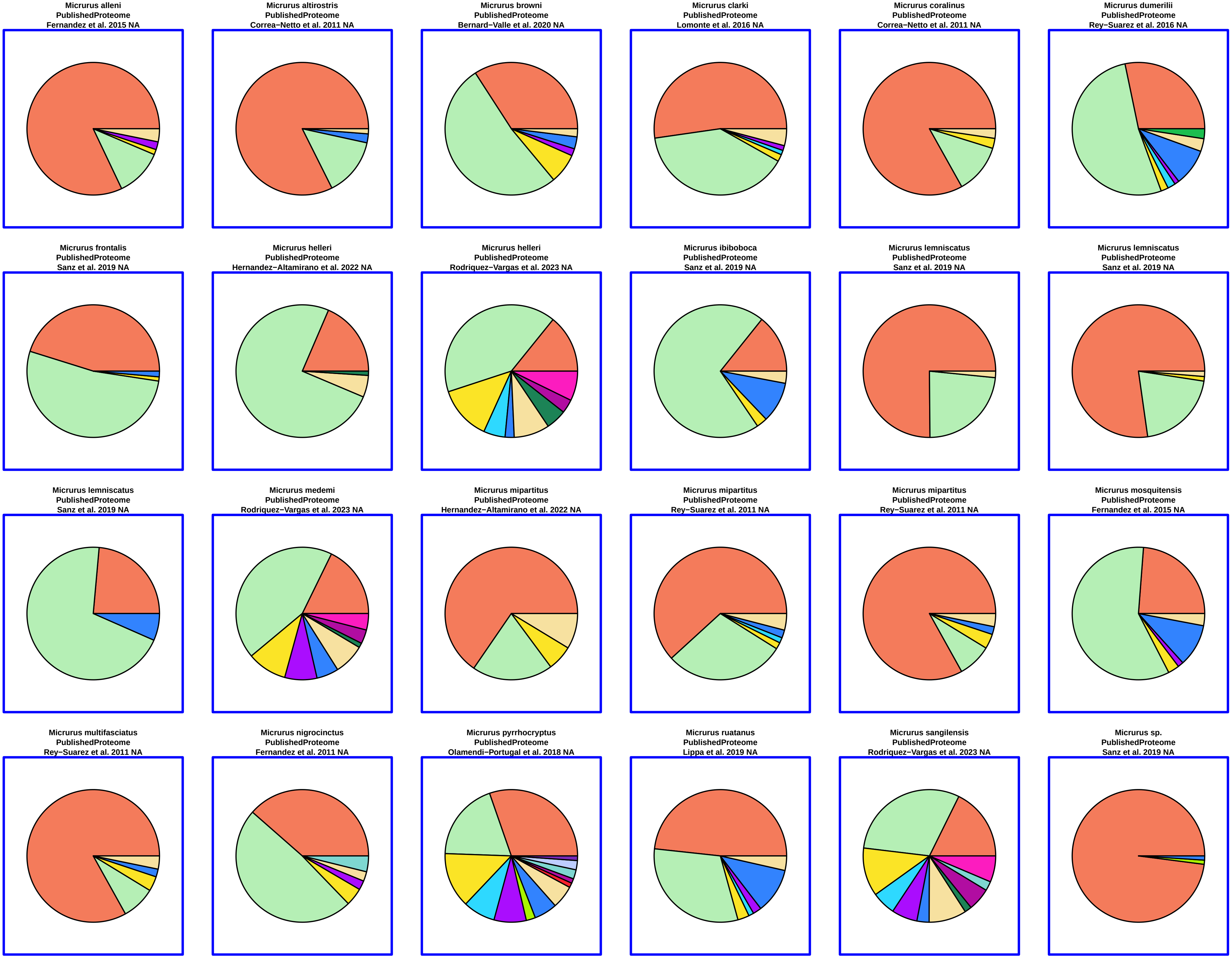

Micrurus alleni
PublishedProteome
Fernandez et al. 2015 NA
Micrurus altirostris
PublishedProteome
Correa−Netto et al. 2011 NA
Micrurus browni
PublishedProteome
Bernard−Valle et al. 2020 NA
Micrurus clarki
PublishedProteome
Lomonte et al. 2016 NA
Micrurus coralinus
PublishedProteome
Correa−Netto et al. 2011 NA
Micrurus dumerilii
PublishedProteome
Rey−Suarez et al. 2016 NA
Micrurus frontalis
PublishedProteome
Sanz et al. 2019 NA
Micrurus helleri
PublishedProteome
Hernandez−Altamirano et al. 2022 NA
Micrurus helleri
PublishedProteome
Rodriquez−Vargas et al. 2023 NA
Micrurus ibiboboca
PublishedProteome
Sanz et al. 2019 NA
Micrurus lemniscatus
PublishedProteome
Sanz et al. 2019 NA
Micrurus lemniscatus
PublishedProteome
Sanz et al. 2019 NA
Micrurus lemniscatus
PublishedProteome
Sanz et al. 2019 NA
Micrurus medemi
PublishedProteome
Rodriquez−Vargas et al. 2023 NA
Micrurus mipartitus
PublishedProteome
Hernandez−Altamirano et al. 2022 NA
Micrurus mipartitus
PublishedProteome
Rey−Suarez et al. 2011 NA
Micrurus mipartitus
PublishedProteome
Rey−Suarez et al. 2011 NA
Micrurus mosquitensis
PublishedProteome
Fernandez et al. 2015 NA
Micrurus multifasciatus
PublishedProteome
Rey−Suarez et al. 2011 NA
Micrurus nigrocinctus
PublishedProteome
Fernandez et al. 2011 NA
Micrurus pyrrhocryptus
PublishedProteome
Olamendi−Portugal et al. 2018 NA
Micrurus ruatanus
PublishedProteome
Lippa et al. 2019 NA
Micrurus sangilensis
PublishedProteome
Rodriquez−Vargas et al. 2023 NA
Micrurus sp.
PublishedProteome
Sanz et al. 2019 NA

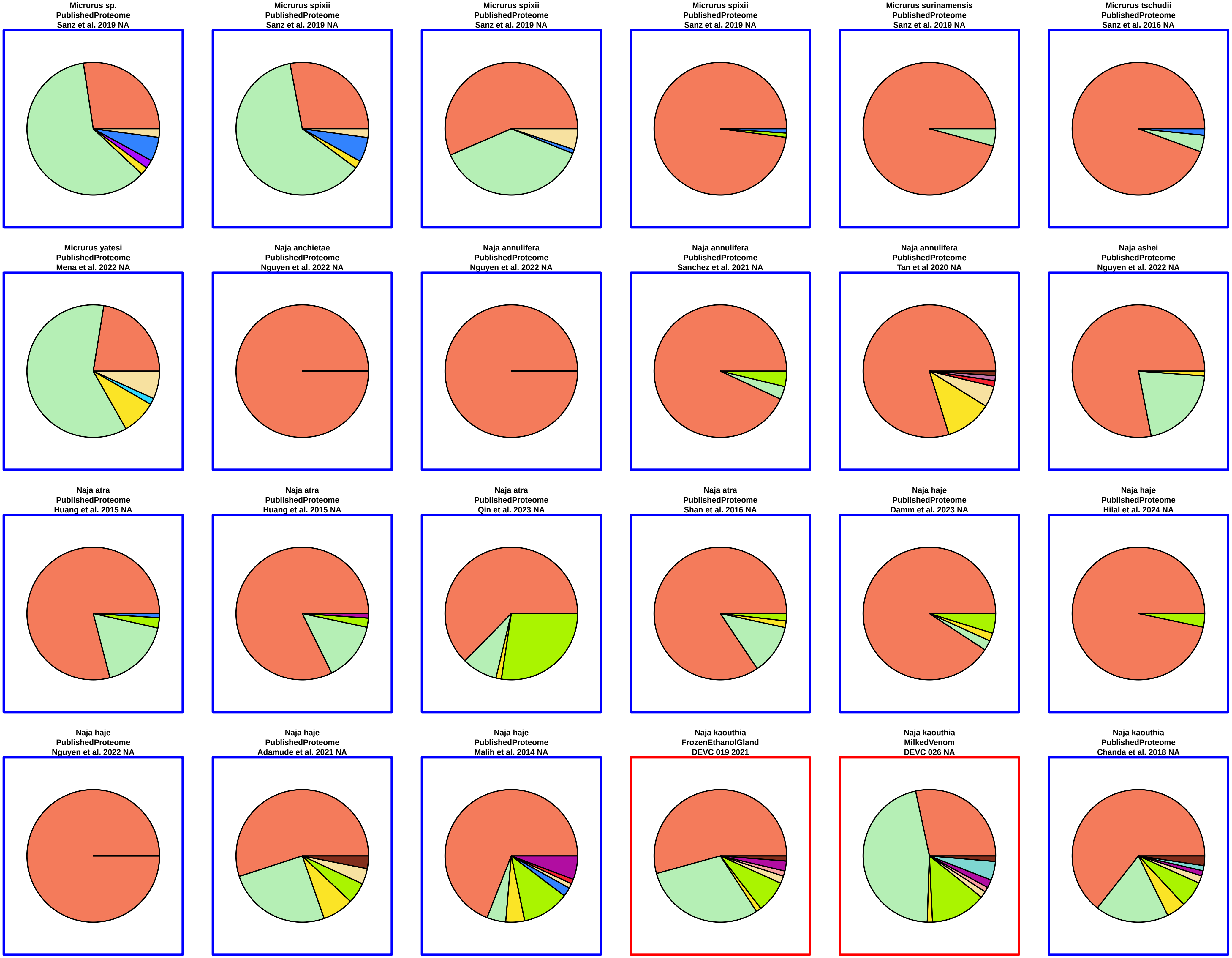
Micrurus sp.
PublishedProteome
Sanz et al. 2019 NA
Micrurus spixii
PublishedProteome
Sanz et al. 2019 NA
Micrurus spixii
PublishedProteome
Sanz et al. 2019 NA
Micrurus spixii
PublishedProteome
Sanz et al. 2019 NA
Micrurus surinamensis
PublishedProteome
Sanz et al. 2019 NA
Micrurus tschudii
PublishedProteome
Sanz et al. 2016 NA
Micrurus yatesi
PublishedProteome
Mena et al. 2022 NA
Naja anchietae
PublishedProteome
Nguyen et al. 2022 NA
Naja annulifera
PublishedProteome
Nguyen et al. 2022 NA
Naja annulifera
PublishedProteome
Sanchez et al. 2021 NA
Naja annulifera
PublishedProteome
Tan et al 2020 NA
Naja ashei
PublishedProteome
Nguyen et al. 2022 NA
Naja atra
PublishedProteome
Huang et al. 2015 NA
Naja atra
PublishedProteome
Huang et al. 2015 NA
Naja atra
PublishedProteome
Qin et al. 2023 NA
Naja atra
PublishedProteome
Shan et al. 2016 NA
Naja haje
PublishedProteome
Damm et al. 2023 NA
Naja haje
PublishedProteome
Hilal et al. 2024 NA
Naja haje
PublishedProteome
Nguyen et al. 2022 NA
Naja haje
PublishedProteome
Adamude et al. 2021 NA
Naja haje
PublishedProteome
Malih et al. 2014 NA
Naja kaouthia
FrozenEthanolGland
DEVC 019 2021
Naja kaouthia
MilkedVenom
DEVC 026 NA
Naja kaouthia
PublishedProteome
Chanda et al. 2018 NA

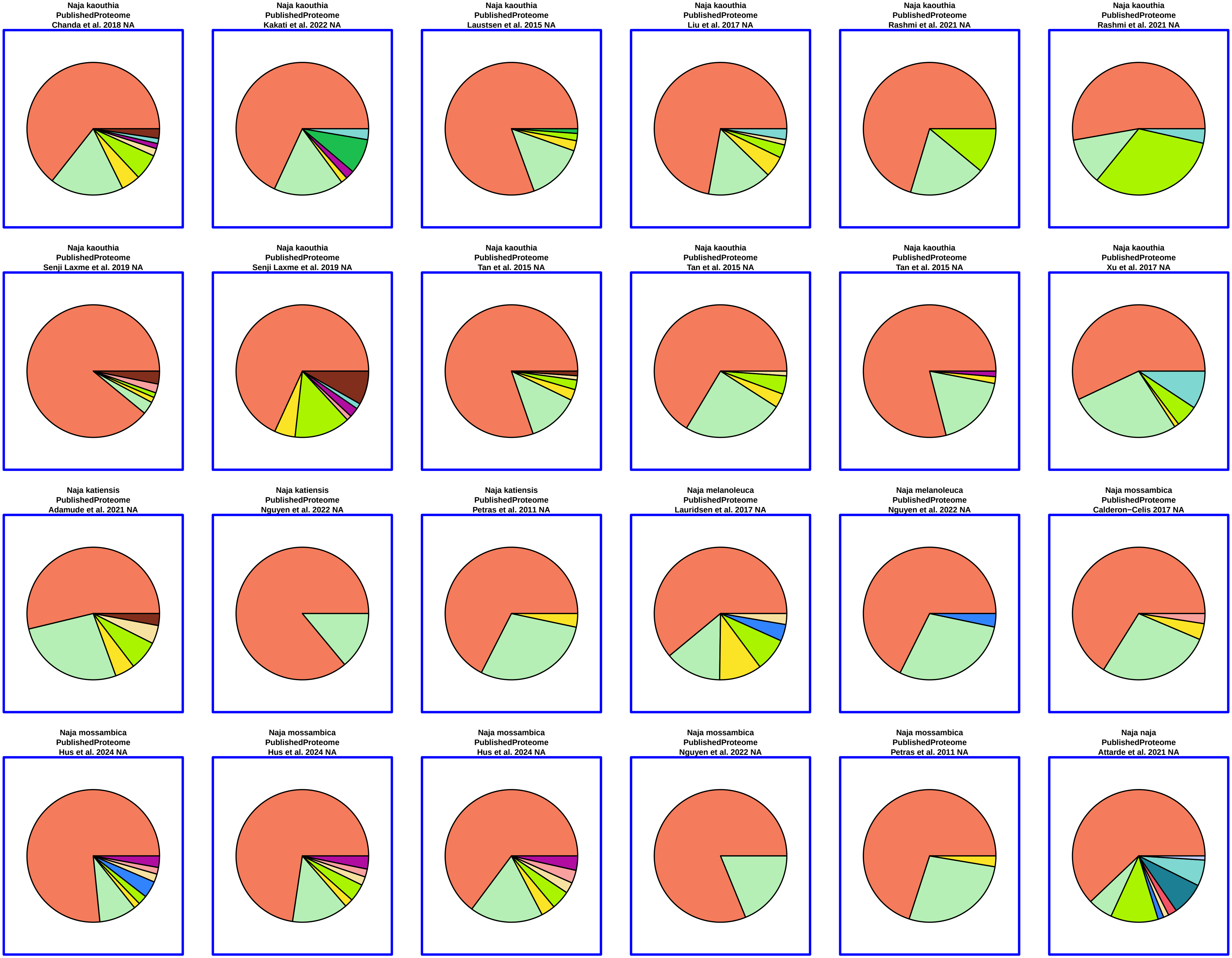
Naja kaouthia
PublishedProteome
Chanda et al. 2018 NA
Naja kaouthia
PublishedProteome
Kakati et al. 2022 NA
Naja kaouthia
PublishedProteome
Laustsen et al. 2015 NA
Naja kaouthia
PublishedProteome
Liu et al. 2017 NA
Naja kaouthia
PublishedProteome
Rashmi et al. 2021 NA
Naja kaouthia
PublishedProteome
Rashmi et al. 2021 NA
Naja kaouthia
PublishedProteome
Senji Laxme et al. 2019 NA
Naja kaouthia
PublishedProteome
Senji Laxme et al. 2019 NA
Naja kaouthia
PublishedProteome
Tan et al. 2015 NA
Naja kaouthia
PublishedProteome
Tan et al. 2015 NA
Naja kaouthia
PublishedProteome
Tan et al. 2015 NA
Naja kaouthia
PublishedProteome
Xu et al. 2017 NA
Naja katiensis
PublishedProteome
Adamude et al. 2021 NA
Naja katiensis
PublishedProteome
Nguyen et al. 2022 NA
Naja katiensis
PublishedProteome
Petras et al. 2011 NA
Naja melanoleuca
PublishedProteome
Lauridsen et al. 2017 NA
Naja melanoleuca
PublishedProteome
Nguyen et al. 2022 NA
Naja mossambica
PublishedProteome
Calderon−Celis 2017 NA
Naja mossambica
PublishedProteome
Hus et al. 2024 NA
Naja mossambica
PublishedProteome
Hus et al. 2024 NA
Naja mossambica
PublishedProteome
Hus et al. 2024 NA
Naja mossambica
PublishedProteome
Nguyen et al. 2022 NA
Naja mossambica
PublishedProteome
Petras et al. 2011 NA
Naja naja
PublishedProteome
Attarde et al. 2021 NA

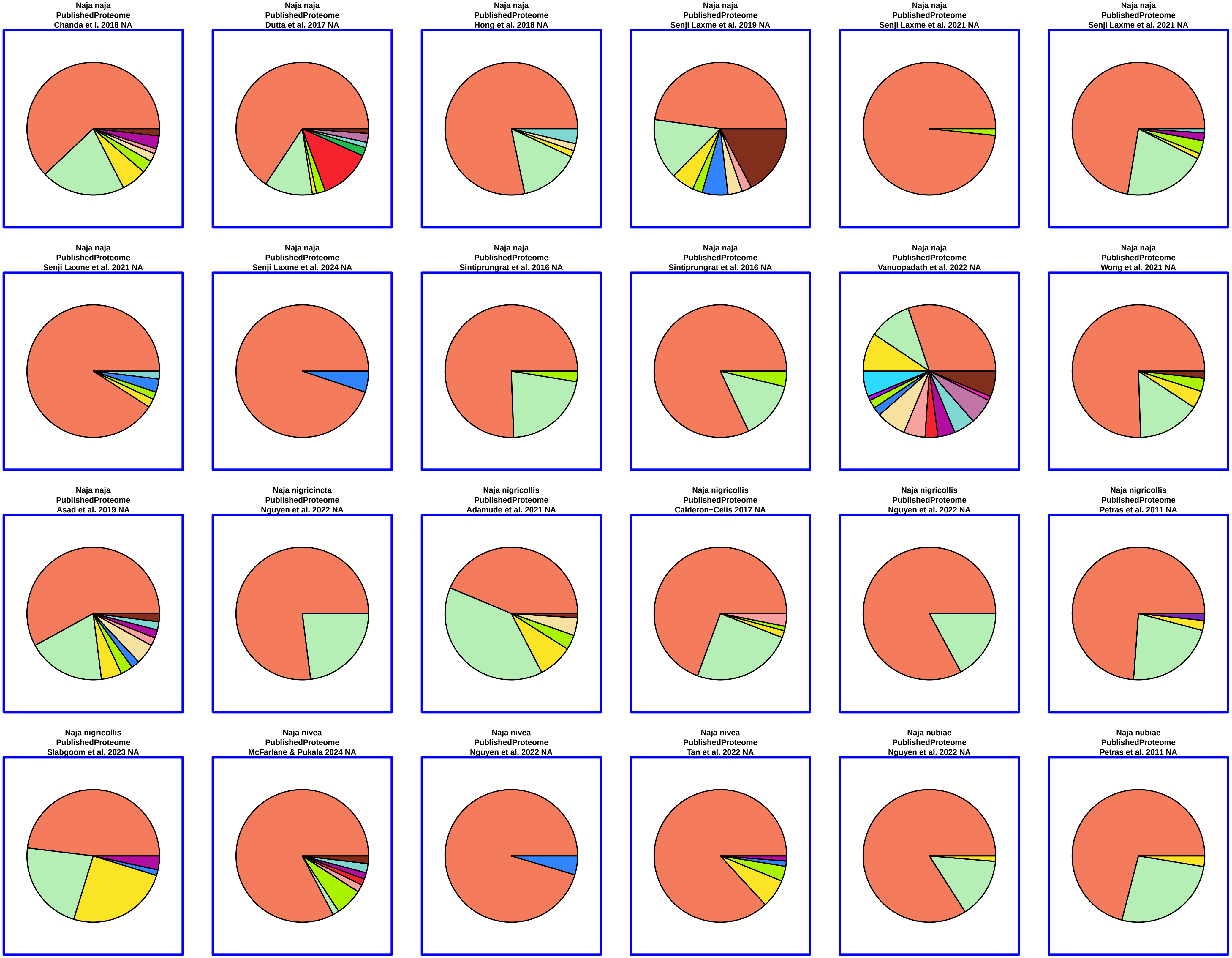
Naja naja
PublishedProteome
Chanda et I. 2018 NA
Naja naja
PublishedProteome
Dutta et al. 2017 NA
Naja naja
PublishedProteome
Hong et al. 2018 NA
Naja naja
PublishedProteome
Senji Laxme et al. 2019 NA
Naja naja
PublishedProteome
Senji Laxme et al. 2021 NA
Naja naja
PublishedProteome
Senji Laxme et al. 2021 NA
Naja naja
PublishedProteome
Senji Laxme et al. 2021 NA
Naja naja
PublishedProteome
Senji Laxme et al. 2024 NA
Naja naja
PublishedProteome
Sintiprungrat et al. 2016 NA
Naja naja
PublishedProteome
Sintiprungrat et al. 2016 NA
Naja naja
PublishedProteome
Vanuopadath et al. 2022 NA
Naja naja
PublishedProteome
Wong et al. 2021 NA
Naja naja
PublishedProteome
Asad et al. 2019 NA
Naja nigricincta
PublishedProteome
Nguyen et al. 2022 NA
Naja nigricollis
PublishedProteome
Adamude et al. 2021 NA
Naja nigricollis
PublishedProteome
Calderon−Celis 2017 NA
Naja nigricollis
PublishedProteome
Nguyen et al. 2022 NA
Naja nigricollis
PublishedProteome
Petras et al. 2011 NA
Naja nigricollis
PublishedProteome
Slabgoom et al. 2023 NA
Naja nivea
PublishedProteome
McFarlane & Pukala 2024 NA
Naja nivea
PublishedProteome
Nguyen et al. 2022 NA
Naja nivea
PublishedProteome
Tan et al. 2022 NA
Naja nubiae
PublishedProteome
Nguyen et al. 2022 NA
Naja nubiae
PublishedProteome
Petras et al. 2011 NA

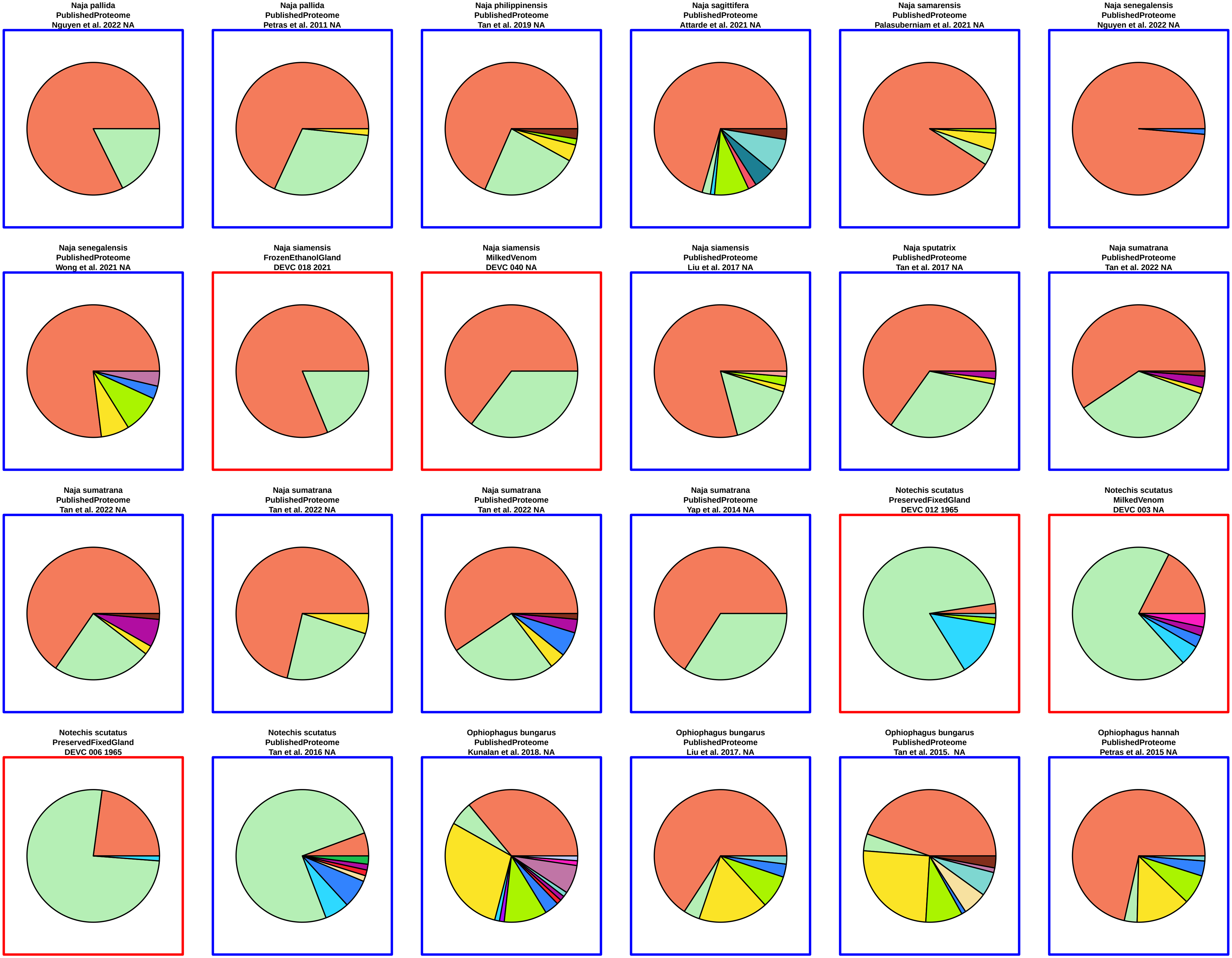
Naja pallida
PublishedProteome
Nguyen et al. 2022 NA
Naja pallida
PublishedProteome
Petras et al. 2011 NA
Naja philippinensis
PublishedProteome
Tan et al. 2019 NA
Naja sagittifera
PublishedProteome
Attarde et al. 2021 NA
Naja samarensis
PublishedProteome
Palasuberniam et al. 2021 NA
Naja senegalensis
PublishedProteome
Nguyen et al. 2022 NA
Naja senegalensis
PublishedProteome
Wong et al. 2021 NA
Naja siamensis
FrozenEthanolGland
DEVC 018 2021
Naja siamensis
MilkedVenom
DEVC 040 NA
Naja siamensis
PublishedProteome
Liu et al. 2017 NA
Naja sputatrix
PublishedProteome
Tan et al. 2017 NA
Naja sumatrana
PublishedProteome
Tan et al. 2022 NA
Naja sumatrana
PublishedProteome
Tan et al. 2022 NA
Naja sumatrana
PublishedProteome
Tan et al. 2022 NA
Naja sumatrana
PublishedProteome
Tan et al. 2022 NA
Naja sumatrana
PublishedProteome
Yap et al. 2014 NA
Notechis scutatus
PreservedFixedGland
DEVC 012 1965
Notechis scutatus
MilkedVenom
DEVC 003 NA
Notechis scutatus
PreservedFixedGland
DEVC 006 1965
Notechis scutatus
PublishedProteome
Tan et al. 2016 NA
Ophiophagus bungarus
PublishedProteome
Kunalan et al. 2018. NA
Ophiophagus bungarus
PublishedProteome
Liu et al. 2017. NA
Ophiophagus bungarus
PublishedProteome
Tan et al. 2015.  NA
Ophiophagus hannah
PublishedProteome
Petras et al. 2015 NA

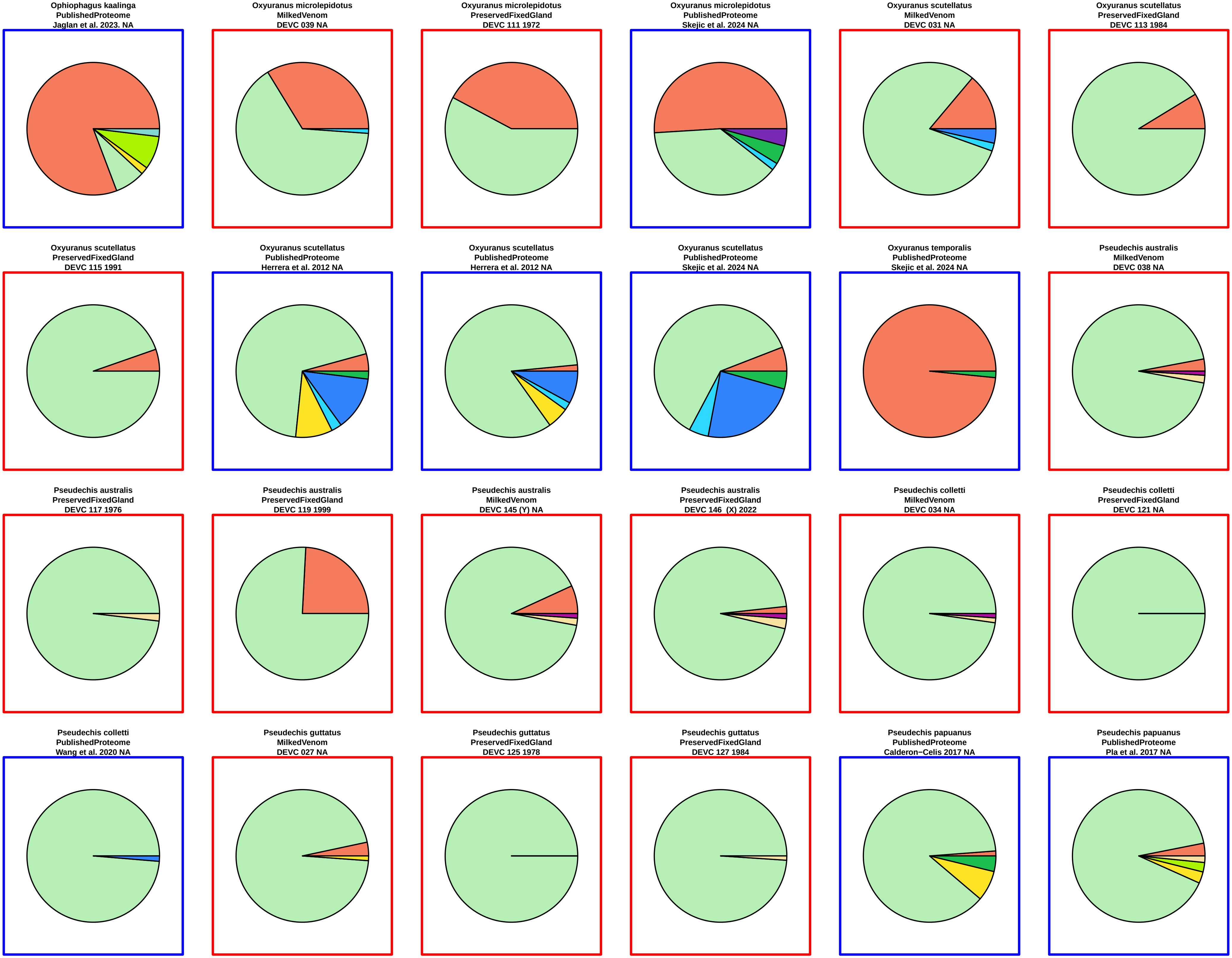
Ophiophagus kaalinga
PublishedProteome
Jaglan et al. 2023. NA
Oxyuranus microlepidotus
MilkedVenom
DEVC 039 NA
Oxyuranus microlepidotus
PreservedFixedGland
DEVC 111 1972
Oxyuranus microlepidotus
PublishedProteome
Skejic et al. 2024 NA
Oxyuranus scutellatus
MilkedVenom
DEVC 031 NA
Oxyuranus scutellatus
PreservedFixedGland
DEVC 113 1984
Oxyuranus scutellatus
PreservedFixedGland
DEVC 115 1991
Oxyuranus scutellatus
PublishedProteome
Herrera et al. 2012 NA
Oxyuranus scutellatus
PublishedProteome
Herrera et al. 2012 NA
Oxyuranus scutellatus
PublishedProteome
Skejic et al. 2024 NA
Oxyuranus temporalis
PublishedProteome
Skejic et al. 2024 NA
Pseudechis australis
MilkedVenom
DEVC 038 NA
Pseudechis australis
PreservedFixedGland
DEVC 117 1976
Pseudechis australis
PreservedFixedGland
DEVC 119 1999
Pseudechis australis
MilkedVenom
DEVC 145 (Y) NA
Pseudechis australis
PreservedFixedGland
DEVC 146 (X) 2022
Pseudechis colletti
MilkedVenom
DEVC 034 NA
Pseudechis colletti
PreservedFixedGland
DEVC 121 NA
Pseudechis colletti
PublishedProteome
Wang et al. 2020 NA
Pseudechis guttatus
MilkedVenom
DEVC 027 NA
Pseudechis guttatus
PreservedFixedGland
DEVC 125 1978
Pseudechis guttatus
PreservedFixedGland
DEVC 127 1984
Pseudechis papuanus
PublishedProteome
Calderon−Celis 2017 NA
Pseudechis papuanus
PublishedProteome
Pla et al. 2017 NA

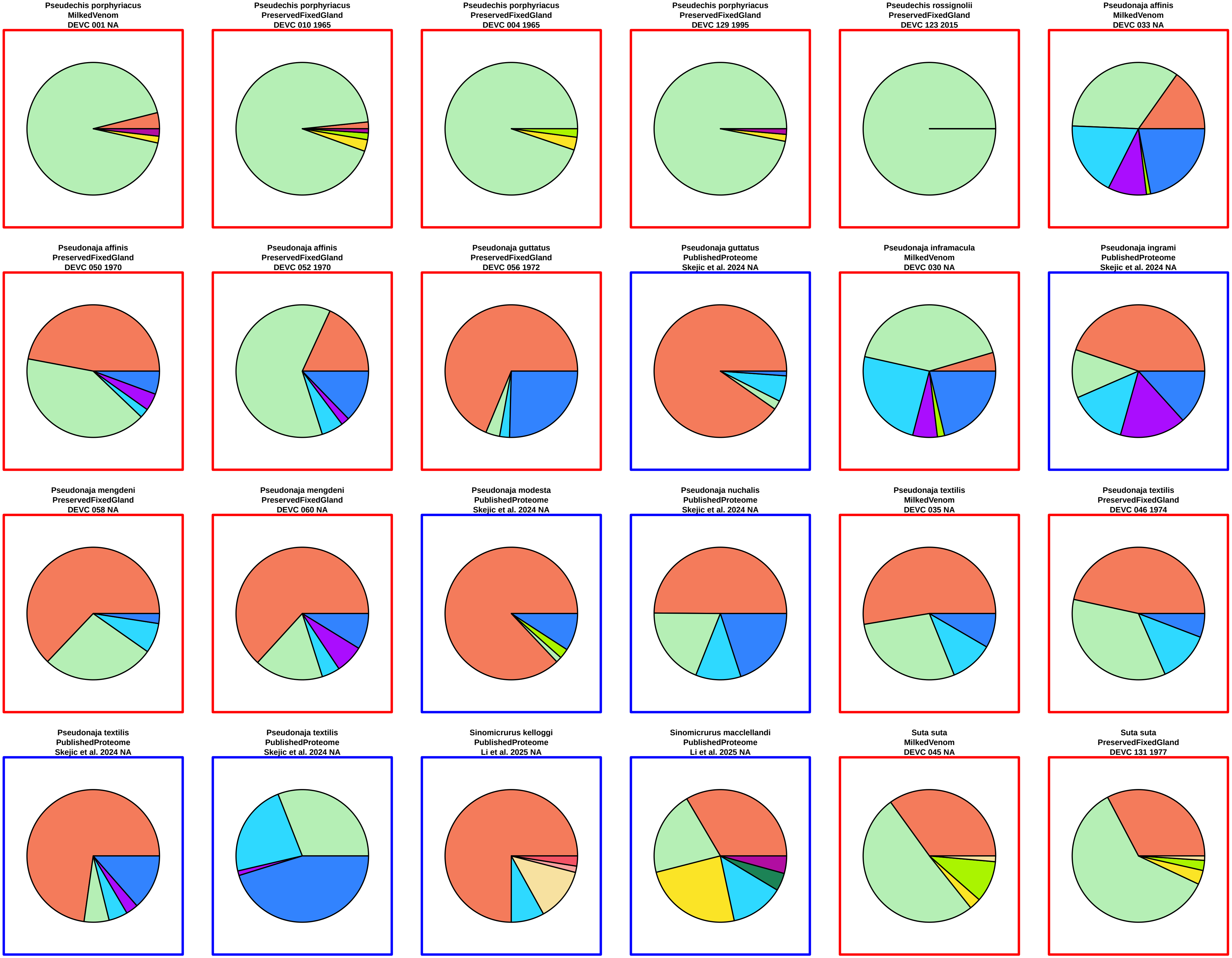
Pseudechis porphyriacus
MilkedVenom
DEVC 001 NA
Pseudechis porphyriacus
PreservedFixedGland
DEVC 010 1965
Pseudechis porphyriacus
PreservedFixedGland
DEVC 004 1965
Pseudechis porphyriacus
PreservedFixedGland
DEVC 129 1995
Pseudechis rossignolii
PreservedFixedGland
DEVC 123 2015
Pseudonaja affinis
MilkedVenom
DEVC 033 NA
Pseudonaja affinis
PreservedFixedGland
DEVC 050 1970
Pseudonaja affinis
PreservedFixedGland
DEVC 052 1970
Pseudonaja guttatus
PreservedFixedGland
DEVC 056 1972
Pseudonaja guttatus
PublishedProteome
Skejic et al. 2024 NA
Pseudonaja inframacula
MilkedVenom
DEVC 030 NA
Pseudonaja ingrami
PublishedProteome
Skejic et al. 2024 NA
Pseudonaja mengdeni
PreservedFixedGland
DEVC 058 NA
Pseudonaja mengdeni
PreservedFixedGland
DEVC 060 NA
Pseudonaja modesta
PublishedProteome
Skejic et al. 2024 NA
Pseudonaja nuchalis
PublishedProteome
Skejic et al. 2024 NA
Pseudonaja textilis
MilkedVenom
DEVC 035 NA
Pseudonaja textilis
PreservedFixedGland
DEVC 046 1974
Pseudonaja textilis
PublishedProteome
Skejic et al. 2024 NA
Pseudonaja textilis
PublishedProteome
Skejic et al. 2024 NA
Sinomicrurus kelloggi
PublishedProteome
Li et al. 2025 NA
Sinomicrurus macclellandi
PublishedProteome
Li et al. 2025 NA
Suta suta
MilkedVenom
DEVC 045 NA
Suta suta
PreservedFixedGland
DEVC 131 1977

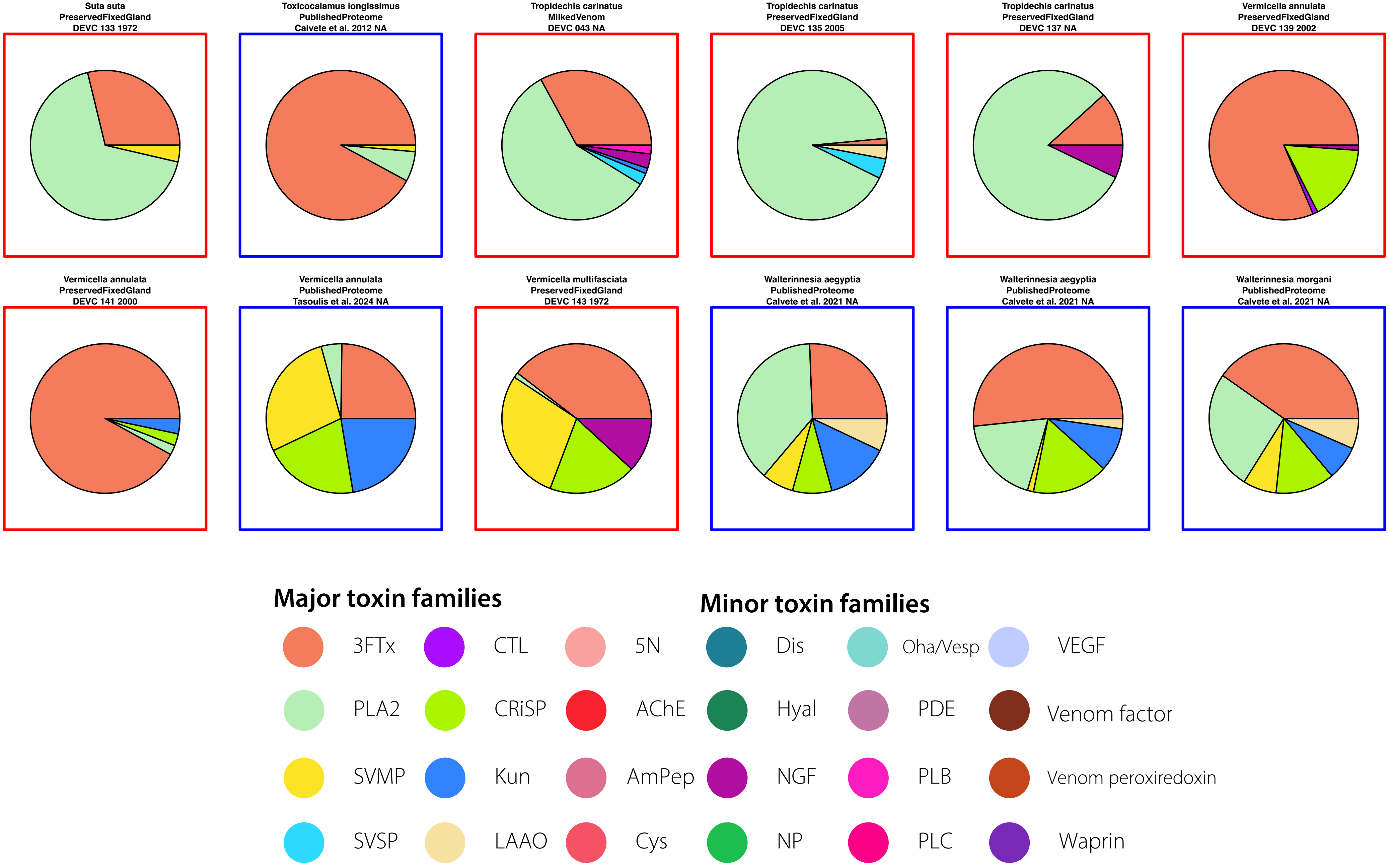
Suta suta
PreservedFixedGland
DEVC 133 1972
Toxicocalamus longissimus
PublishedProteome
Calvete et al. 2012 NA
Tropidechis carinatus
MilkedVenom
DEVC 043 NA
Tropidechis carinatus
PreservedFixedGland
DEVC 135 2005
Tropidechis carinatus
PreservedFixedGland
DEVC 137 NA
Vermicella annulata
PreservedFixedGland
DEVC 139 2002
Vermicella annulata
PreservedFixedGland
DEVC 141 2000
Vermicella annulata
PublishedProteome
Tasoulis et al. 2024 NA
Vermicella multifasciata
PreservedFixedGland
DEVC 143 1972
Walterinnesia aegyptia
PublishedProteome
Calvete et al. 2021 NA
Walterinnesia aegyptia
PublishedProteome
Calvete et al. 2021 NA
Walterinnesia morgani
PublishedProteome
Calvete et al. 2021 NA
Major toxin families
3FTx
CTL
5N
PLA2
CRiSP
AChE
SVMP
Kun
AmPep
SVSP
LAAO
Cys
Minor toxin families
Dis
Oha/Vesp
VEGF
Hyal
PDE
Venom factor
NGF
PLB
Venom peroxiredoxin
NP
PLC
Waprin

Agkistrodon bilineatus
MilkedVenom
DEVC 024 NA

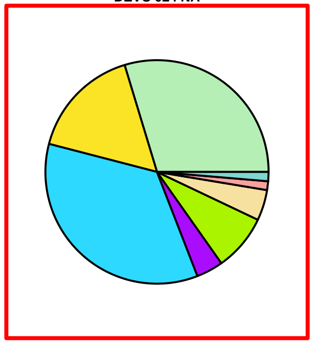

Agkistrodon bilineatus
PublishedProteome
Lomonte et al. 2014 NA

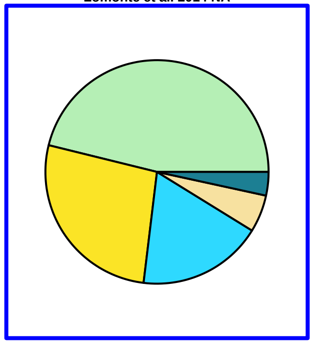

Agkistrodon conanti
PublishedProteome
Lomonte et al. 2014 NA

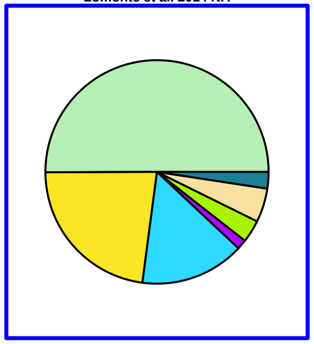

Agkistrodon contortrix
PublishedProteome
Bocian et al. 2016 NA

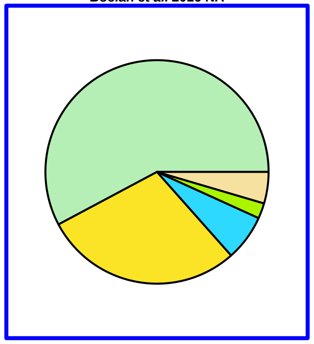

Agkistrodon contortrix
PublishedProteome
Lomonte et al. 2014 NA

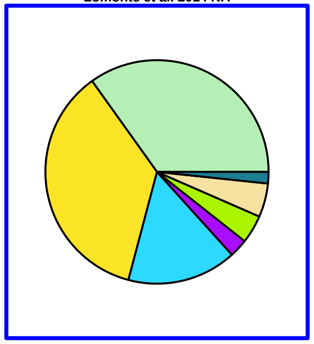

Agkistrodon contortrix
PublishedProteome
Lomonte et al. 2014 NA

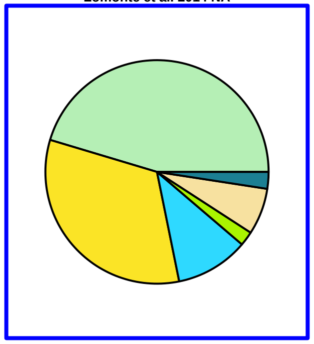

Agkistrodon contortrix
PublishedProteome
Lomonte et al. 2015 NA

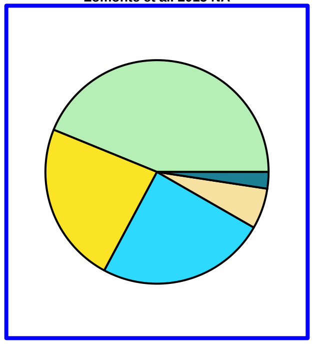

Agkistrodon howardgloydi
PublishedProteome
Lomonte et al. 2014 NA

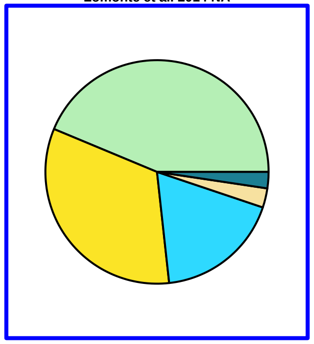

Agkistrodon laticinctus
PublishedProteome
Lomonte et al. 2014 NA

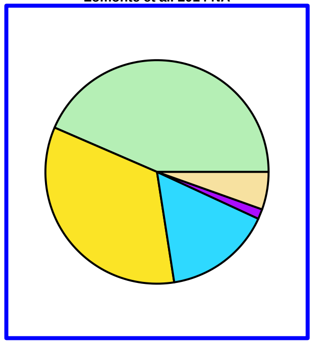

Agkistrodon laticinctus
PublishedProteome
Lomonte et al. 2014 NA

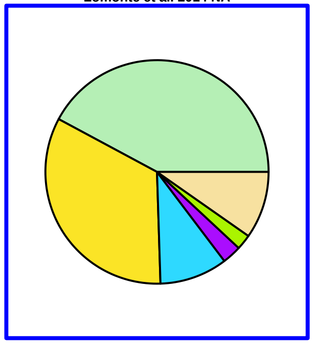

Agkistrodon piscivorus
PublishedProteome
Lomonte et al. 2014 NA

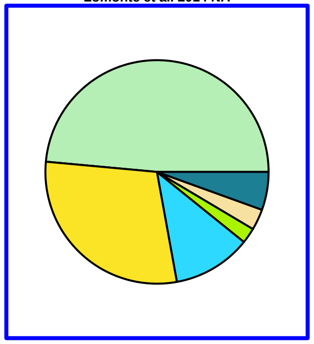

Agkistrodon piscivorus
PublishedProteome
Lomonte et al. 2014 NA

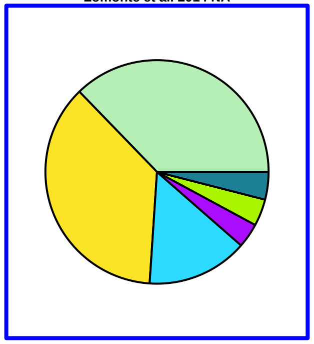

Agkistrodon taylori
PublishedProteome
Lomonte et al. 2014 NA

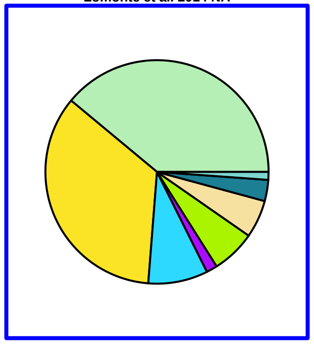

Atropoides picadoi
PublishedProteome
Angulo et al. 2008 NA

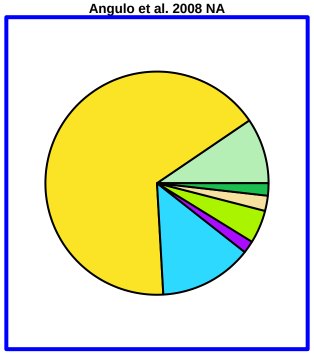

Bitis arietans
FrozenEthanolGland
DEVC 017 2021

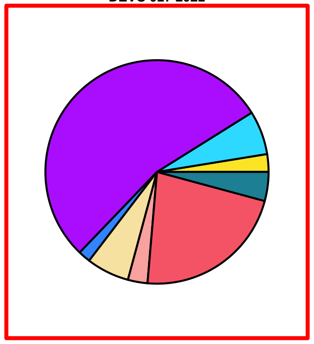

Bitis arietans
MilkedVenom
DEVC 028 NA

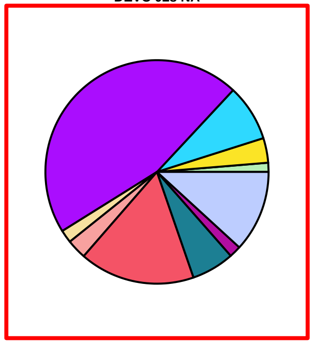

Bitis arietans
PublishedProteome
Dingwoke et al. 2021 NA

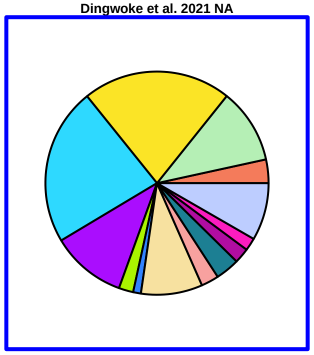

Bitis arietans
PublishedProteome
Juarez et al. 2006 NA

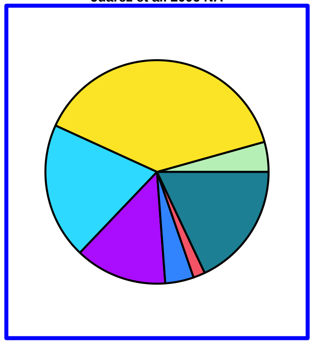

Bitis arietans
PublishedProteome
Nguyen et al. 2022 NA

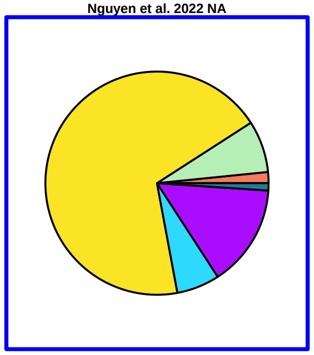

Bitis arietans
PublishedProteome
Wang et al. 2020 NA

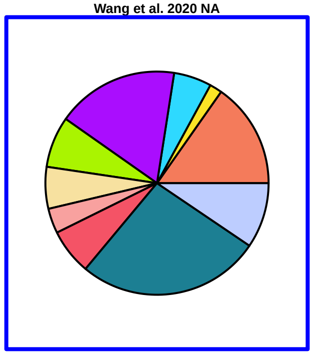

Bitis caudalis
PublishedProteome
Calvete et al. 2007 NA

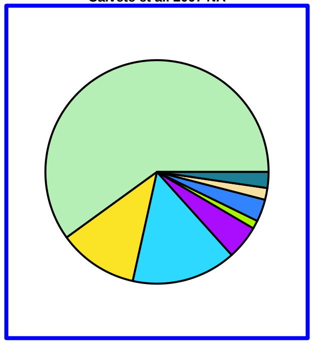

Bitis gabonica
PublishedProteome
Calvete et al. 2007 NA

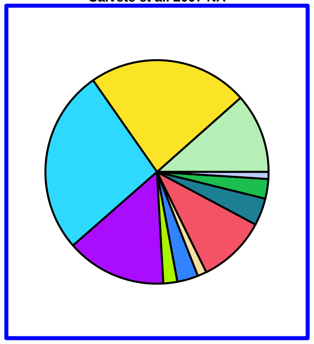

Bitis gabonica
PublishedProteome
Nguyen et al. 2022 NA

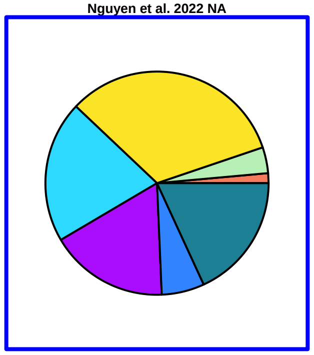

Bitis nasicornis
PublishedProteome
Calvete et al. 2007 NA

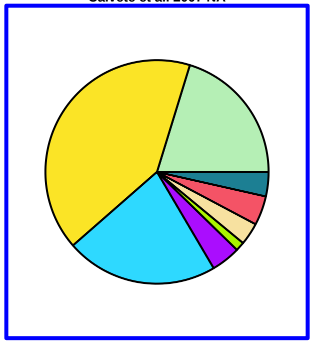

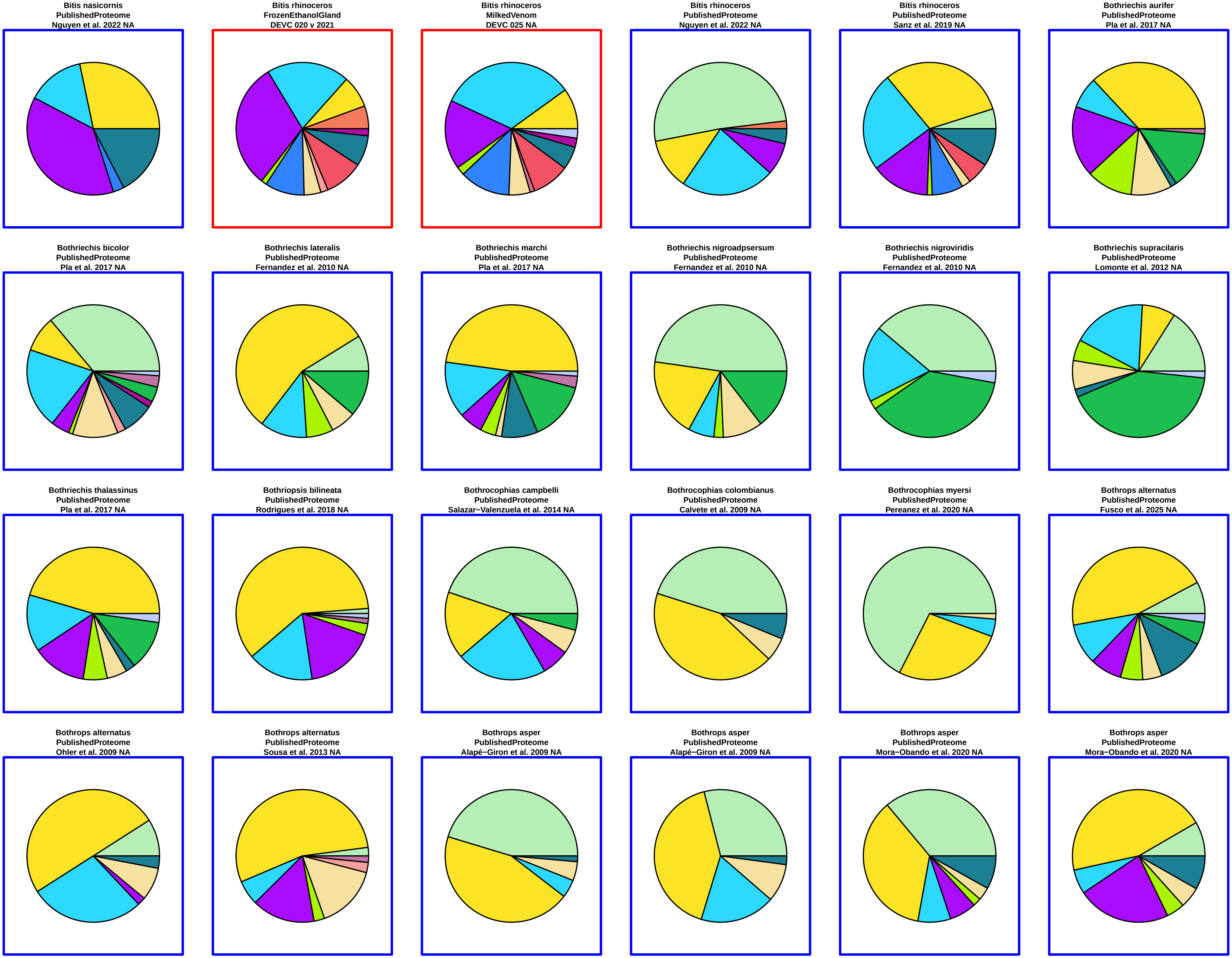
Bitis nasicornis
PublishedProteome
Nguyen et al. 2022 NA
Bitis rhinoceros
FrozenEthanolGland
DEVC 020 v 2021
Bitis rhinoceros
MilkedVenom
DEVC 025 NA
Bitis rhinoceros
PublishedProteome
Nguyen et al. 2022 NA
Bitis rhinoceros
PublishedProteome
Sanz et al. 2019 NA
Bothriechis aurifer
PublishedProteome
Pla et al. 2017 NA
Bothriechis bicolor
PublishedProteome
Pla et al. 2017 NA
Bothriechis lateralis
PublishedProteome
Fernandez et al. 2010 NA
Bothriechis marchi
PublishedProteome
Pla et al. 2017 NA
Bothriechis nigroadpsersum
PublishedProteome
Fernandez et al. 2010 NA
Bothriechis nigroviridis
PublishedProteome
Fernandez et al. 2010 NA
Bothriechis supracilaris
PublishedProteome
Lomonte et al. 2012 NA
Bothriechis thalassinus
PublishedProteome
Pla et al. 2017 NA
Bothriopsis bilineata
PublishedProteome
Rodrigues et al. 2018 NA
Bothrocophias campbelli
PublishedProteome
Salazar−Valenzuela et al. 2014 NA
Bothrocophias colombianus
PublishedProteome
Calvete et al. 2009 NA
Bothrocophias myersi
PublishedProteome
Pereanez et al. 2020 NA
Bothrops alternatus
PublishedProteome
Fusco et al. 2025 NA
Bothrops alternatus
PublishedProteome
Ohler et al. 2009 NA
Bothrops alternatus
PublishedProteome
Sousa et al. 2013 NA
Bothrops asper
PublishedProteome
Alapé−Giron et al. 2009 NA
Bothrops asper
PublishedProteome
Alapé−Giron et al. 2009 NA
Bothrops asper
PublishedProteome
Mora−Obando et al. 2020 NA
Bothrops asper
PublishedProteome
Mora−Obando et al. 2020 NA

Bothrops asper
PublishedProteome
Mora−Obando et al. 2020 NA

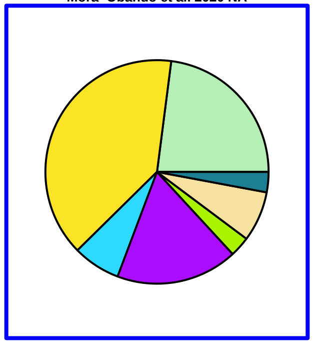

Bothrops asper
PublishedProteome
Mora−Obando et al. 2020 NA

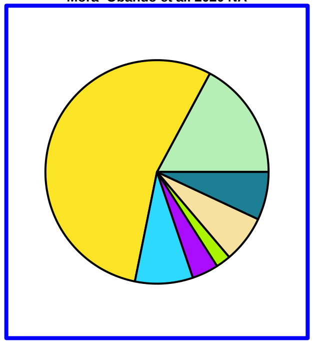

Bothrops asper
PublishedProteome
Mora−Obando et al. 2020 NA

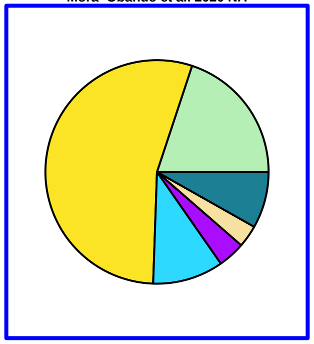

Bothrops asper
PublishedProteome
Mora−Obando et al. 2020 NA

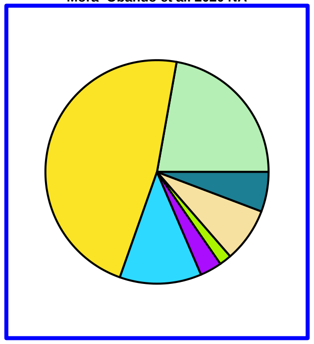

Bothrops asper
PublishedProteome
Mora−Obando et al. 2020 NA

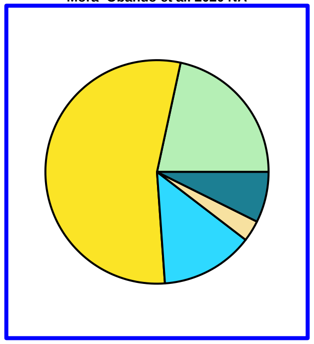

Bothrops asper
PublishedProteome
Mora−Obando et al. 2020 NA

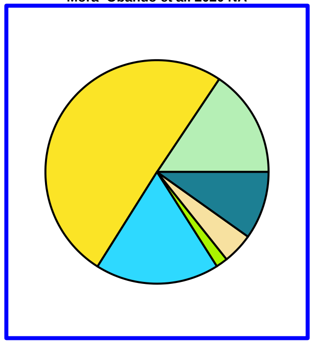

Bothrops asper
PublishedProteome
Mora−Obando et al. 2020 NA

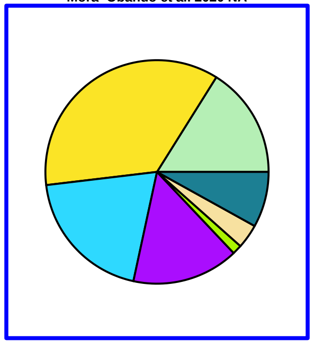

Bothrops atrox
PublishedProteome
Amazonas et al. 2018 NA

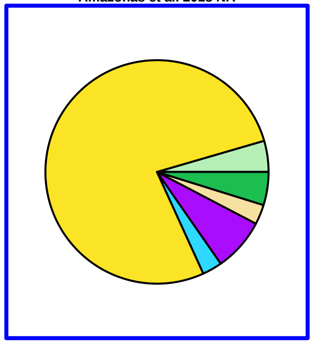

Bothrops atrox
PublishedProteome
Amazonas et al. 2018 NA

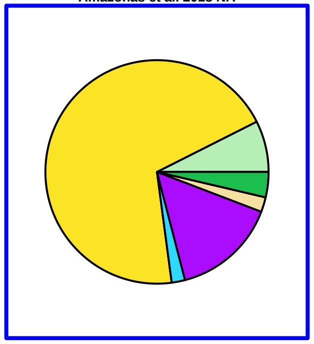

Bothrops atrox
PublishedProteome
Amazonas et al. 2018 NA

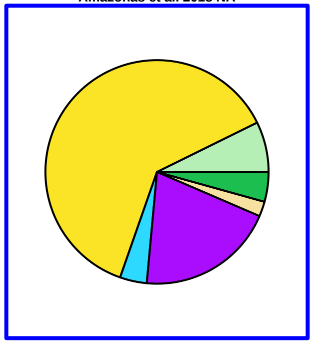

Bothrops atrox
PublishedProteome
Amazonas et al. 2018 NA

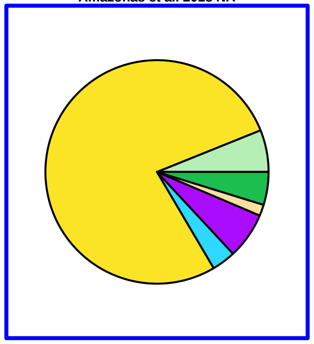

Bothrops atrox
PublishedProteome
Amazonas et al. 2018 NA

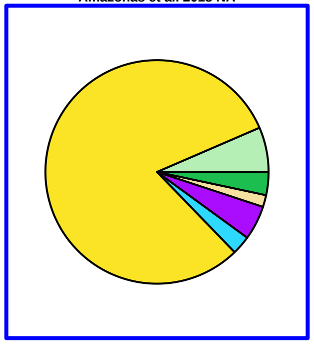

Bothrops atrox
PublishedProteome
Kohlhoff et al. 2012 NA

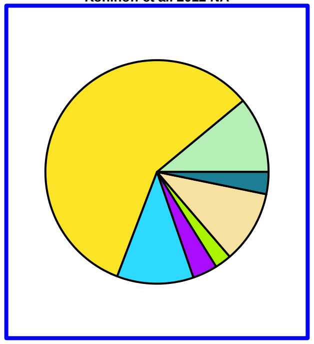

Bothrops atrox
PublishedProteome
Nunez et a. 2009 NA

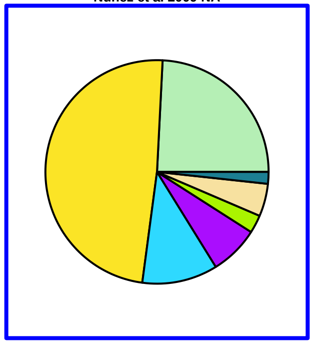

Bothrops atrox
PublishedProteome
Nunez et a. 2009 NA

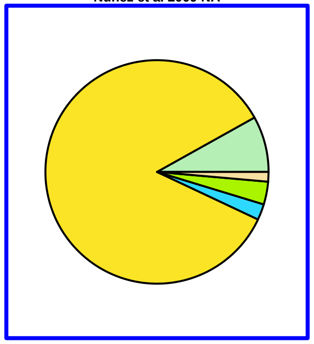

Bothrops atrox
PublishedProteome
Sousa et al. 2017 NA

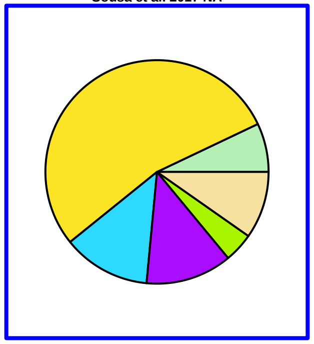

Bothrops ayerbei
PublishedProteome
Mora−Obando et al. 2014 NA

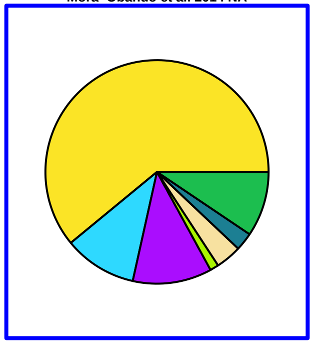

Bothrops barnetti
PublishedProteome
Kohlhoff et al. 2012 NA

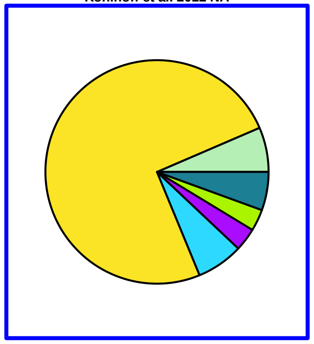

Bothrops bilineatus
PublishedProteome
Sanz et al. 2018 NA

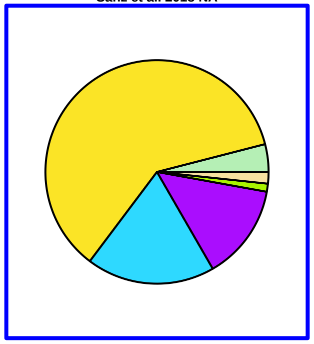

Bothrops barnetti
PublishedProteome
Sanz et al. 2018 NA

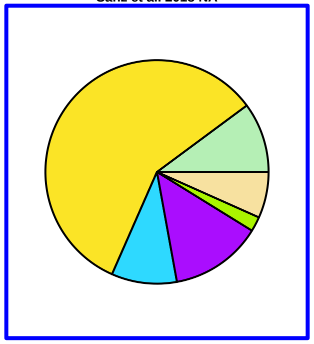

Bothrops barnetti
PublishedProteome
Sanz et al. 2018 NA

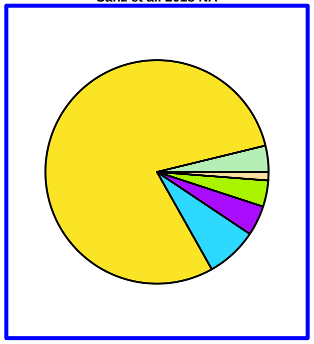

Bothrops brazilli
PublishedProteome
Rodrigues et al. 2020 NA

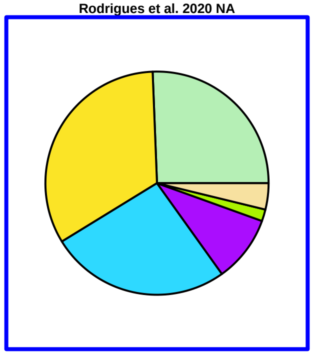

Bothrops brazilli
PublishedProteome
Sanz et al. 2020 NA

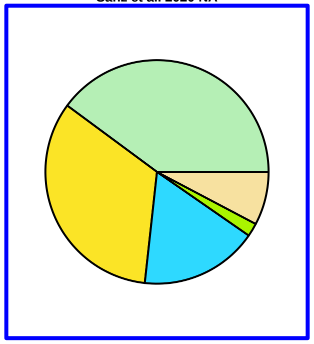

Bothrops carribbaeus
PublishedProteome
Gutierrez et al. 2008 NA

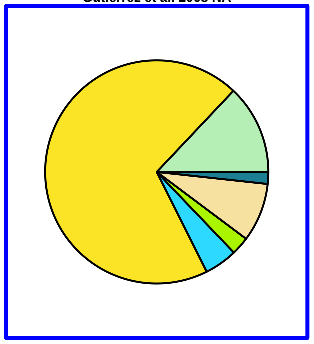

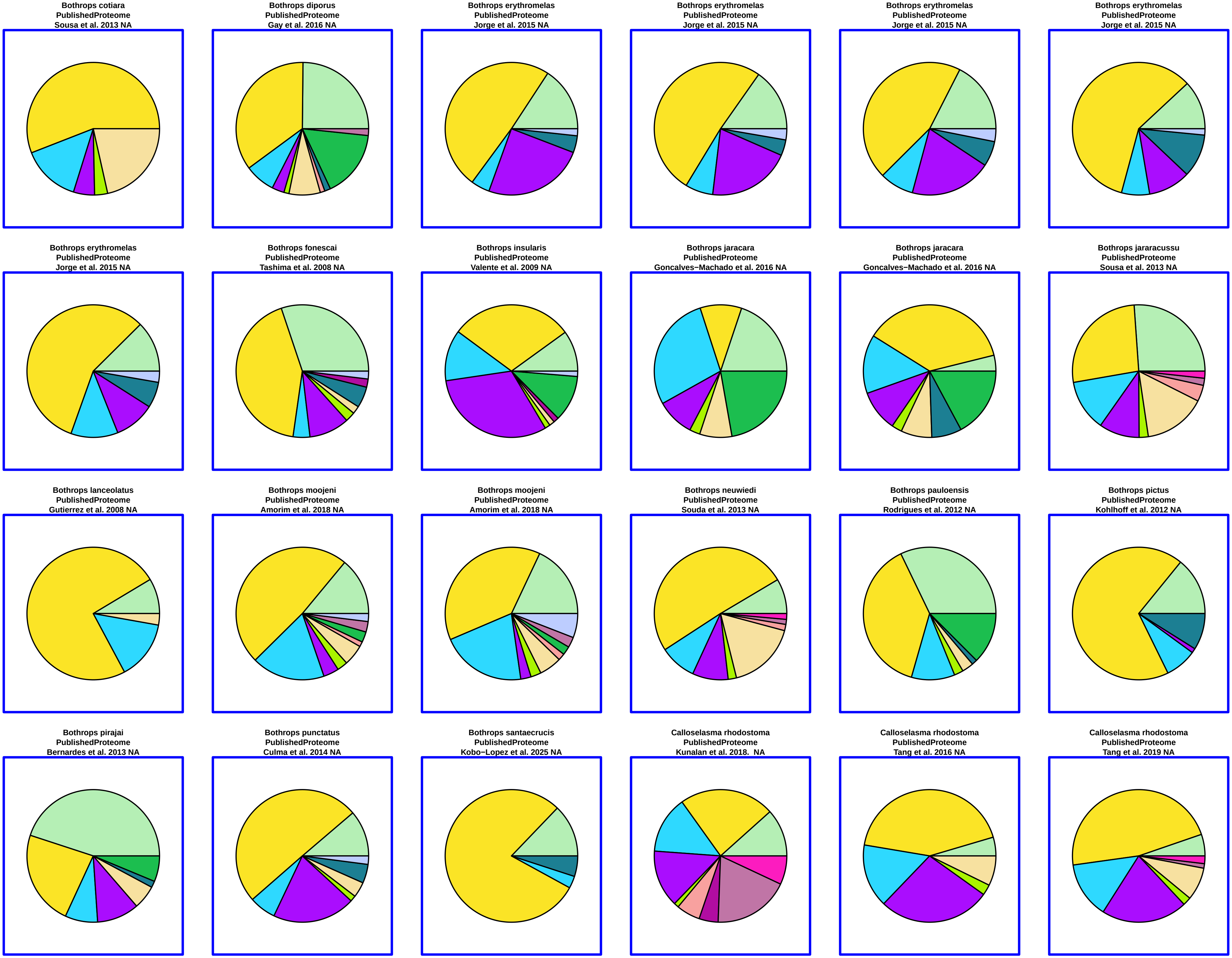
Bothrops cotiara
PublishedProteome
Sousa et al. 2013 NA
Bothrops diporus
PublishedProteome
Gay et al. 2016 NA
Bothrops erythromelas
PublishedProteome
Jorge et al. 2015 NA
Bothrops erythromelas
PublishedProteome
Jorge et al. 2015 NA
Bothrops erythromelas
PublishedProteome
Jorge et al. 2015 NA
Bothrops erythromelas
PublishedProteome
Jorge et al. 2015 NA
Bothrops erythromelas
PublishedProteome
Jorge et al. 2015 NA
Bothrops fonescai
PublishedProteome
Tashima et al. 2008 NA
Bothrops insularis
PublishedProteome
Valente et al. 2009 NA
Bothrops jaracara
PublishedProteome
Goncalves−Machado et al. 2016 NA
Bothrops jaracara
PublishedProteome
Goncalves−Machado et al. 2016 NA
Bothrops jararacussu
PublishedProteome
Sousa et al. 2013 NA
Bothrops lanceolatus
PublishedProteome
Gutierrez et al. 2008 NA
Bothrops moojeni
PublishedProteome
Amorim et al. 2018 NA
Bothrops moojeni
PublishedProteome
Amorim et al. 2018 NA
Bothrops neuwiedi
PublishedProteome
Souda et al. 2013 NA
Bothrops pauloensis
PublishedProteome
Rodrigues et al. 2012 NA
Bothrops pictus
PublishedProteome
Kohlhoff et al. 2012 NA
Bothrops pirajai
PublishedProteome
Bernardes et al. 2013 NA
Bothrops punctatus
PublishedProteome
Culma et al. 2014 NA
Bothrops santaecrucis
PublishedProteome
Kobo−Lopez et al. 2025 NA
Calloselasma rhodostoma
PublishedProteome
Kunalan et al. 2018.  NA
Calloselasma rhodostoma
PublishedProteome
Tang et al. 2016 NA
Calloselasma rhodostoma
PublishedProteome
Tang et al. 2019 NA

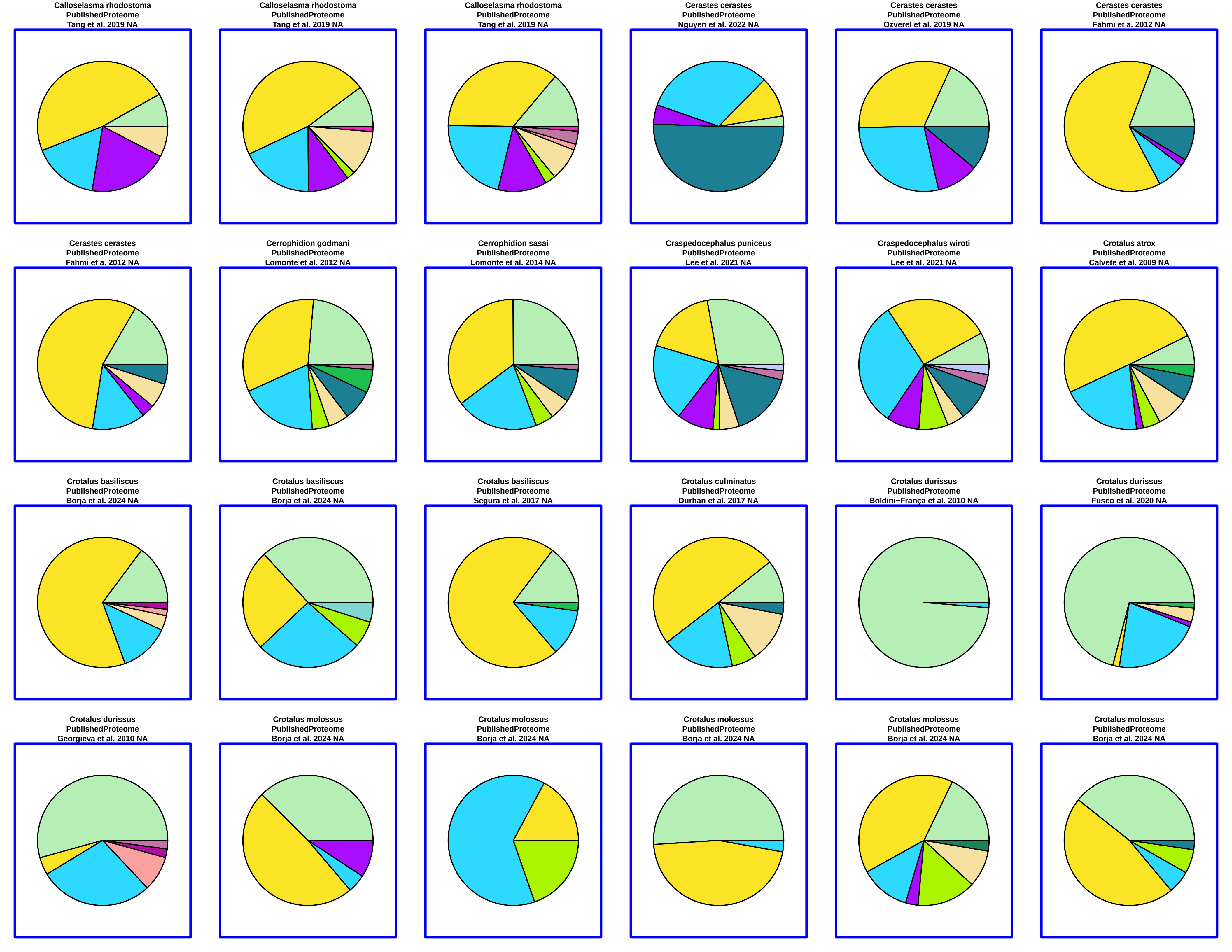
Calloselasma rhodostoma
PublishedProteome
Tang et al. 2019 NA
Calloselasma rhodostoma
PublishedProteome
Tang et al. 2019 NA
Calloselasma rhodostoma
PublishedProteome
Tang et al. 2019 NA
Cerastes cerastes
PublishedProteome
Nguyen et al. 2022 NA
Cerastes cerastes
PublishedProteome
Ozverel et al. 2019 NA
Cerastes cerastes
PublishedProteome
Fahmi et a. 2012 NA
Cerastes cerastes
PublishedProteome
Fahmi et a. 2012 NA
Cerrophidion godmani
PublishedProteome
Lomonte et al. 2012 NA
Cerrophidion sasai
PublishedProteome
Lomonte et al. 2014 NA
Craspedocephalus puniceus
PublishedProteome
Lee et al. 2021 NA
Craspedocephalus wiroti
PublishedProteome
Lee et al. 2021 NA
Crotalus atrox
PublishedProteome
Calvete et al. 2009 NA
Crotalus basiliscus
PublishedProteome
Borja et al. 2024 NA
Crotalus basiliscus
PublishedProteome
Borja et al. 2024 NA
Crotalus basiliscus
PublishedProteome
Segura et al. 2017 NA
Crotalus culminatus
PublishedProteome
Durban et al. 2017 NA
Crotalus durissus
PublishedProteome
Boldini−França et al. 2010 NA
Crotalus durissus
PublishedProteome
Fusco et al. 2020 NA
Crotalus durissus
PublishedProteome
Georgieva et al. 2010 NA
Crotalus molossus
PublishedProteome
Borja et al. 2024 NA
Crotalus molossus
PublishedProteome
Borja et al. 2024 NA
Crotalus molossus
PublishedProteome
Borja et al. 2024 NA
Crotalus molossus
PublishedProteome
Borja et al. 2024 NA
Crotalus molossus
PublishedProteome
Borja et al. 2024 NA

**Crotalus polystictus**
**PublishedProteome**
**Mackessy et al. 2018 NA**

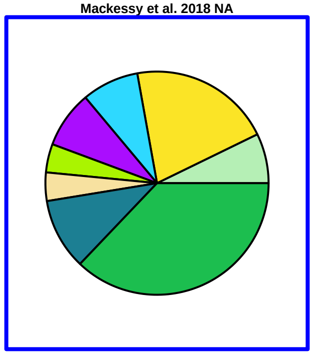

**Crotalus simus**
**PublishedProteome**
**Calvete et al. 2009 NA**

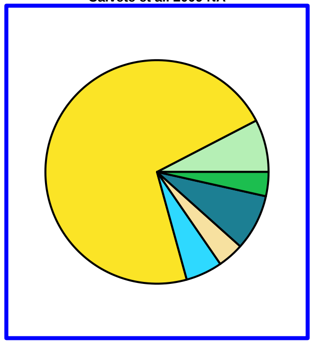

**Crotalus simus**
**PublishedProteome**
**Durban et al. 2017 NA**

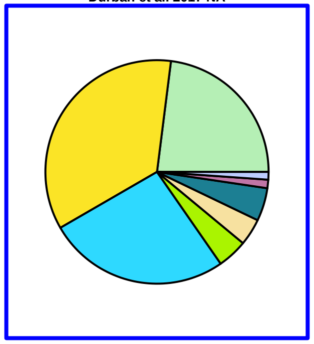

**Crotalus simus**
**PublishedProteome**
**Castro et al 2013 NA**

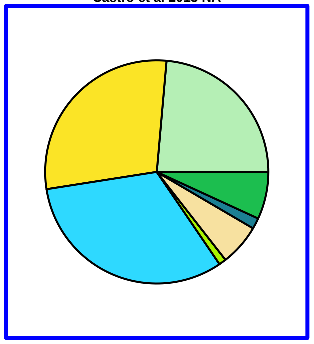

**Crotalus tigris**
**PublishedProteome**
**Calvete et al. 2012 NA**

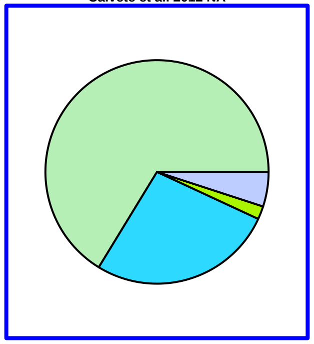

**Crotalus tzabcan**
**PublishedProteome**
**Durban et al. 2017 NA**

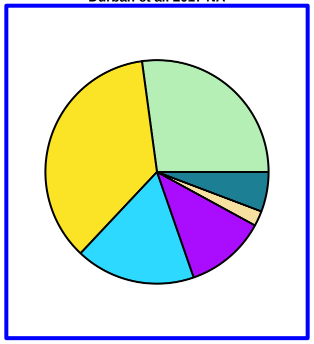

**Crotalus vegrandis**
**FrozenEthanolGland**
**DEVC 021 v 2021**

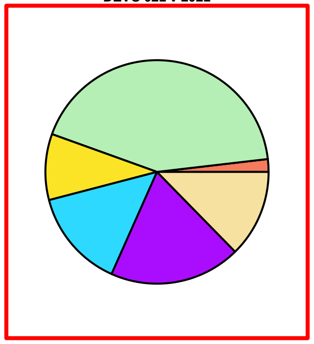

**Crotalus vegrandis**
**MilkedVenom**
**DEVC 032 NA**

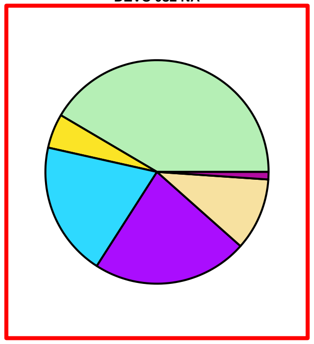

**Crotalus viridis**
**PublishedProteome**
**Saviola et al. 2015 NA**

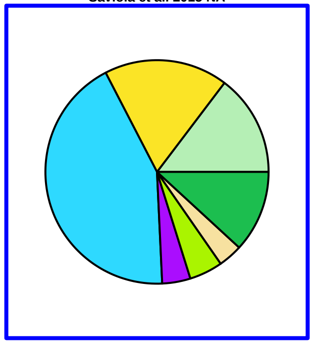

**Daboia mauritanica**
**PublishedProteome**
**Makran et al. 2012 NA**

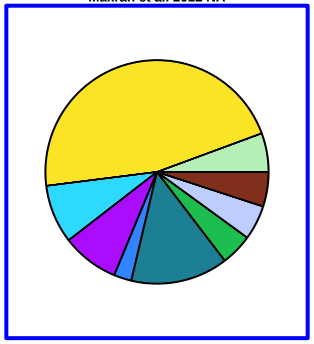

**Daboia russelii**
**PublishedProteome**
**Kalita et al. 2017 NA**

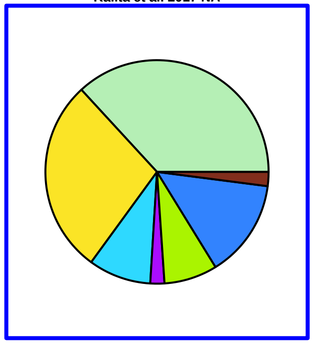

**Daboia russelii**
**PublishedProteome**
**Mukherjee et al. 2016 NA**

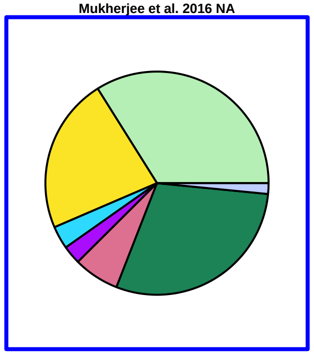

**Daboia russelii**
**PublishedProteome**
**Pla et al. 2019 NA**

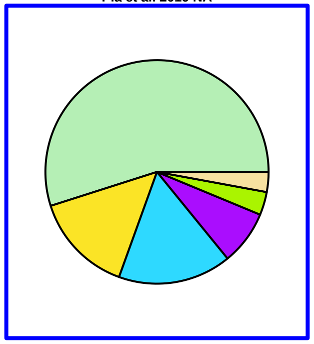

**Daboia russelii**
**PublishedProteome**
**Pla et al. 2019 NA**

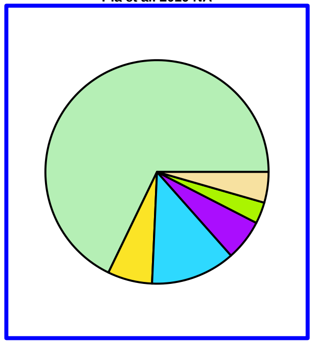

**Daboia russelii**
**PublishedProteome**
**Pla et al. 2019 NA**

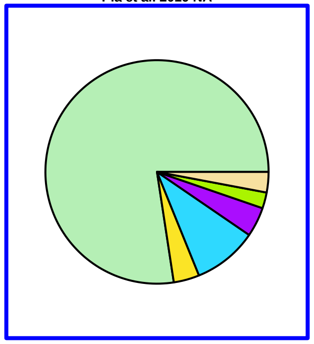

**Daboia russelii**
**PublishedProteome**
**Pla et al. 2019 NA**

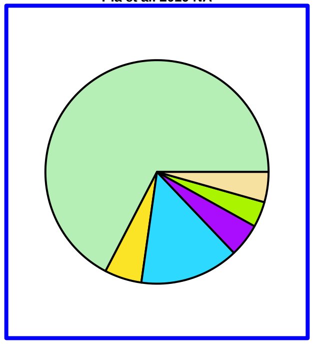

**Daboia russelii**
**PublishedProteome**
**Senji Laxme et al. 2021 NA**

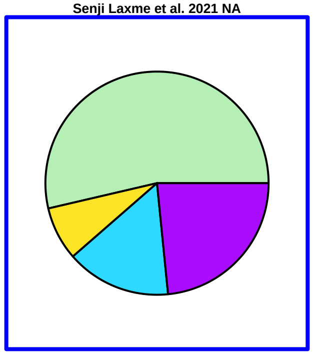

**Daboia russelii**
**PublishedProteome**
**Senji Laxme et al. 2021 NA**

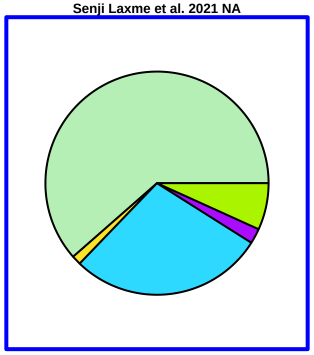

**Daboia russelii**
**PublishedProteome**
**Senji Laxme et al. 2021 NA**

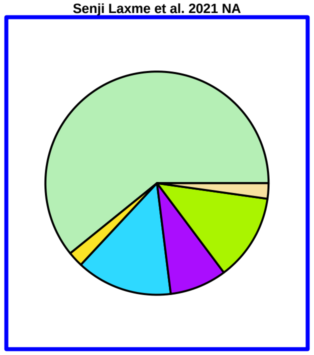

**Daboia russelii**
**PublishedProteome**
**Senji Laxme et al. 2024 NA**

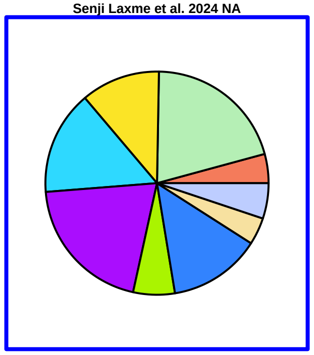

**Daboia russelii**
**PublishedProteome**
**Tan et al. 2015 NA**

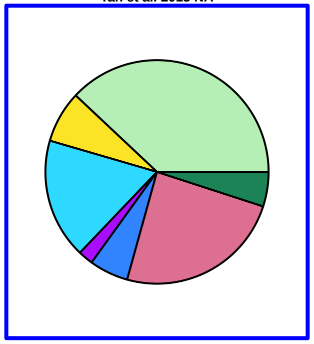

**Daboia siamensis**
**PublishedProteome**
**Lingam et al. 2020 NA**

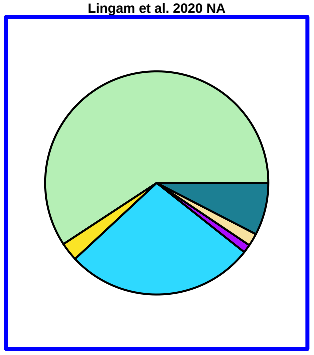

**Daboia siamensis**
**PublishedProteome**
**Lingam et al. 2020 NA**

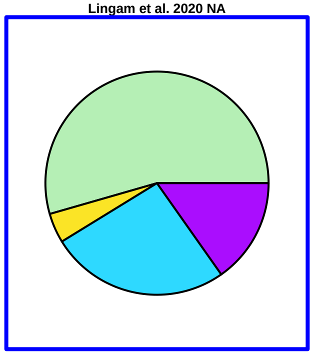

**Daboia siamensis**
**PublishedProteome**
**Tan et al. 2018 NA**

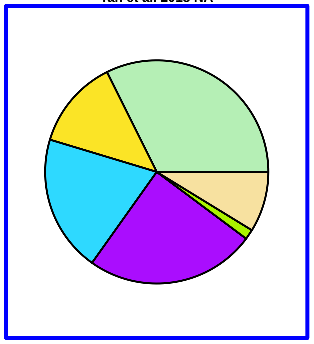

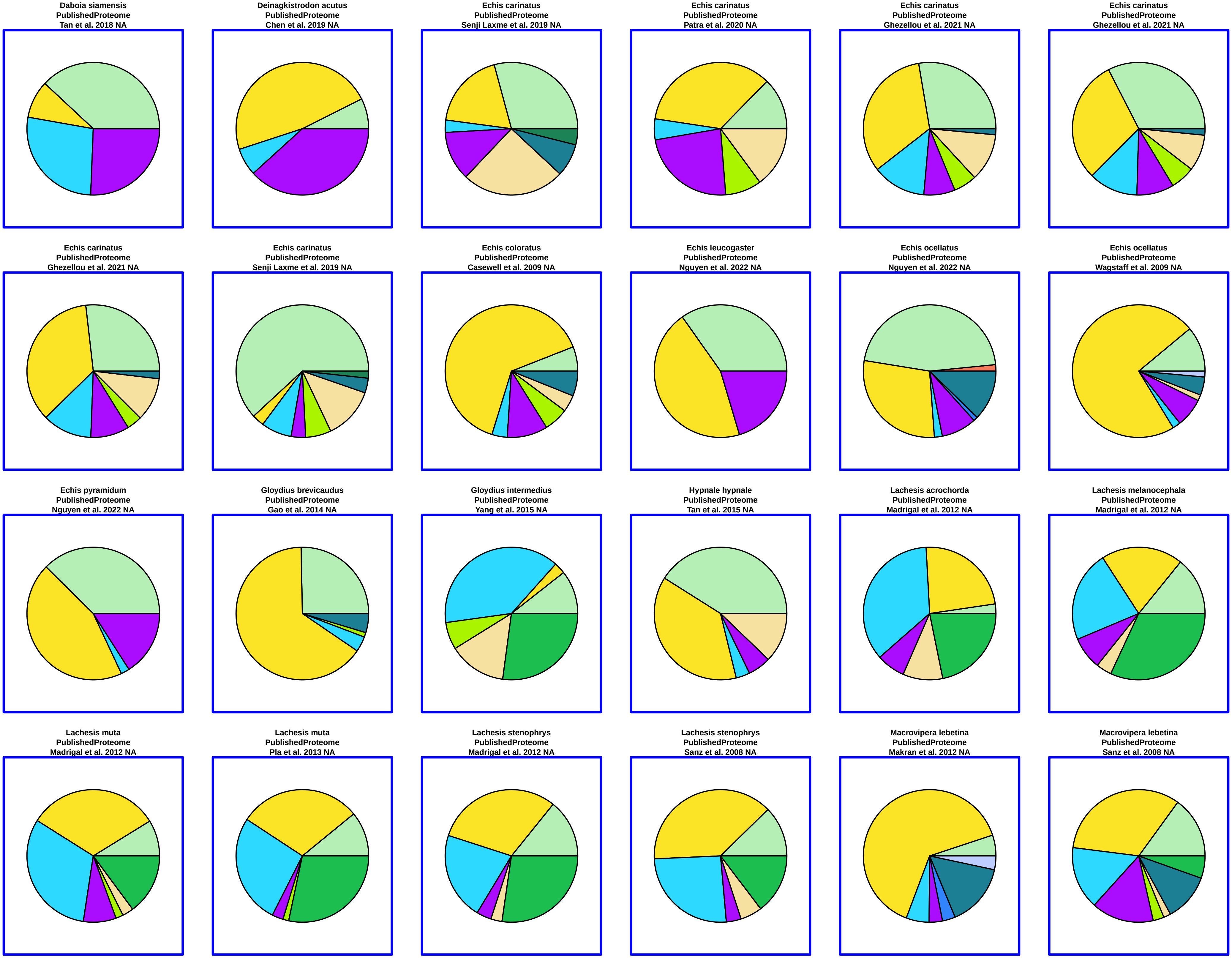
Daboia siamensis
PublishedProteome
Tan et al. 2018 NA
Deinagkistrodon acutus
PublishedProteome
Chen et al. 2019 NA
Echis carinatus
PublishedProteome
Senji Laxme et al. 2019 NA
Echis carinatus
PublishedProteome
Patra et al. 2020 NA
Echis carinatus
PublishedProteome
Ghezellou et al. 2021 NA
Echis carinatus
PublishedProteome
Ghezellou et al. 2021 NA
Echis carinatus
PublishedProteome
Ghezellou et al. 2021 NA
Echis carinatus
PublishedProteome
Senji Laxme et al. 2019 NA
Echis coloratus
PublishedProteome
Casewell et al. 2009 NA
Echis leucogaster
PublishedProteome
Nguyen et al. 2022 NA
Echis ocellatus
PublishedProteome
Nguyen et al. 2022 NA
Echis ocellatus
PublishedProteome
Wagstaff et al. 2009 NA
Echis pyramidum
PublishedProteome
Nguyen et al. 2022 NA
Gloydius brevicaudus
PublishedProteome
Gao et al. 2014 NA
Gloydius intermedius
PublishedProteome
Yang et al. 2015 NA
Hypnale hypnale
PublishedProteome
Tan et al. 2015 NA
Lachesis acrochorda
PublishedProteome
Madrigal et al. 2012 NA
Lachesis melanocephala
PublishedProteome
Madrigal et al. 2012 NA
Lachesis muta
PublishedProteome
Madrigal et al. 2012 NA
Lachesis muta
PublishedProteome
Pla et al. 2013 NA
Lachesis stenophrys
PublishedProteome
Madrigal et al. 2012 NA
Lachesis stenophrys
PublishedProteome
Sanz et al. 2008 NA
Macrovipera lebetina
PublishedProteome
Makran et al. 2012 NA
Macrovipera lebetina
PublishedProteome
Sanz et al. 2008 NA

Macrovipera lebetina
PublishedProteome
Bazaa et al. 2005 NA

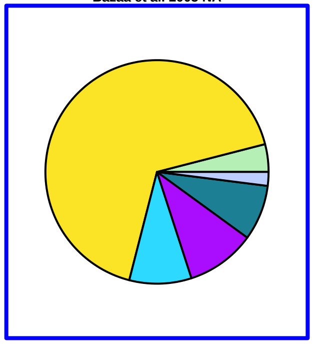

Macrovipera mauritanica
PublishedProteome
Makran et al. 2012 NA

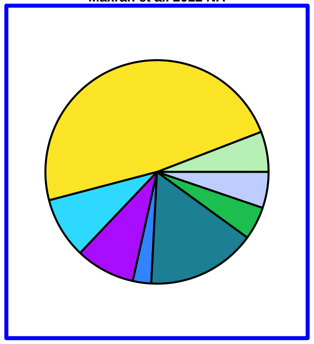

Metlapilcoatlus nummifer
PublishedProteome
Angulo et al. 2007 NA

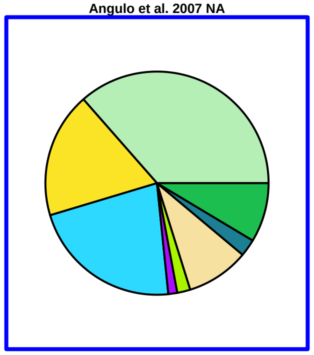

Metlapilcoatlus nummifer
PublishedProteome
Garcia−Osorio et al. 2020 NA

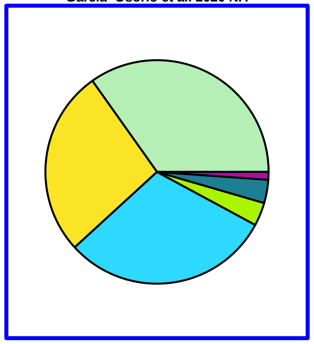

Mixcoatlus melanurus
PublishedProteome
Neri−Castro et al. 2020 NA

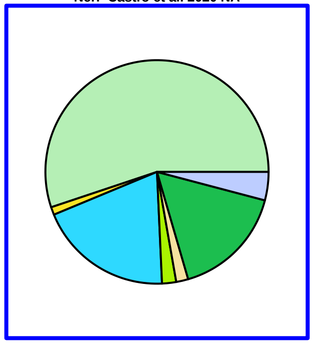

Montivipera bulgardaghica
PublishedProteome
Nalbantsoy et al. 2017 NA

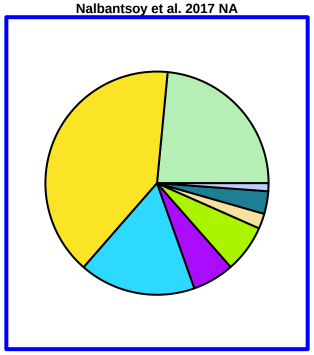

Montivipera raddei
PublishedProteome
Nalbantsoy et al. 2017 NA

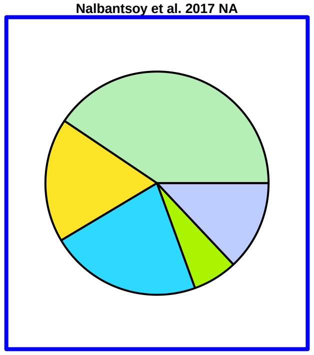

Montivipera raddei
PublishedProteome
Sanz et al. 2008 NA

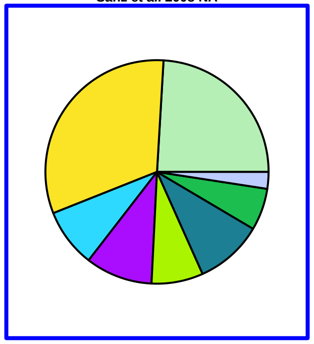

Ophryacus sphenophrys
PublishedProteome
Neri−Castro et al. 2019 NA

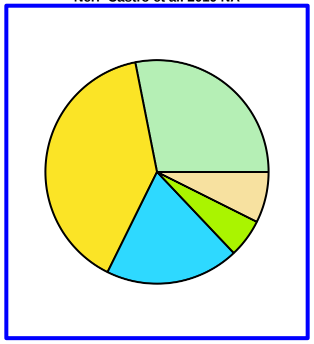

Ovophis convictus
PublishedProteome
Tan et al. 2021 NA

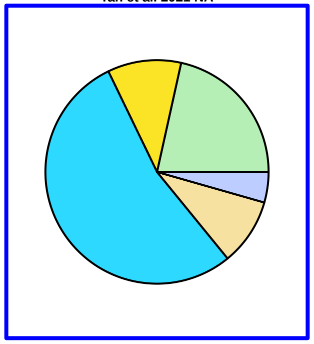

Ovophis monticola
PublishedProteome
Sitprija et al. 2021 NA

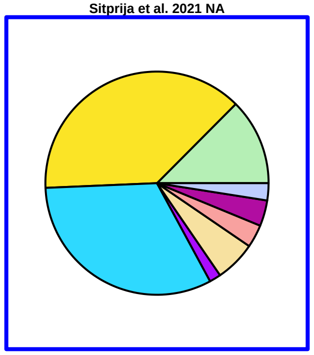

Ovophis okinavensis
PublishedProteome
Aird et al. 2013 NA

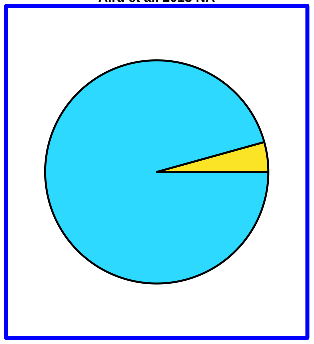

Ovophis okinavensis
PublishedProteome
Tan et al. 2021 NA

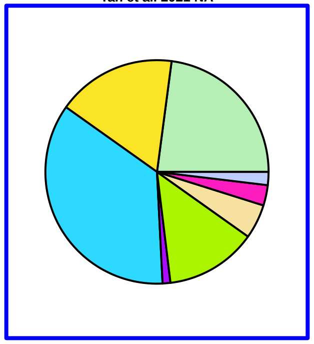

Ovophis tonkinensis
PublishedProteome
Tan et al. 2021 NA

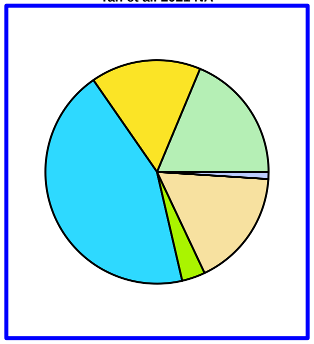

Ovophis tonkinensis
PublishedProteome
Tan et al. 2021 NA

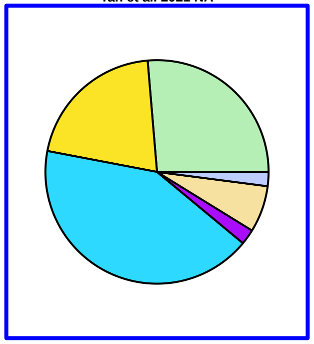

Photobothrops mucrosquamatus
PublishedProteome
Liu et al. 2021 NA

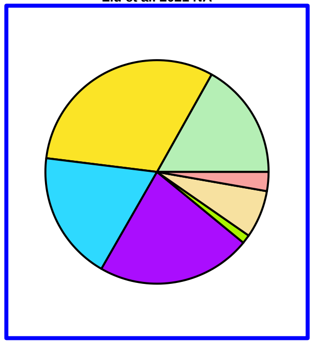

Porthidium arcosae
PublishedProteome
Ruiz−Campos et al. 2021 NA

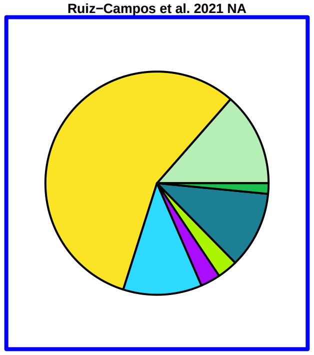

Porthidium landsbergii
PublishedProteome
Jimenez−Charris et al. 2015 NA

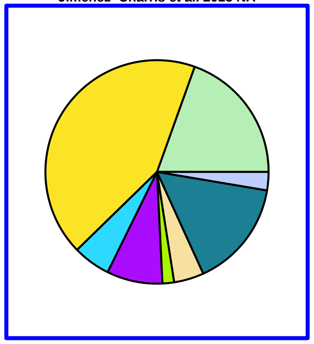

Porthidium nasutum
PublishedProteome
Lomonte et al. 2012 NA

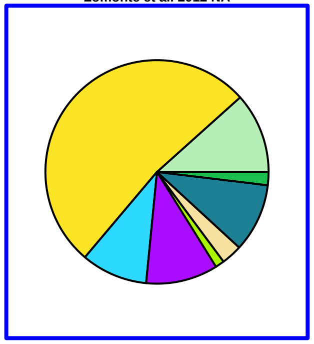

Porthidium ophryomegus
PublishedProteome
Lomonte et al. 2012 NA

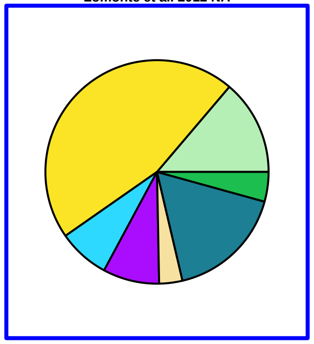

Porthidium porrasi
PublishedProteome
Mendez et al. 2019 NA

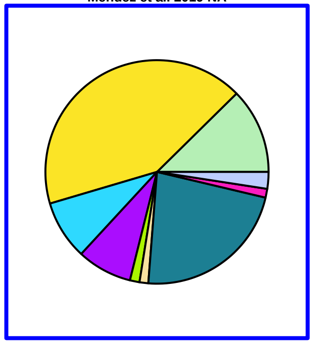

Porthidium volcanicum
PublishedProteome
Ruiz−Campos et al. 2021 NA

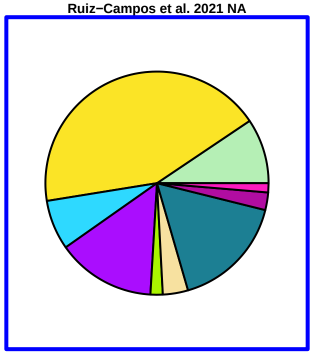

Protobothrops elegans
PublishedProteome
Tasoulis points to Aird paper which is not it NA

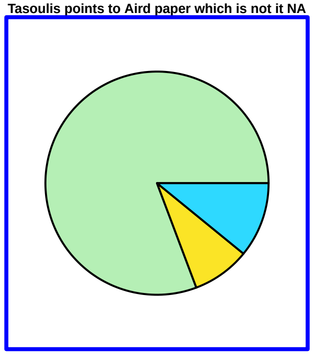

Protobothrops flavoviridis
PublishedProteome
Damm et al. 2018 NA

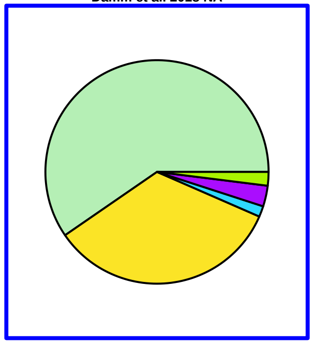

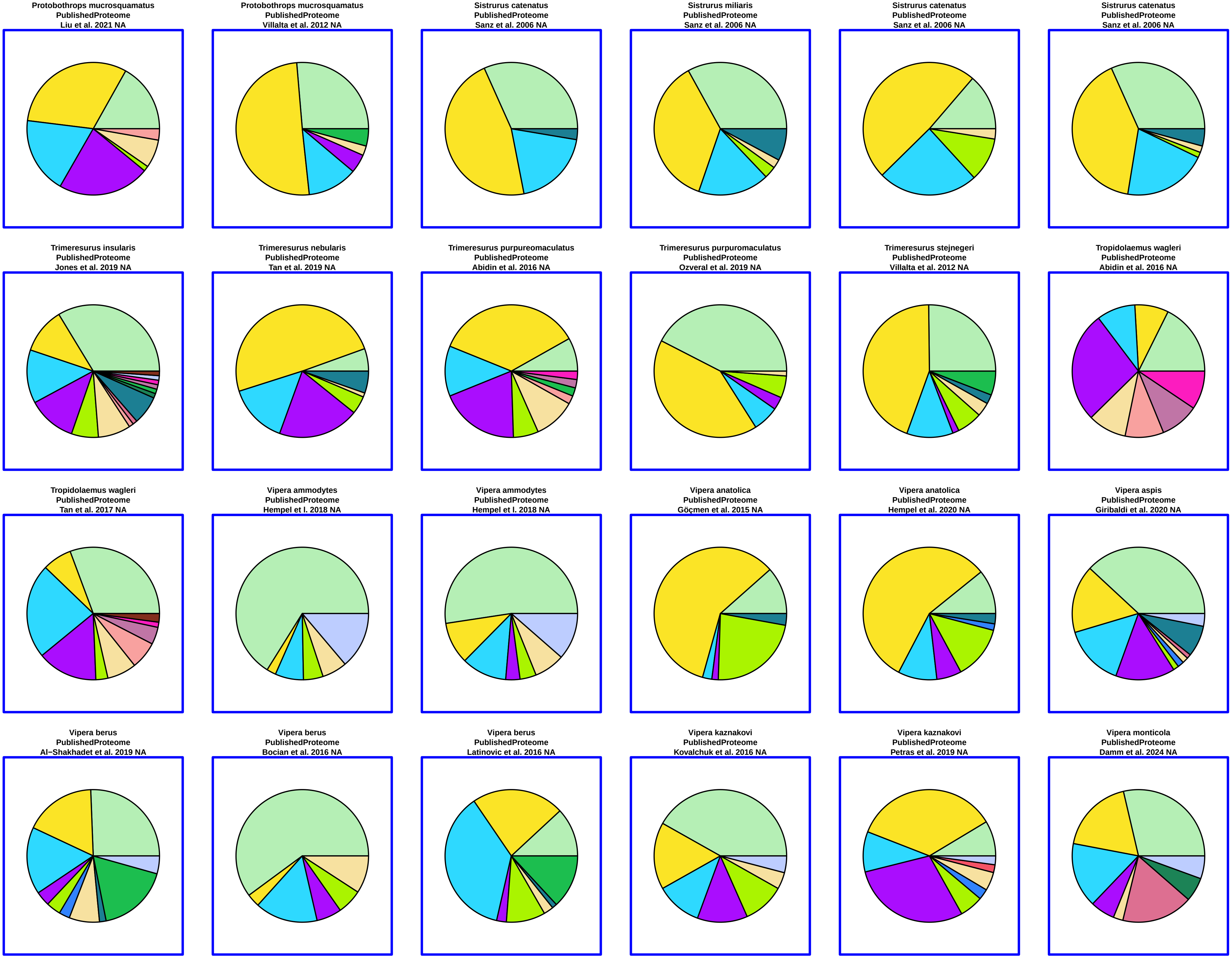
Protobothrops mucrosquamatus
PublishedProteome
Liu et al. 2021 NA
Protobothrops mucrosquamatus
PublishedProteome
Villalta et al. 2012 NA
Sistrurus catenatus
PublishedProteome
Sanz et al. 2006 NA
Sistrurus miliaris
PublishedProteome
Sanz et al. 2006 NA
Sistrurus catenatus
PublishedProteome
Sanz et al. 2006 NA
Sistrurus catenatus
PublishedProteome
Sanz et al. 2006 NA
Trimeresurus insularis
PublishedProteome
Jones et al. 2019 NA
Trimeresurus nebularis
PublishedProteome
Tan et al. 2019 NA
Trimeresurus purpureomaculatus
PublishedProteome
Abidin et al. 2016 NA
Trimeresurus purpuromaculatus
PublishedProteome
Ozveral et al. 2019 NA
Trimeresurus stejnegeri
PublishedProteome
Villalta et al. 2012 NA
Tropidolaemus wagleri
PublishedProteome
Abidin et al. 2016 NA
Tropidolaemus wagleri
PublishedProteome
Tan et al. 2017 NA
Vipera ammodytes
PublishedProteome
Hempel et l. 2018 NA
Vipera ammodytes
PublishedProteome
Hempel et l. 2018 NA
Vipera anatolica
PublishedProteome
Göçmen et al. 2015 NA
Vipera anatolica
PublishedProteome
Hempel et al. 2020 NA
Vipera aspis
PublishedProteome
Giribaldi et al. 2020 NA
Vipera berus
PublishedProteome
Al−Shakhadet et al. 2019 NA
Vipera berus
PublishedProteome
Bocian et al. 2016 NA
Vipera berus
PublishedProteome
Latinovic et al. 2016 NA
Vipera kaznakovi
PublishedProteome
Kovalchuk et al. 2016 NA
Vipera kaznakovi
PublishedProteome
Petras et al. 2019 NA
Vipera monticola
PublishedProteome
Damm et al. 2024 NA

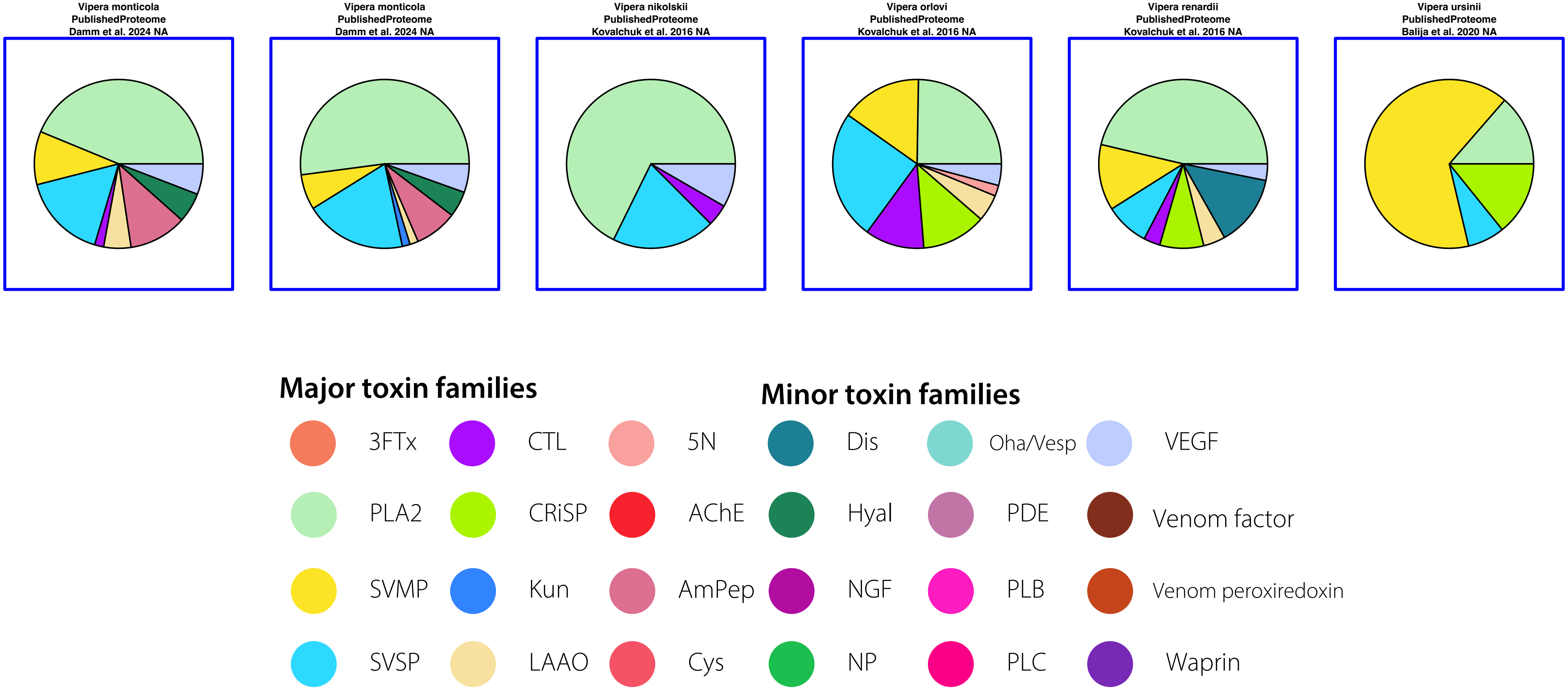

Vipera monticola
PublishedProteome
Damm et al. 2024 NA
Vipera monticola
PublishedProteome
Damm et al. 2024 NA
Vipera nikolskii
PublishedProteome
Kovalchuk et al. 2016 NA
Vipera orlovi
PublishedProteome
Kovalchuk et al. 2016 NA
Vipera renardii
PublishedProteome
Kovalchuk et al. 2016 NA
Vipera ursinii
PublishedProteome
Balija et al. 2020 NA
Major toxin families
Minor toxin families
3FTx
CTL
5N
Dis
Oha/Vesp
VEGF
PLA2
CRiSP
AChE
Hyal
PDE
Venom factor
SVMP
Kun
AmPep
NGF
PLB
Venom peroxiredoxin
SVSP
LAAO
Cys
NP
PLC
Waprin

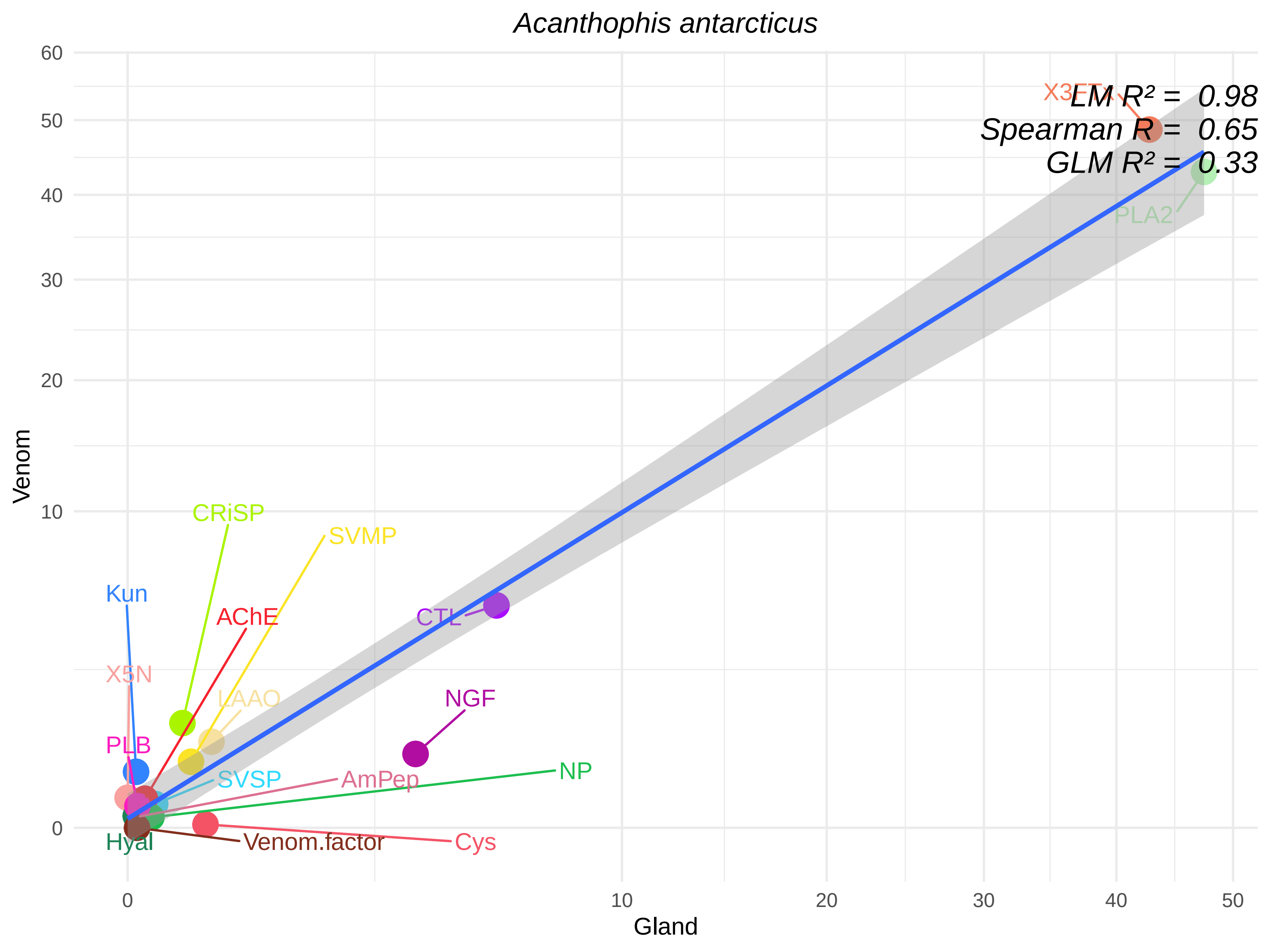

Acanthophis antarcticus
LM R² = 0.98
Spearman R = 0.65
GLM R² = 0.33
X3FTx
PLA2
CTL
NGF
NP
CRiSP
SVMP
Kun
AChE
X5N
LAAO
PLB
SVSP
AmPep
Hyal
Venom.factor
Cys
Venom
Gland
60
50
40
30
20
10
0
0
10
20
30
40
50

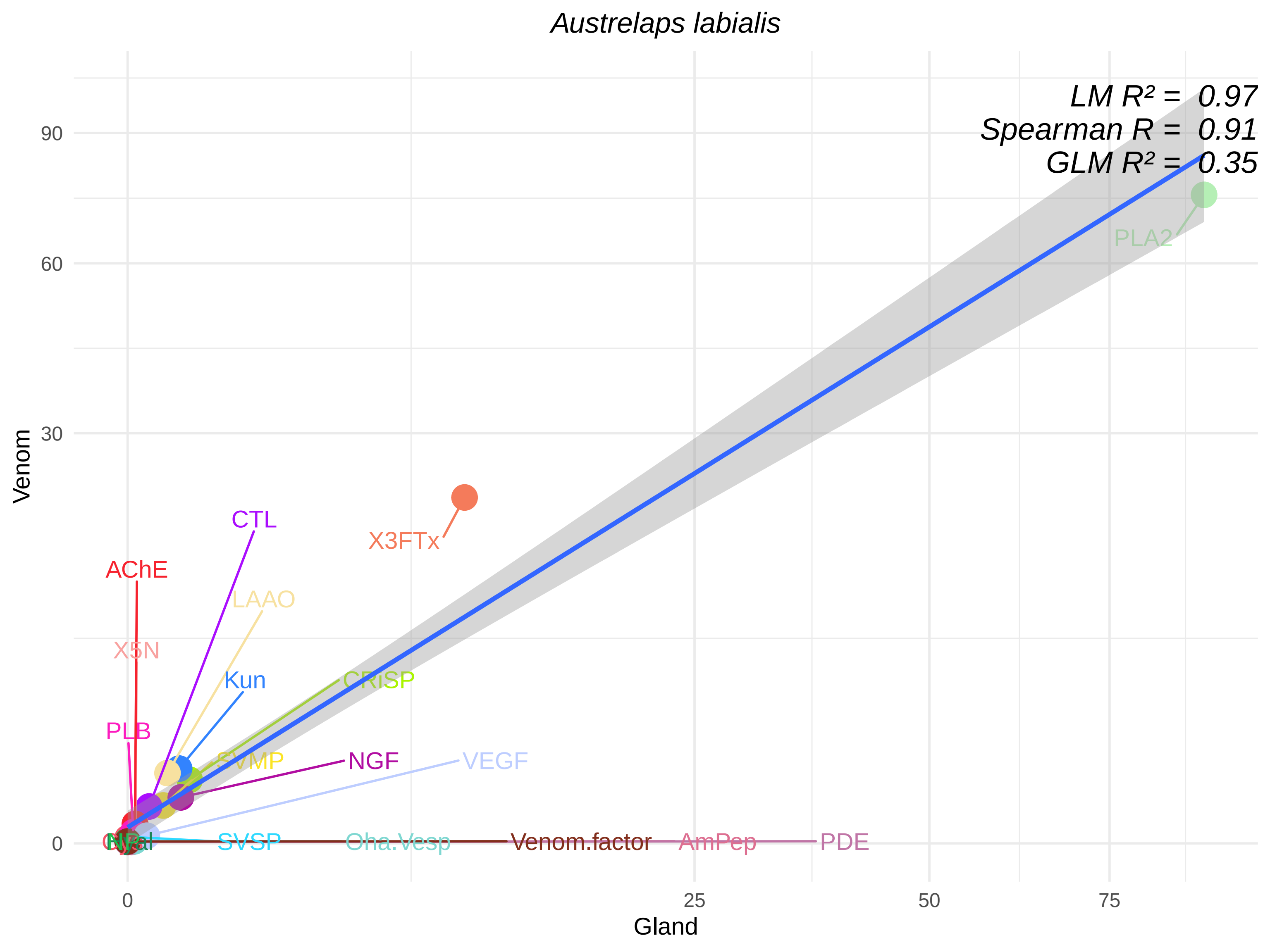

Austrelaps labialis
LM R² = 0.97
Spearman R = 0.91
GLM R² = 0.35
Venom
Gland
0
30
60
90
0
25
50
75
PLA2
X3FTx
CTL
AChE
LAAO
X5N
Kun
CRiSP
PLB
SVMP
NGF
VEGF
SVSP
Oha.Vesp
Venom.factor
AmPep
PDE

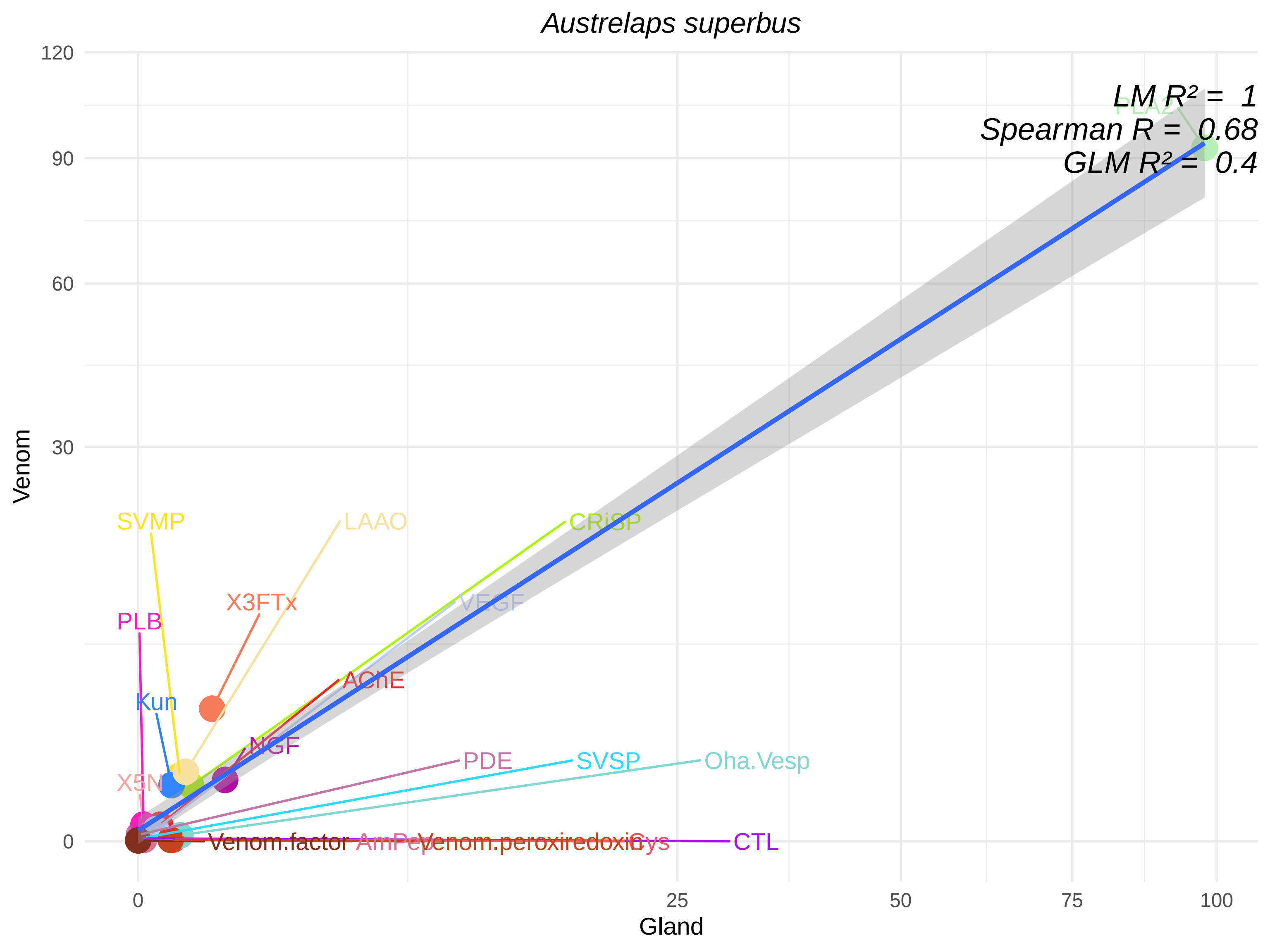
Austrelaps superbus
LM R² = 1
Spearman R = 0.68
GLM R² = 0.4
PLA2
SVMP
LAAO
CRiSP
X3FTx
VEGF
PLB
AChE
Kun
NGF
PDE
SVSP
Oha.Vesp
X5N
Venom.factor
AmPep
Venom.peroxiredoxin
Cys
CTL
Venom
Gland
0
30
60
90
120
25
50
75
100

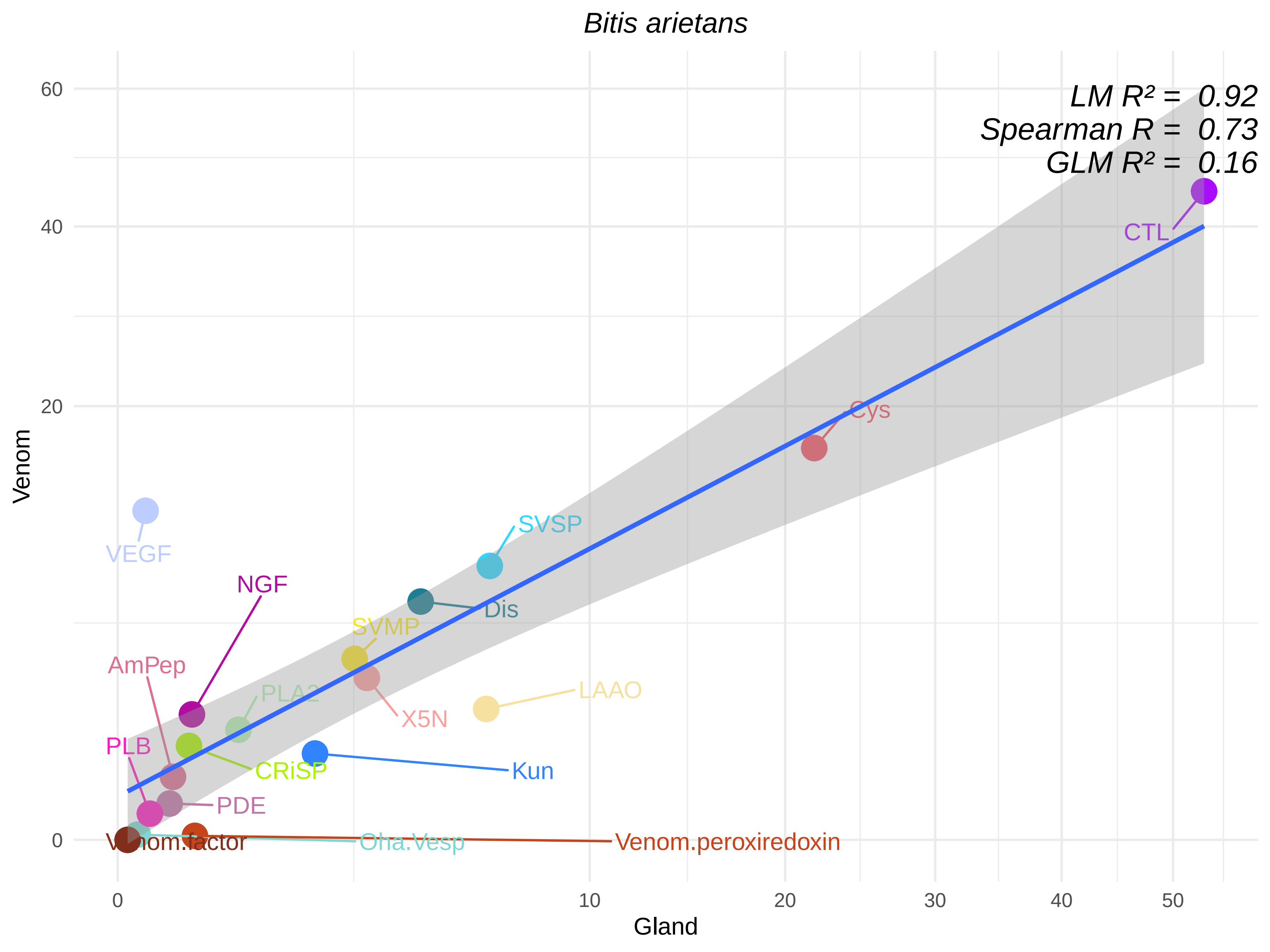

Bitis arietans
LM R² = 0.92
Spearman R = 0.73
GLM R² = 0.16
Venom
Gland
0
20
40
60
0
10
20
30
40
50
CTL
Cys
SVSP
Dis
VEGF
NGF
SVMP
AmPep
PLA2
X5N
LAAO
PLB
CRiSP
Kun
PDE
Venom.factor
Oha.Vesp
Venom.peroxiredoxin

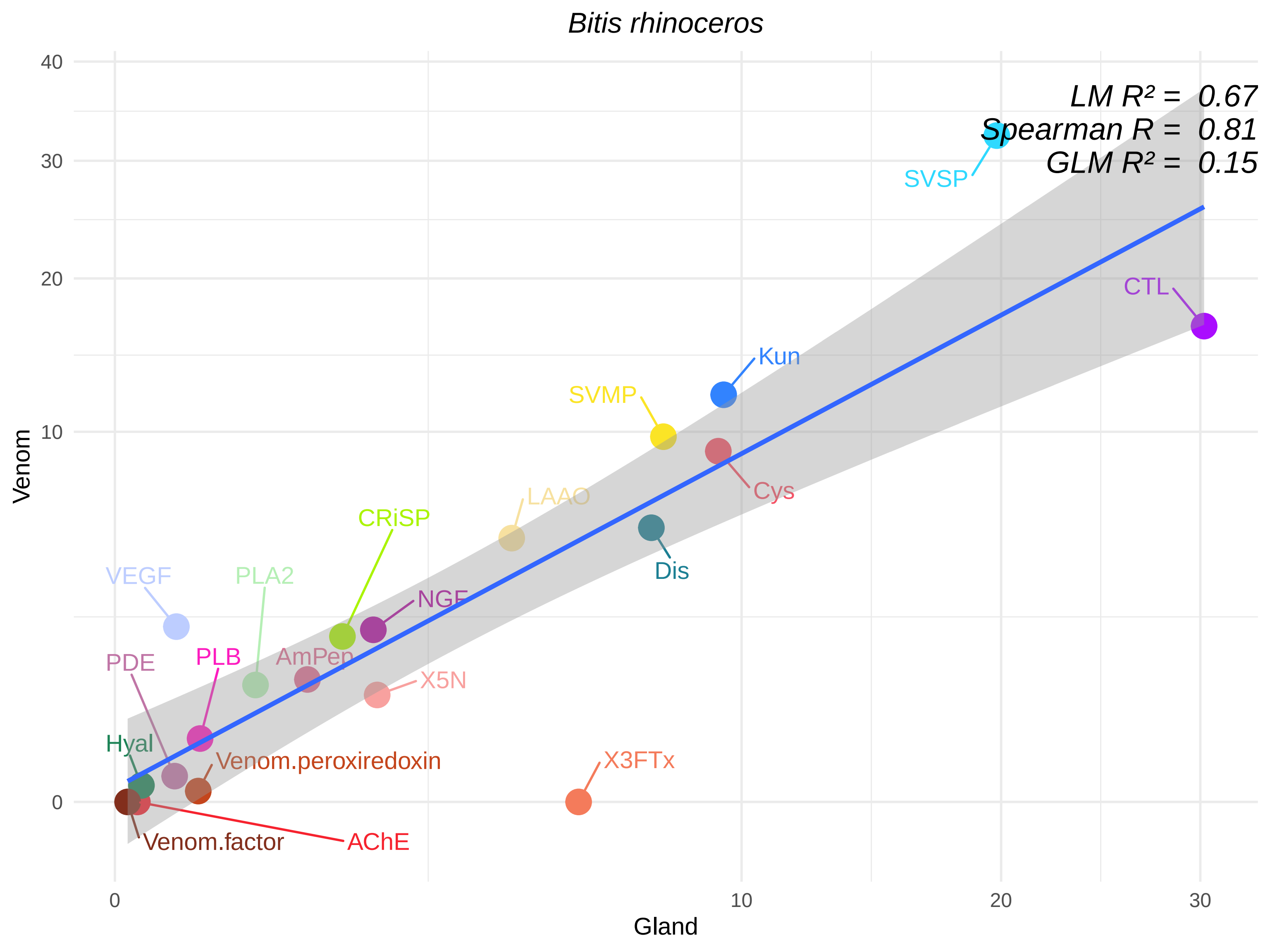

Bitis rhinoceros
LM R² = 0.67
Spearman R = 0.81
GLM R² = 0.15
SVSP
CTL
Kun
SVMP
Cys
LAAO
CRiSP
Dis
VEGF
PLA2
NGF
PDE
PLB
AmPep
X5N
Hyal
Venom.peroxiredoxin
X3FTx
Venom.factor
AChE
Venom
Gland
0
10
20
30
40

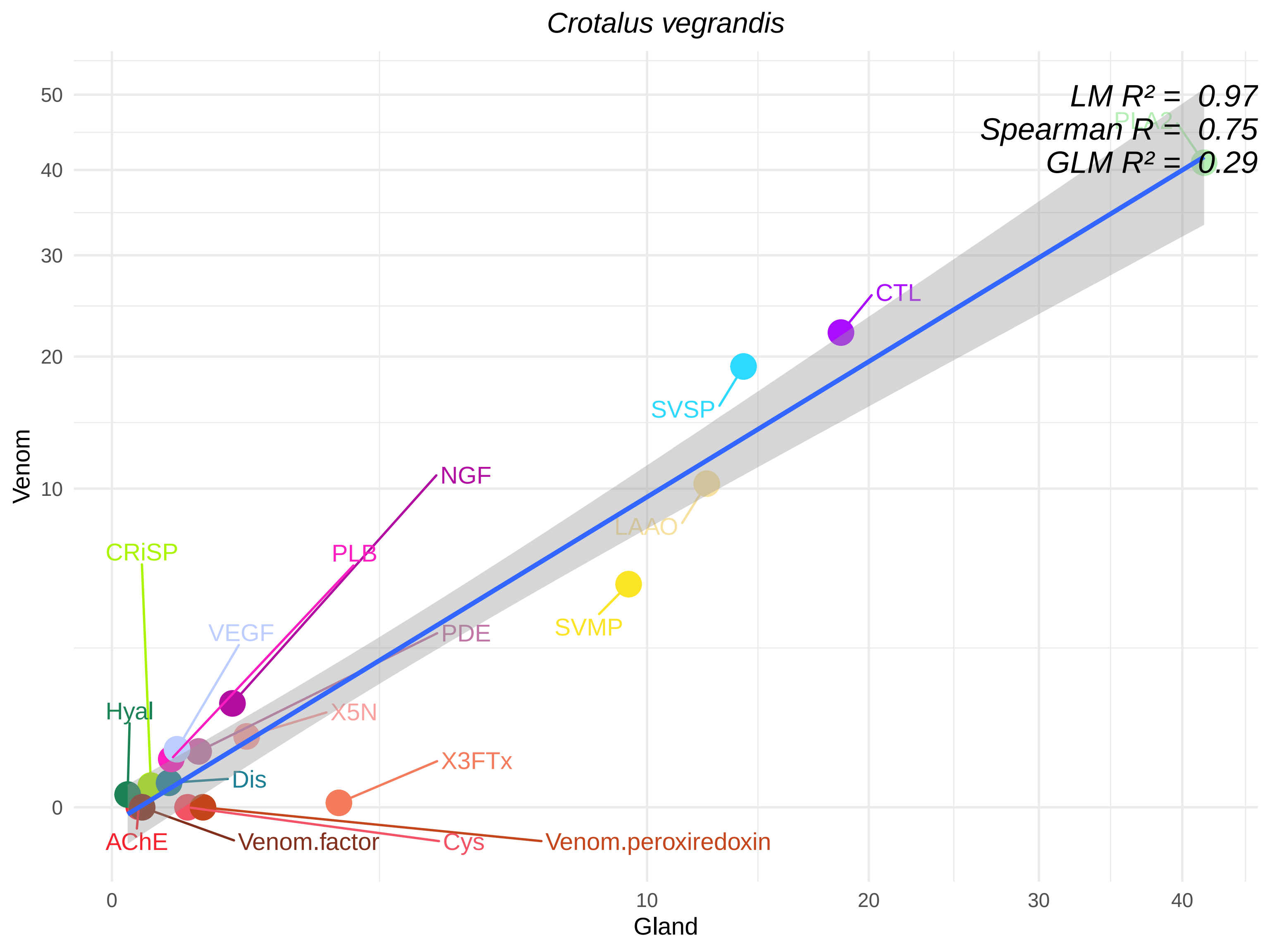
Crotalus vegrandis
LM R² = 0.97
Spearman R = 0.75
GLM R² = 0.29
PLA2
CTL
SVSP
NGF
LAAO
CRiSP
PLB
SVMP
VEGF
PDE
Hyal
X5N
X3FTx
Dis
AChE
Venom.factor
Cys
Venom.peroxiredoxin
Venom
Gland
0
10
20
30
40
50

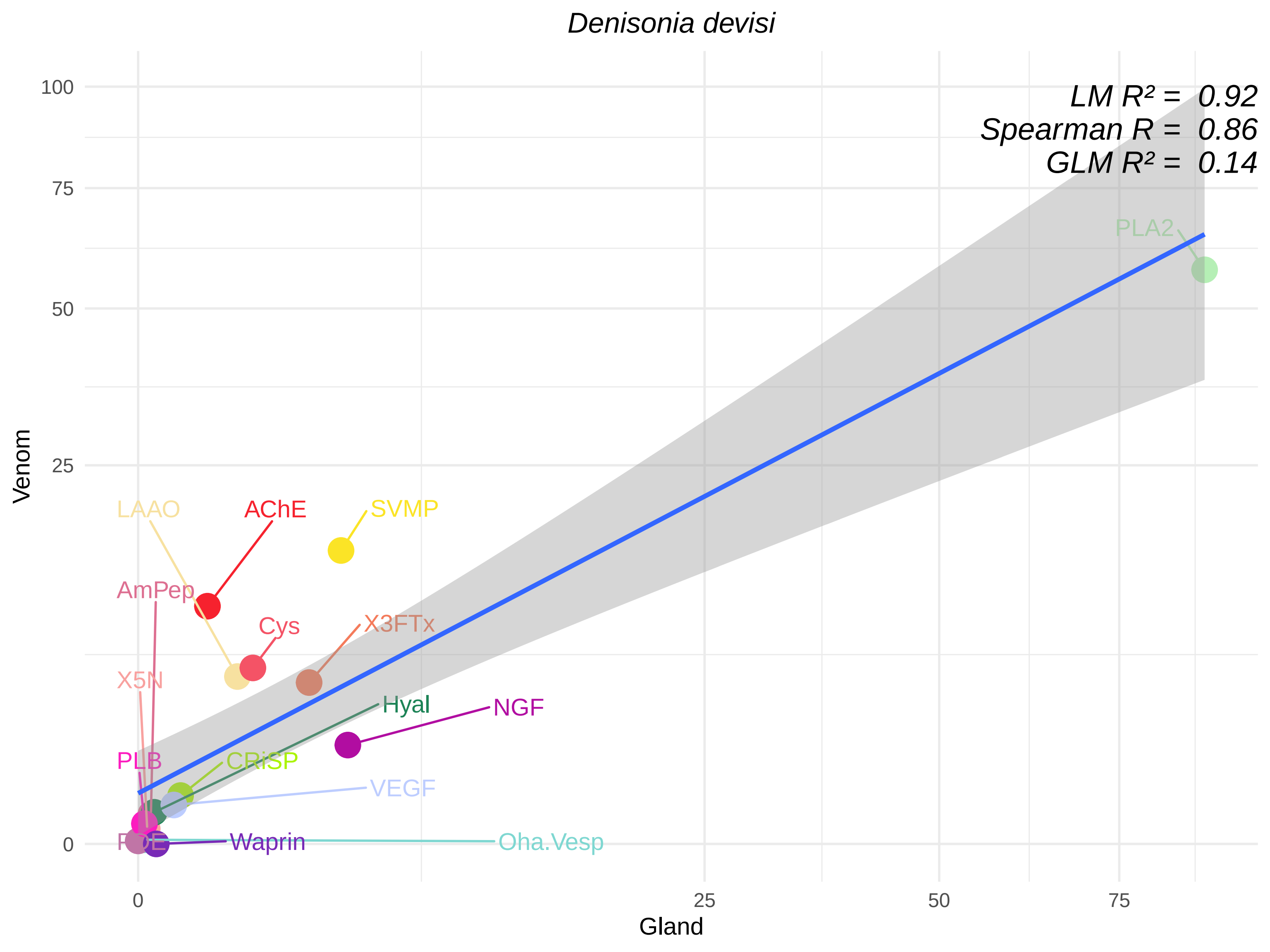
Denisonia devisi
LM R² = 0.92
Spearman R = 0.86
GLM R² = 0.14
PLA2
LAAO
AChE
SVMP
AmPep
Cys
X3FTx
X5N
Hyal
NGF
PLB
CRiSP
VEGF
PDE
Waprin
Oha.Vesp
Venom
Gland
100
75
50
25
0
0
25
50
75

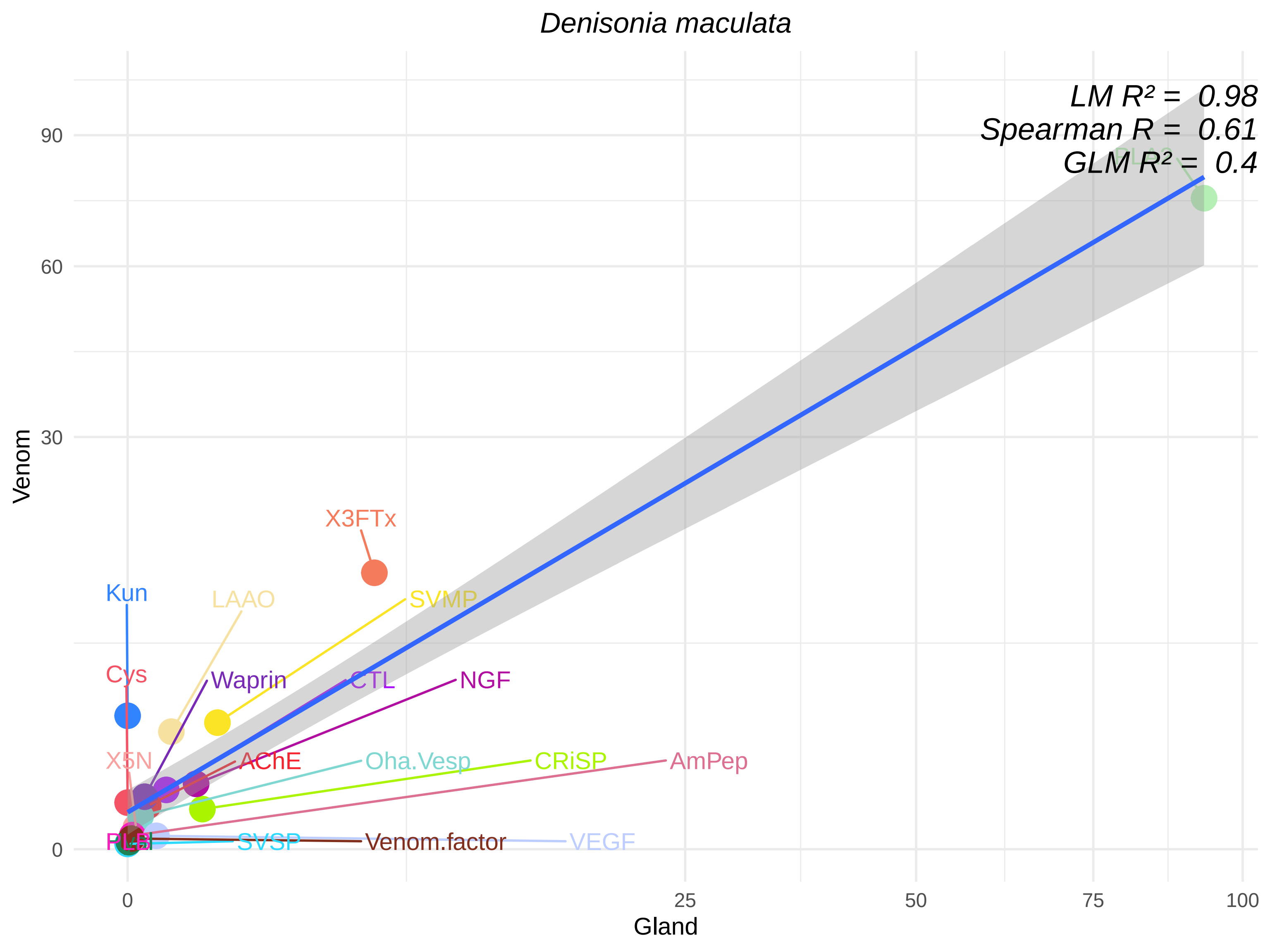

*Denisonia maculata*
LM R² = 0.98
Spearman R = 0.61
GLM R² = 0.4
X3FTx
Kun
LAAO
SVMP
Cys
Waprin
CTL
NGF
X5N
AChE
Oha.Vesp
CRiSP
AmPep
PLB
SVSP
Venom.factor
VEGF
Venom
Gland
0
30
60
90
0
25
50
75
100

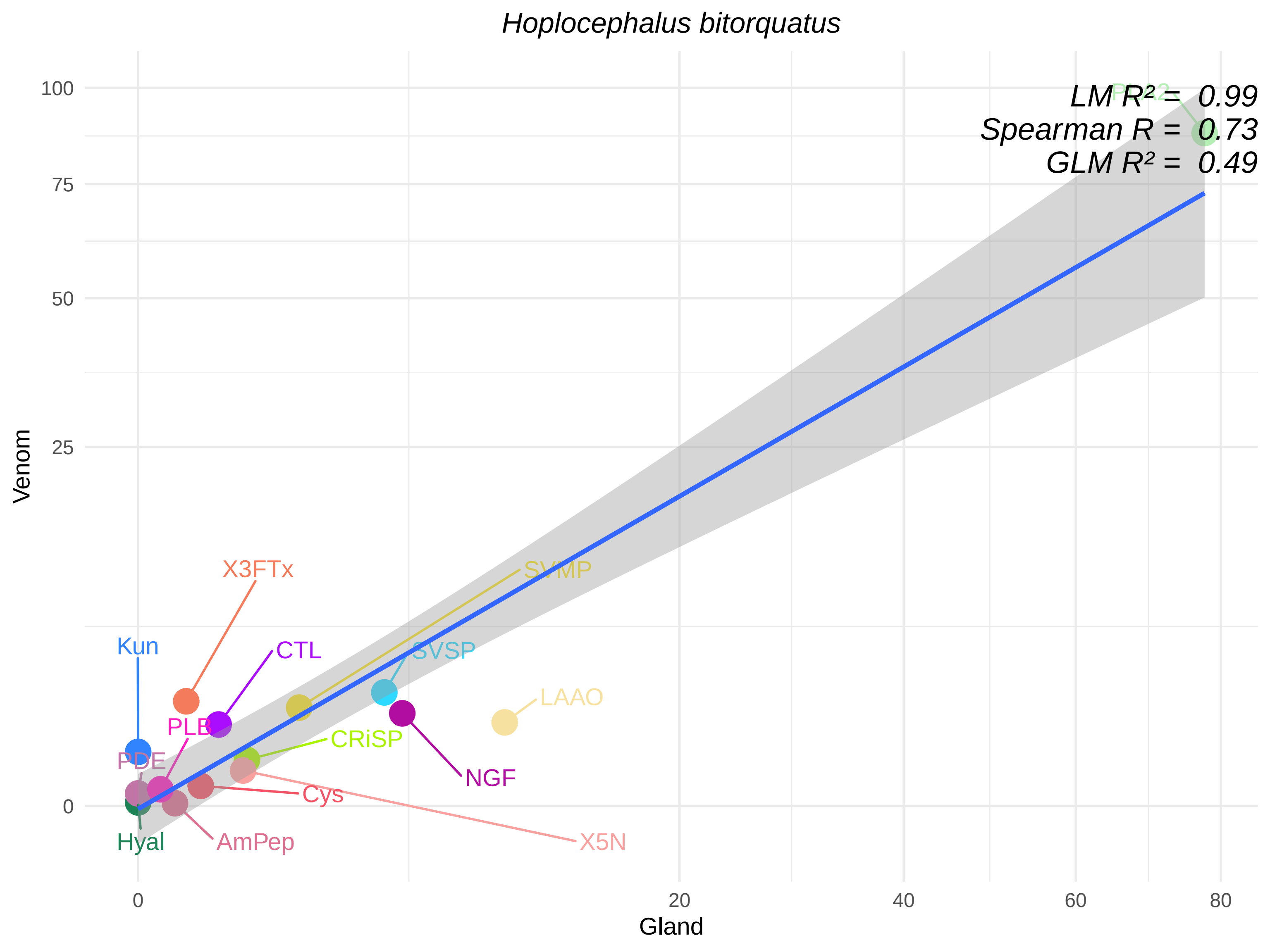

Hoplocephalus bitorquatus
LM R² = 0.99
Spearman R = 0.73
GLM R² = 0.49
PLA2
X3FTx
SVMP
Kun
CTL
SVSP
LAAO
PLB
CRiSP
PDE
NGF
Cys
Hyal
AmPep
X5N
Venom
Gland
100
75
50
25
0
20
40
60
80

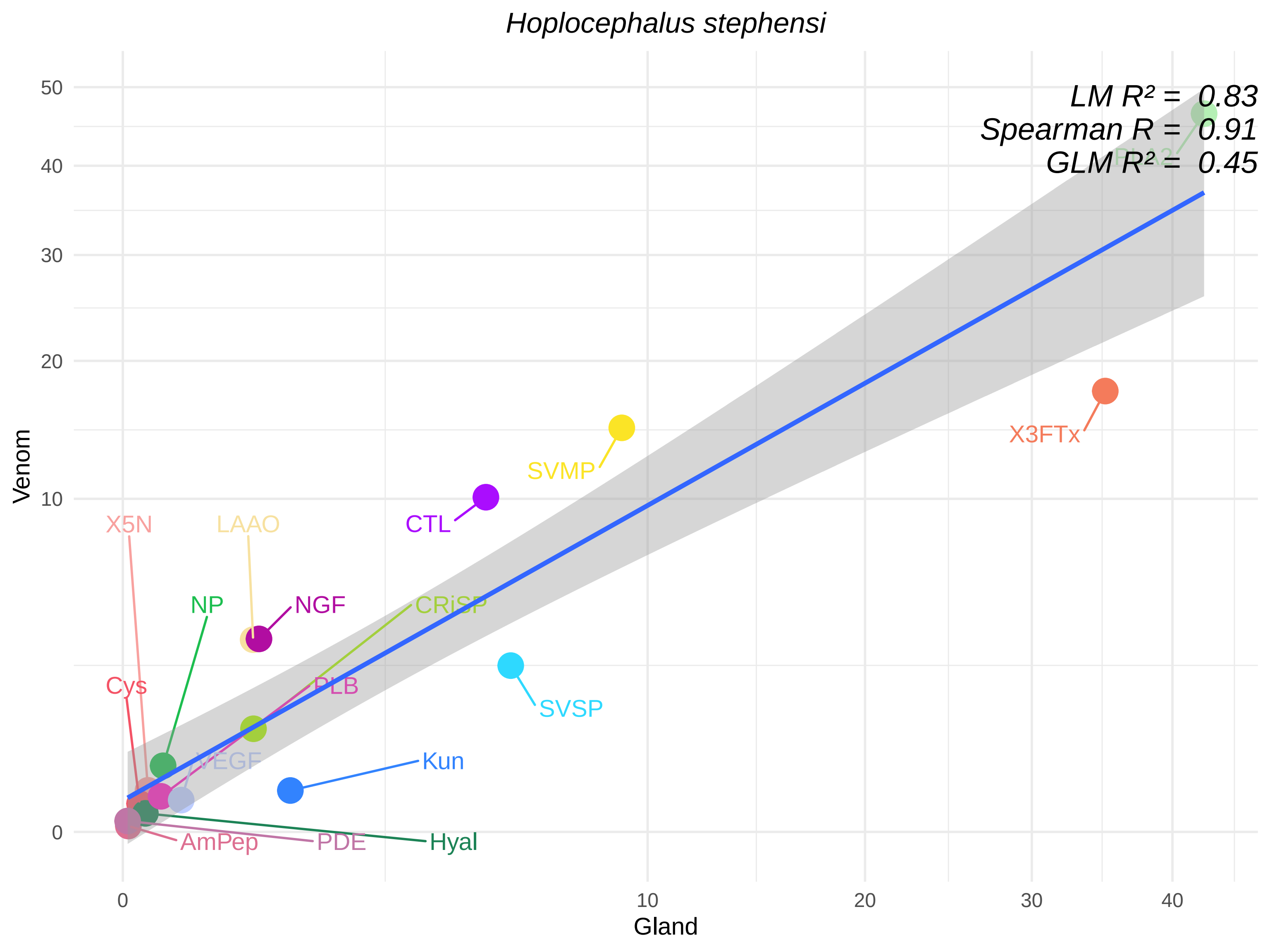

Hoplocephalus stephensi
LM R² = 0.83
Spearman R = 0.91
GLM R² = 0.45
PLA2
X3FTx
SVMP
CTL
X5N
LAAO
NP
NGF
CRiSP
SVSP
Cys
PLB
VEGF
Kun
AmPep
PDE
Hyal
Venom
Gland
50
40
30
20
10
0
0
10
20
30
40

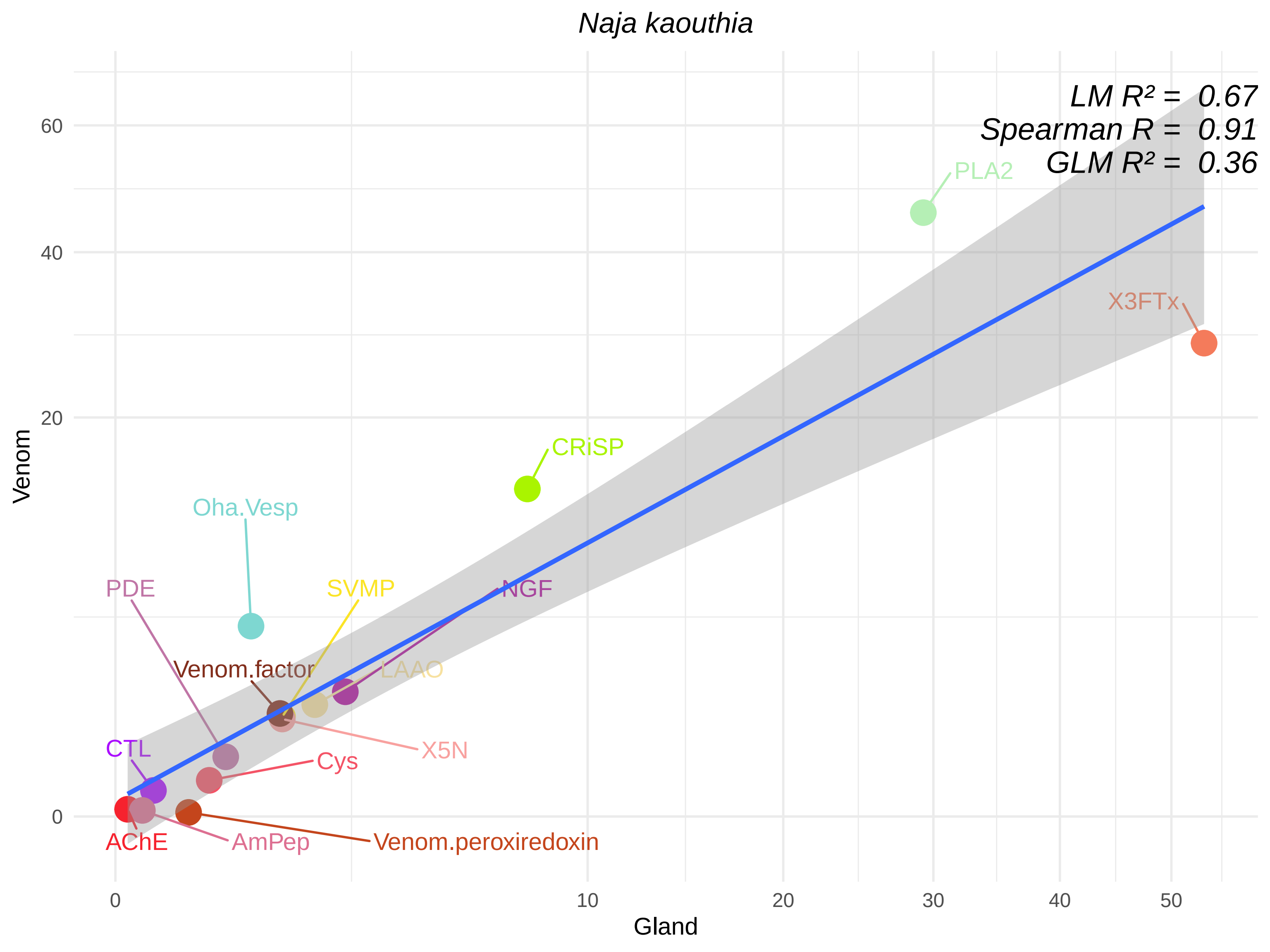
Naja kaouthia
LM R² = 0.67
Spearman R = 0.91
GLM R² = 0.36
PLA2
X3FTx
CRiSP
Oha.Vesp
PDE
SVMP
NGF
Venom.factor
LAAO
X5N
CTL
Cys
AChE
AmPep
Venom.peroxiredoxin
Venom
Gland
0
20
40
60
0
10
20
30
40
50

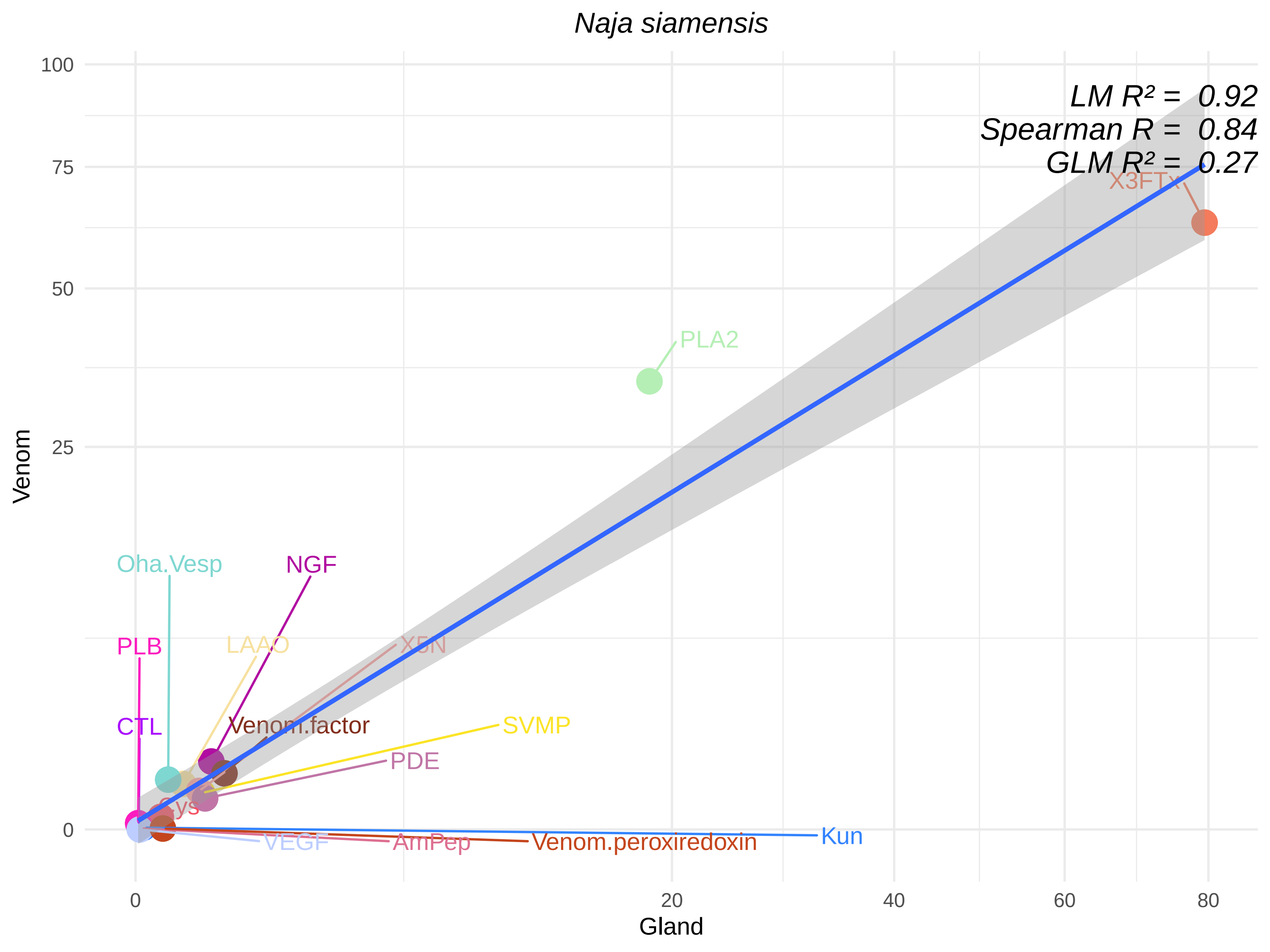

Naja siamensis
LM R² = 0.92
Spearman R = 0.84
GLM R² = 0.27
X3FTx
PLA2
Oha.Vesp
NGF
PLB
LAAO
X5N
CTL
Venom.factor
SVMP
PDE
Cys
VEGF
AmPep
Venom.peroxiredoxin
Kun
100
75
50
25
0
Venom
0
20
40
60
80
Gland

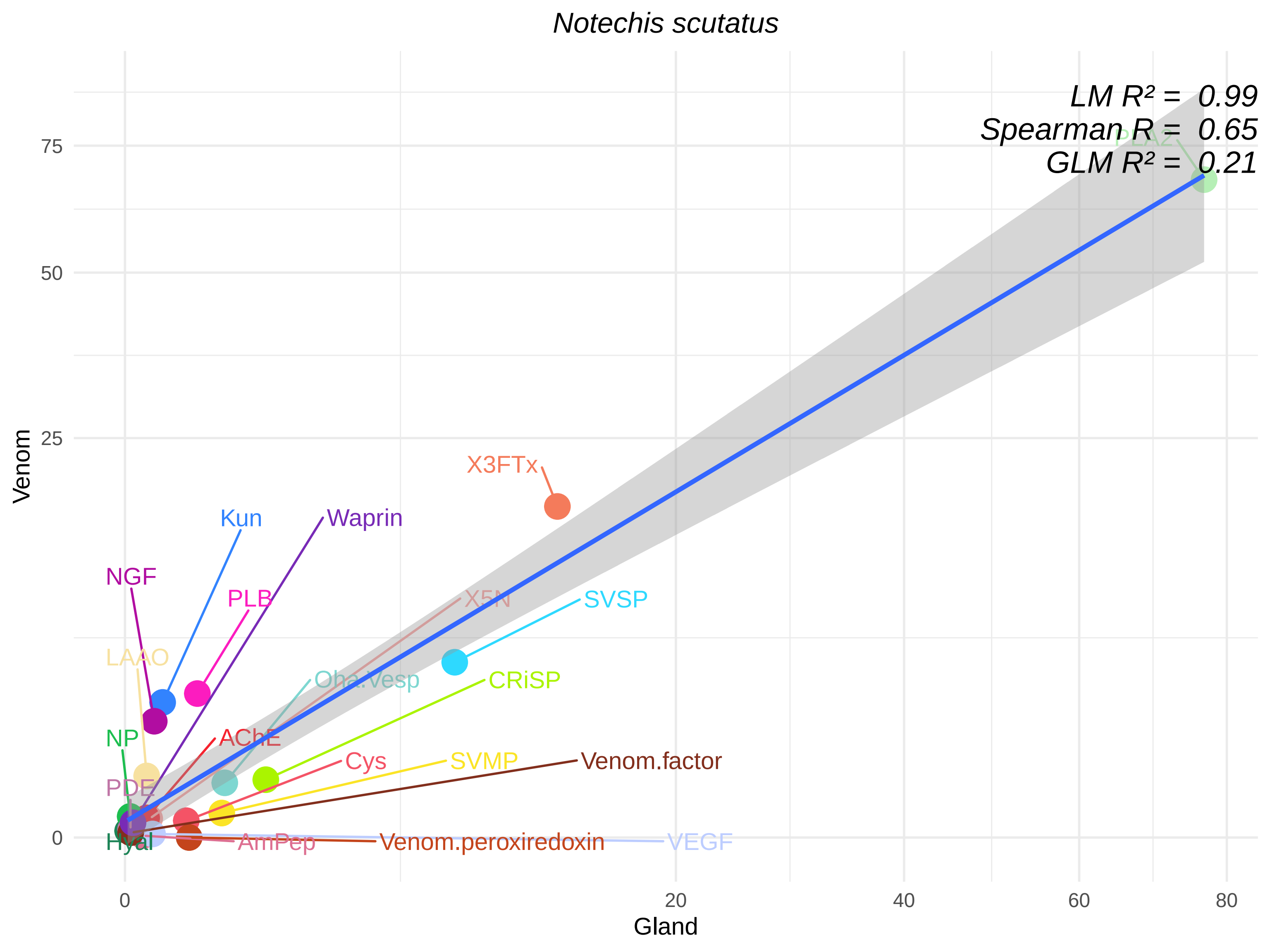

Notechis scutatus
LM R² = 0.99
Spearman R = 0.65
GLM R² = 0.21
PLA2
X3FTx
Kun
Waprin
NGF
PLB
X5N
SVSP
LAAO
Oha.Vesp
CRiSP
NP
AChE
Cys
SVMP
Venom.factor
PDE
Hyal
AmPep
Venom.peroxiredoxin
VEGF
Venom
Gland
0
25
50
75
0
20
40
60
80

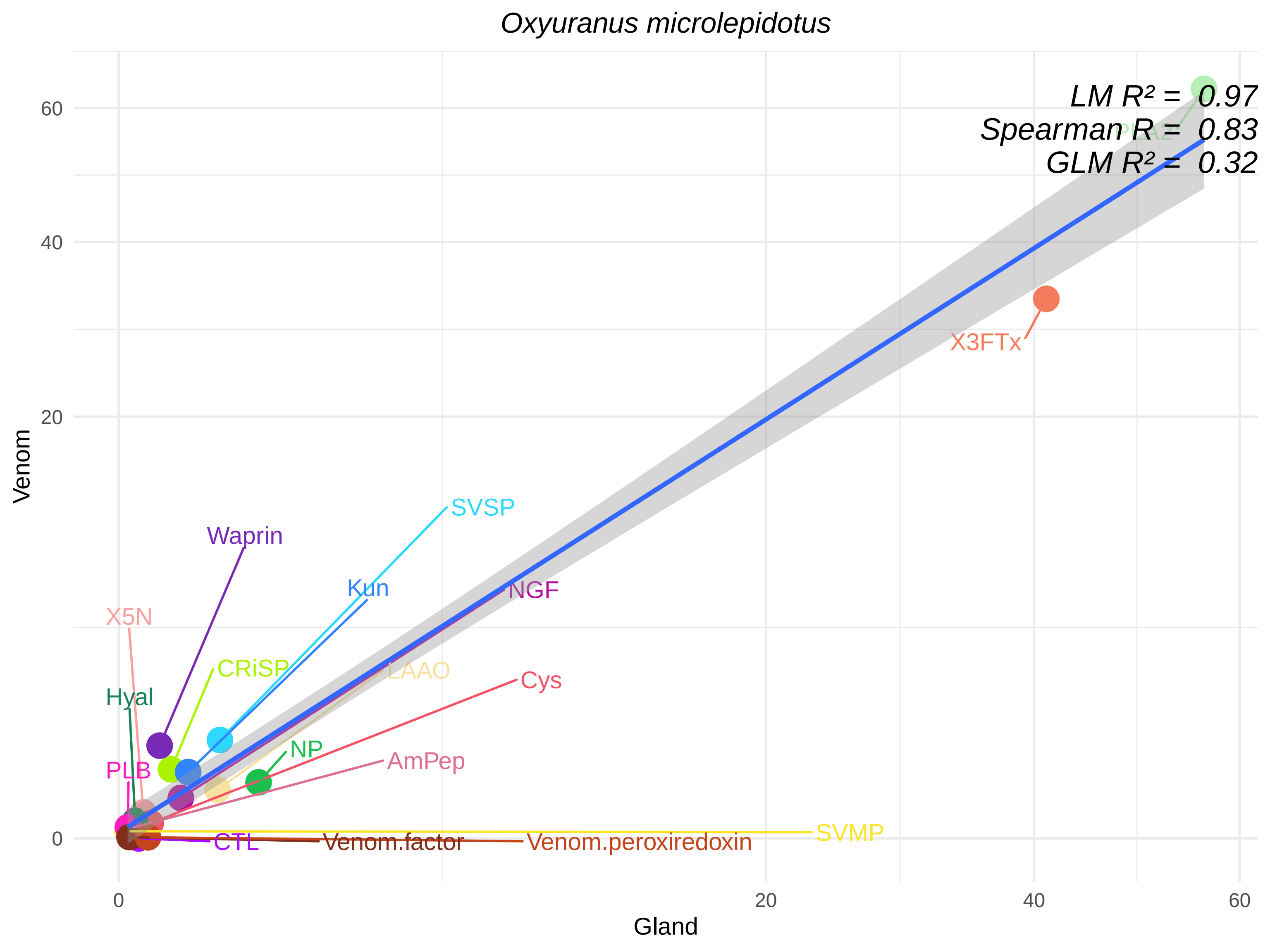

Oxyuranus microlepidotus
LM R² = 0.97
Spearman R = 0.83
GLM R² = 0.32
PLA2
X3FTx
SVSP
Waprin
Kun
NGF
X5N
CRiSP
LAAO
Cys
Hyal
NP
AmPep
PLB
SVMP
CTL
Venom.factor
Venom.peroxiredoxin
Venom
Gland
0
20
40
60

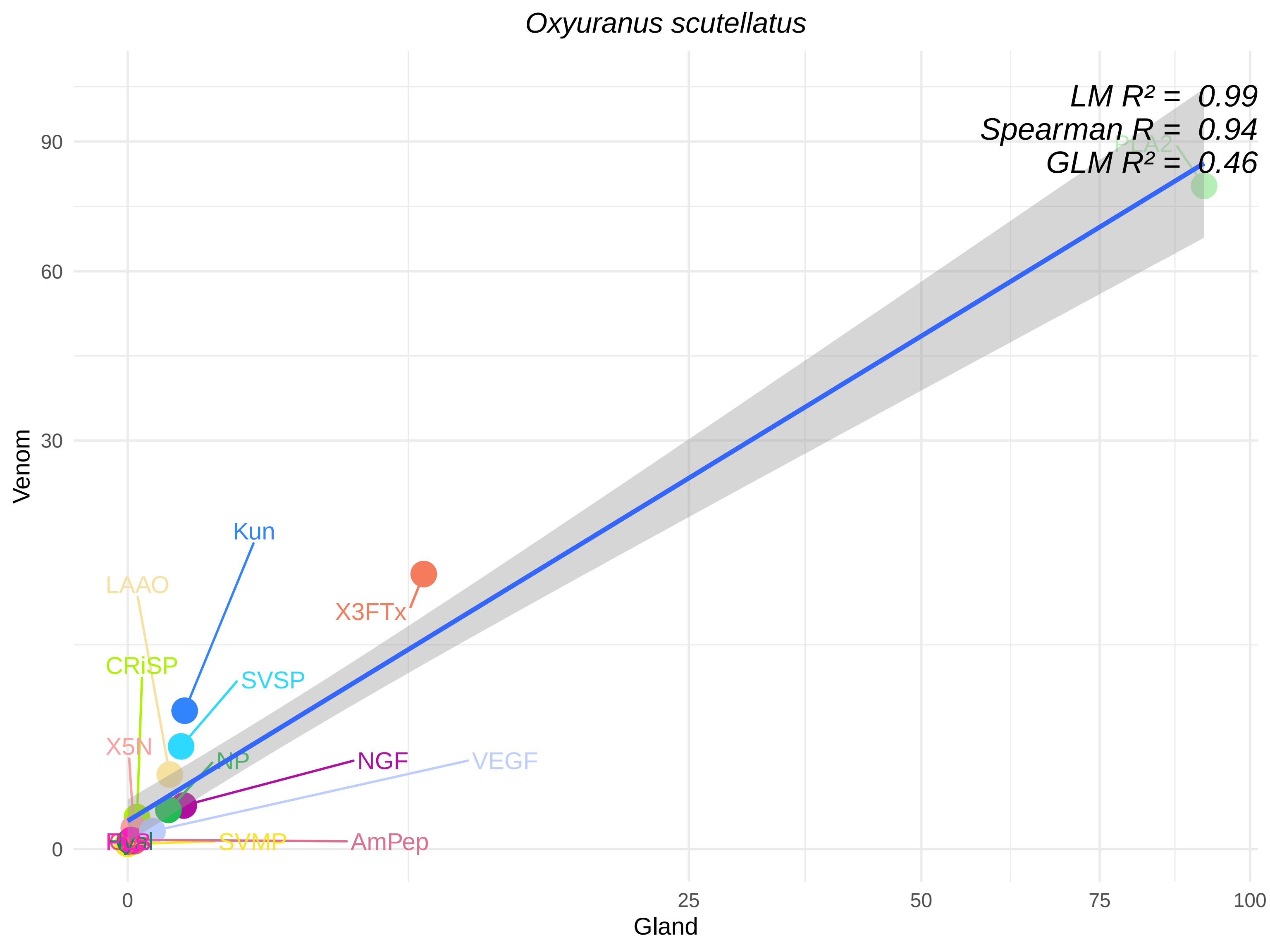

Oxyuranus scutellatus
LM R² = 0.99
Spearman R = 0.94
GLM R² = 0.46
PLA2
Kun
LAAO
X3FTx
CRiSP
SVSP
X5N
NP
NGF
VEGF
SVMP
AmPep
Venom
Gland
0
30
60
90
0
25
50
75
100

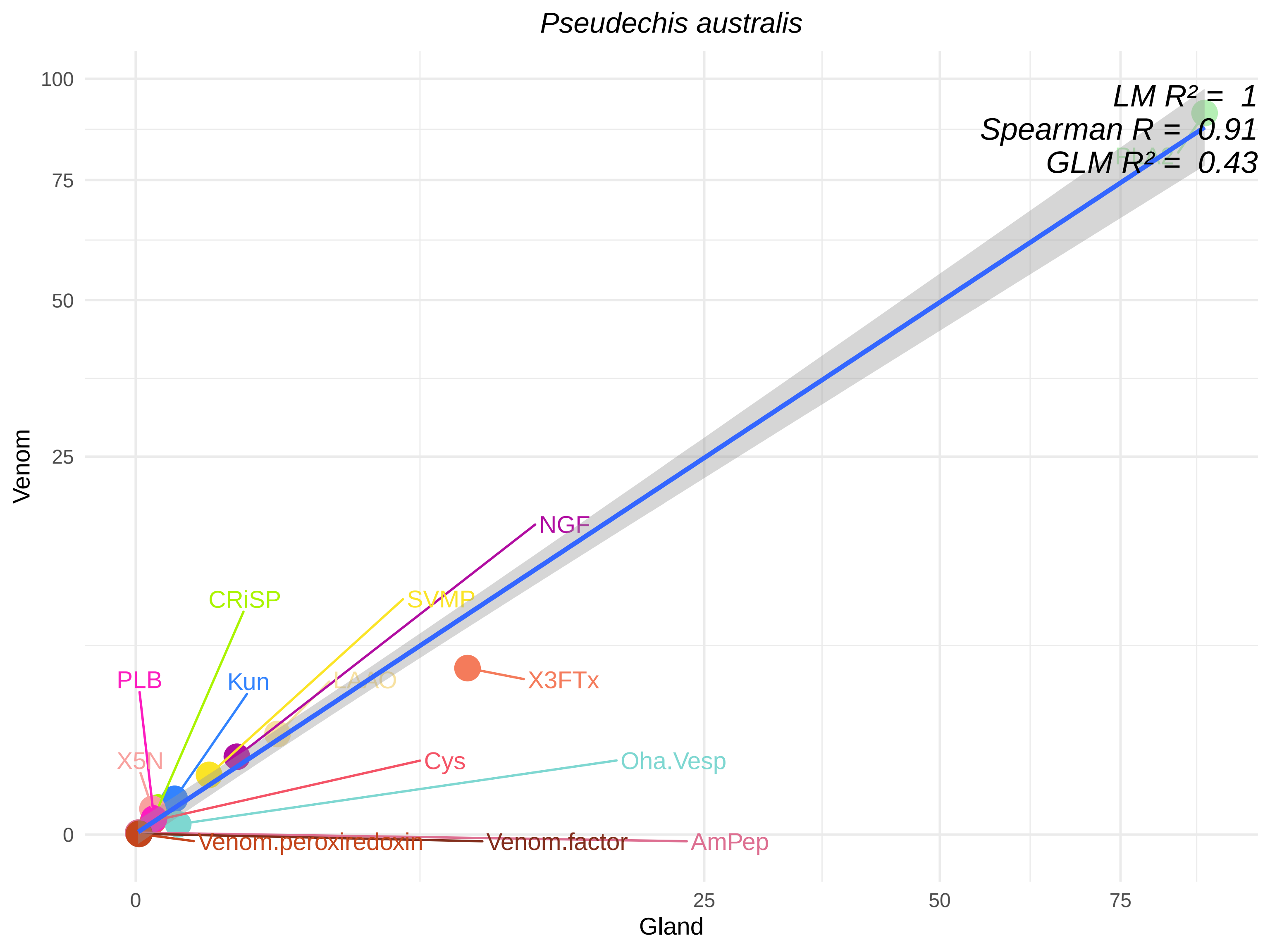

Pseudechis australis
LM R² = 1
Spearman R = 0.91
GLM R² = 0.43
NGF
CRiSP
SVMP
PLB
Kun
LAAO
X3FTx
X5N
Cys
Oha.Vesp
Venom.peroxiredoxin
Venom.factor
AmPep
Venom
Gland
0
25
50
75
100

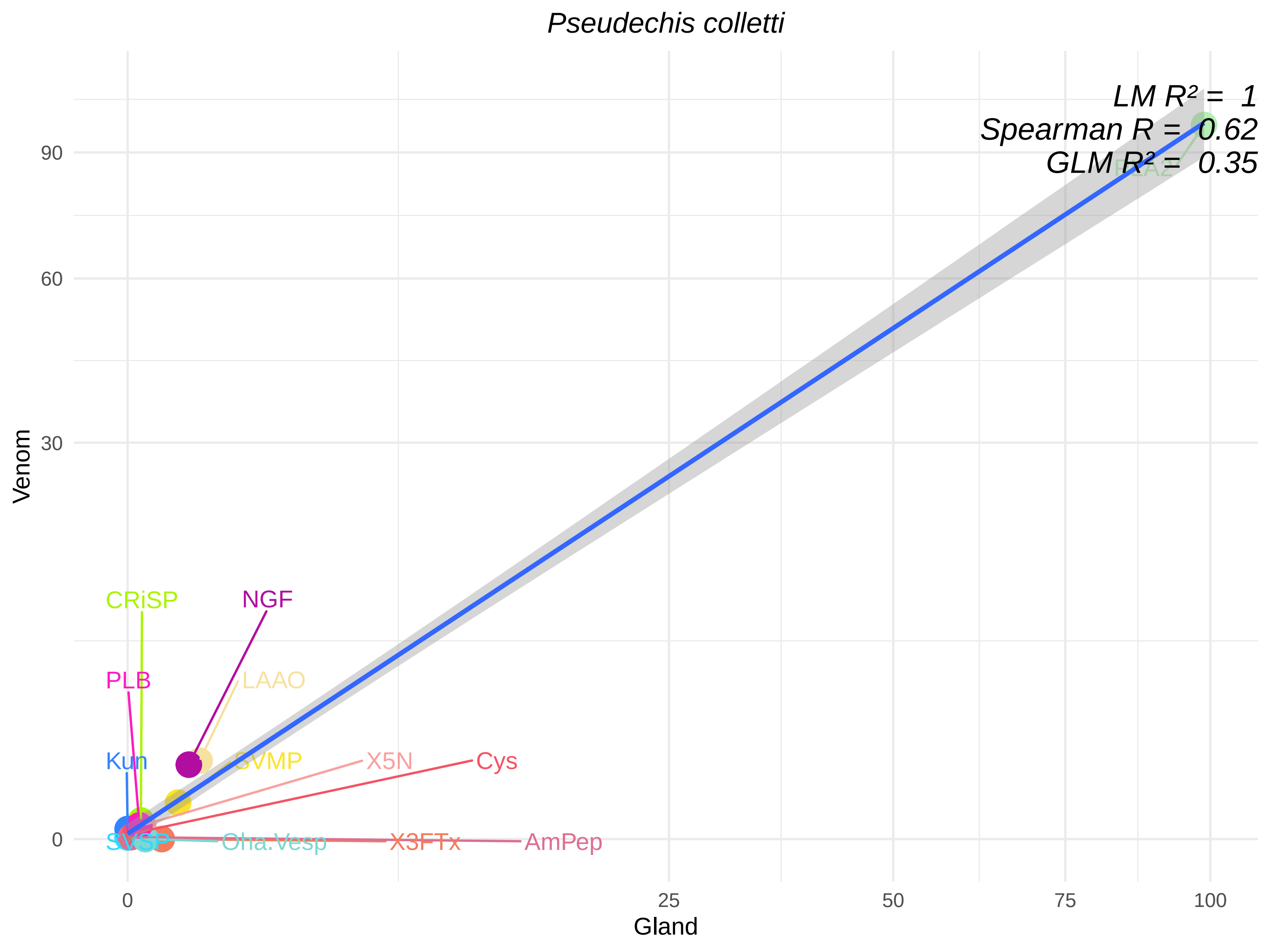

Pseudechis colletti
LM R² = 1
Spearman R = 0.62
GLM R² = 0.35
PLA2
CRiSP
NGF
PLB
LAAO
Kun
SVMP
X5N
Cys
SVSP
Oha.Vesp
X3FTx
AmPep
Venom
Gland
0
30
60
90
0
25
50
75
100

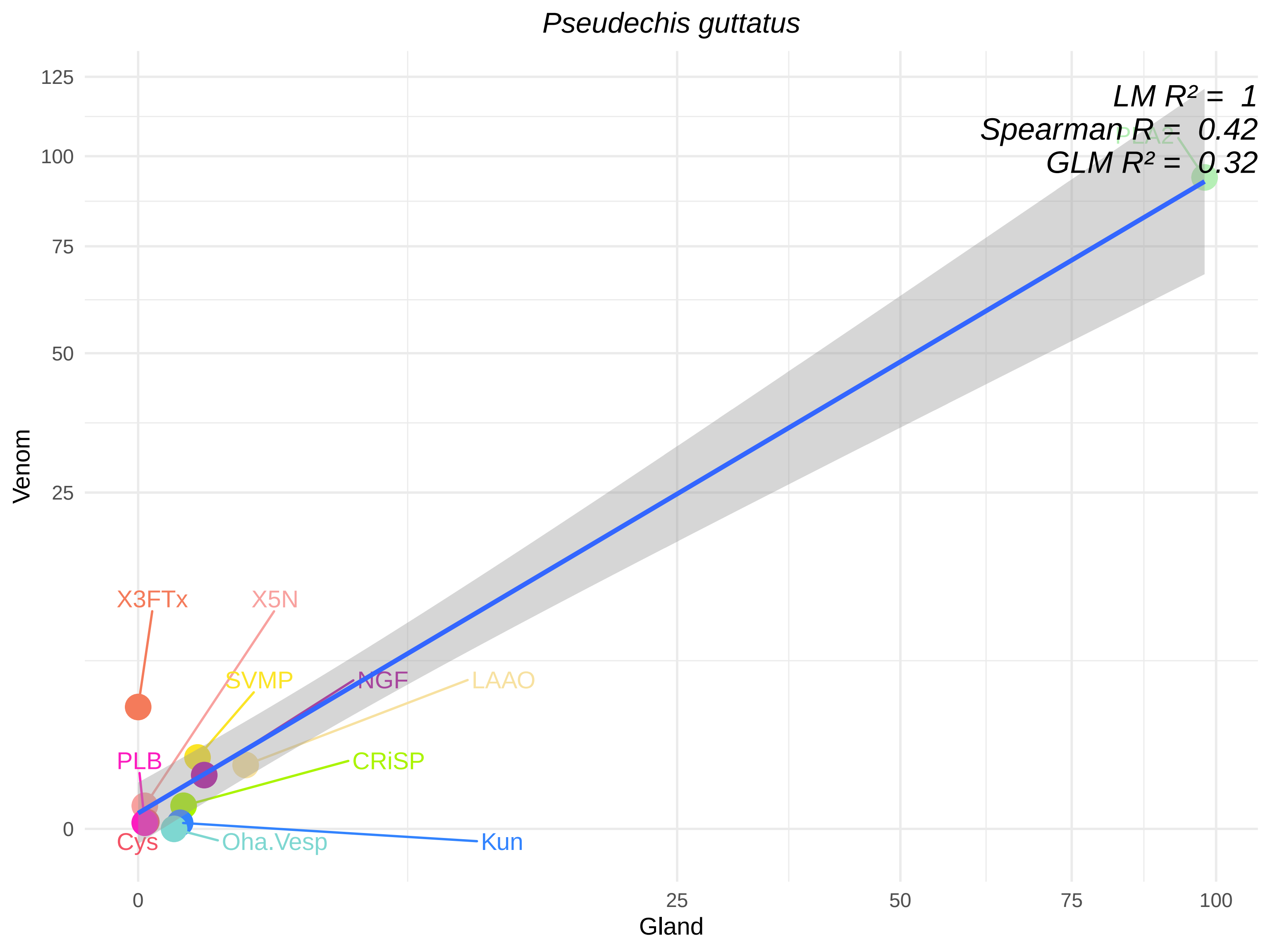
Pseudechis guttatus
LM R² = 1
Spearman R = 0.42
GLM R² = 0.32
PLA2
X3FTx
X5N
SVMP
NGF
LAAO
PLB
CRiSP
Cys
Oha.Vesp
Kun
Venom
Gland
0
25
50
75
100
125

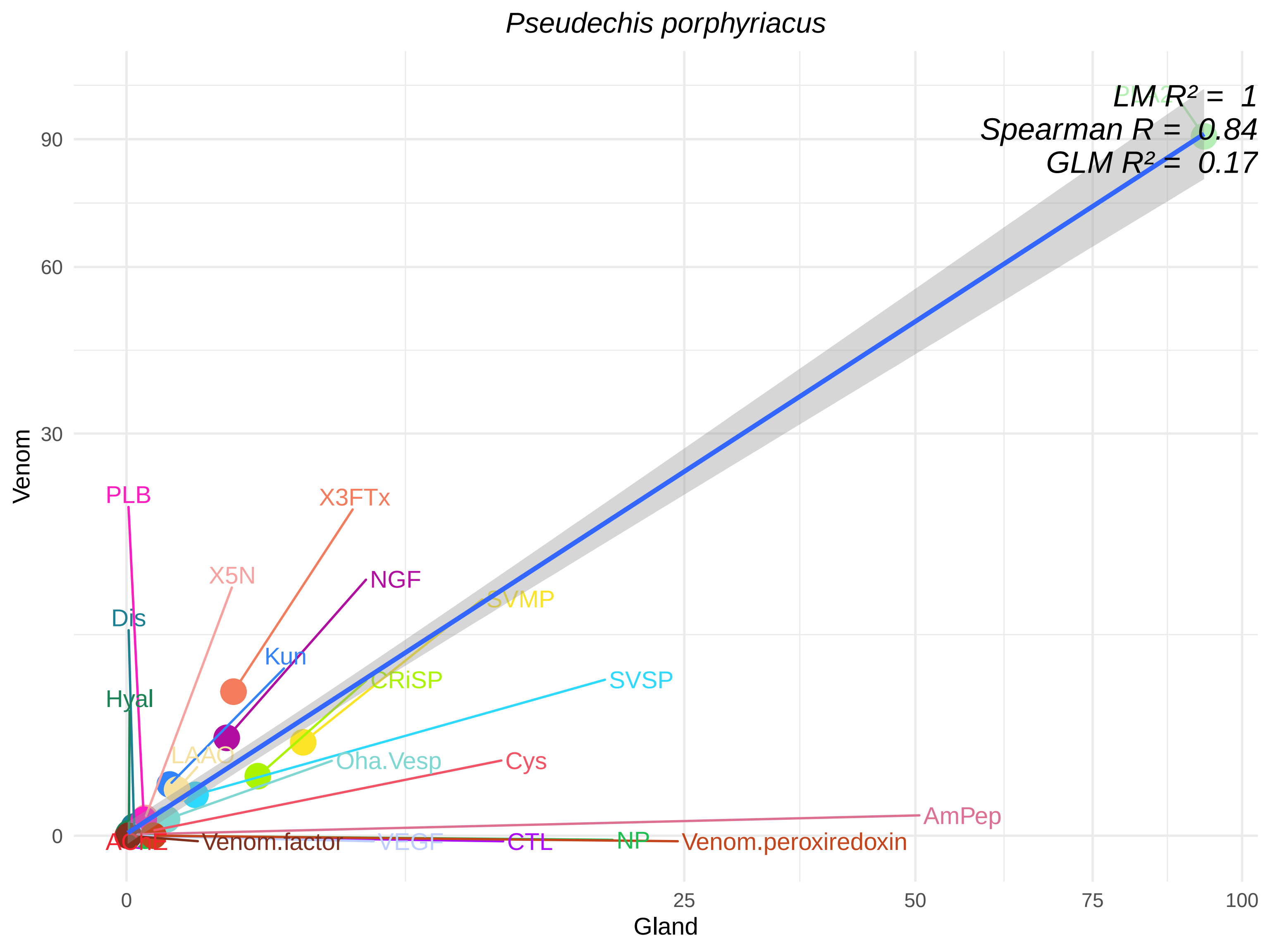

Pseudechis porphyriacus
LM R² = 1
Spearman R = 0.84
GLM R² = 0.17
Venom
Gland
0
30
60
90
0
25
50
75
100
PLA2
PLB
X3FTx
X5N
NGF
SVMP
Dis
Kun
CRiSP
SVSP
Hyal
LAAO
Oha.Vesp
Cys
AmPep
AChE
Venom.factor
VEGF
CTL
NP
Venom.peroxiredoxin

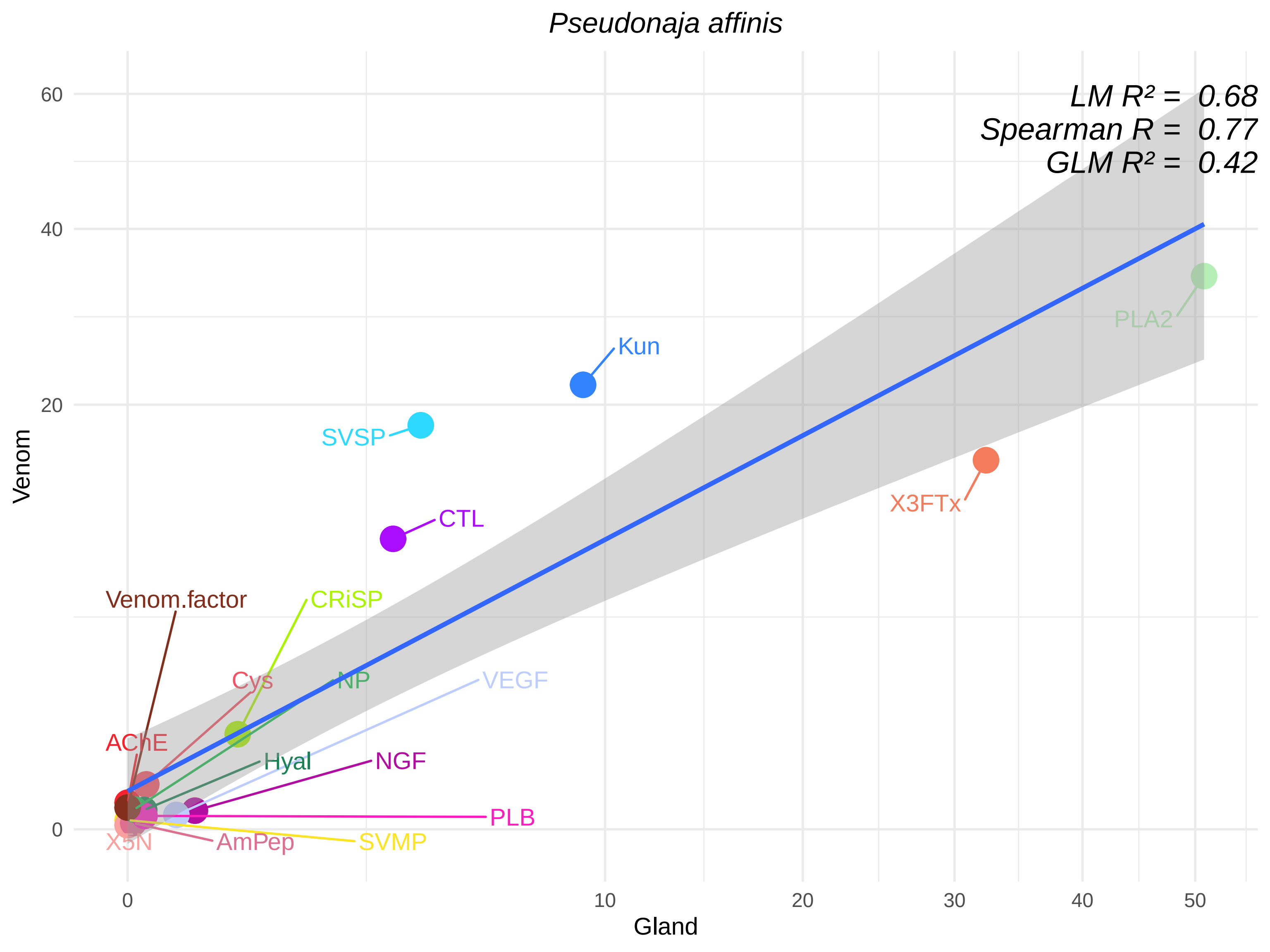

Pseudonaja affinis
LM R² = 0.68
Spearman R = 0.77
GLM R² = 0.42
Venom
Gland
60
40
20
0
0
10
20
30
40
50
PLA2
Kun
SVSP
X3FTx
CTL
Venom.factor
CRiSP
Cys
NP
VEGF
AChE
Hyal
NGF
PLB
X5N
AmPep
SVMP

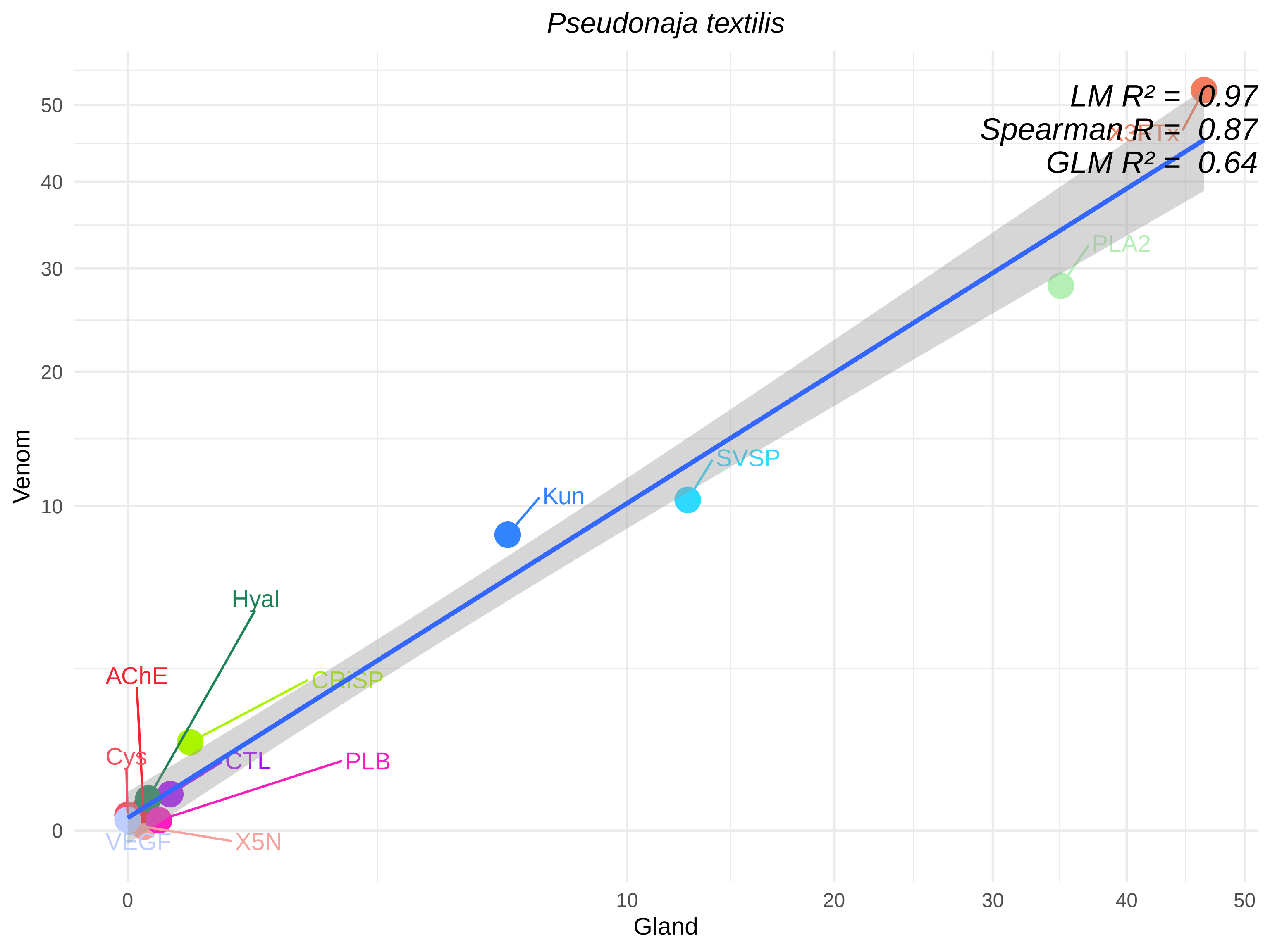

Pseudonaja textilis
LM R² = 0.97
Spearman R = 0.87
GLM R² = 0.64
X3FTx
PLA2
SVSP
Kun
Hyal
AChE
CRiSP
Cys
CTL
PLB
VEGF
X5N
Venom
Gland
0
10
20
30
40
50

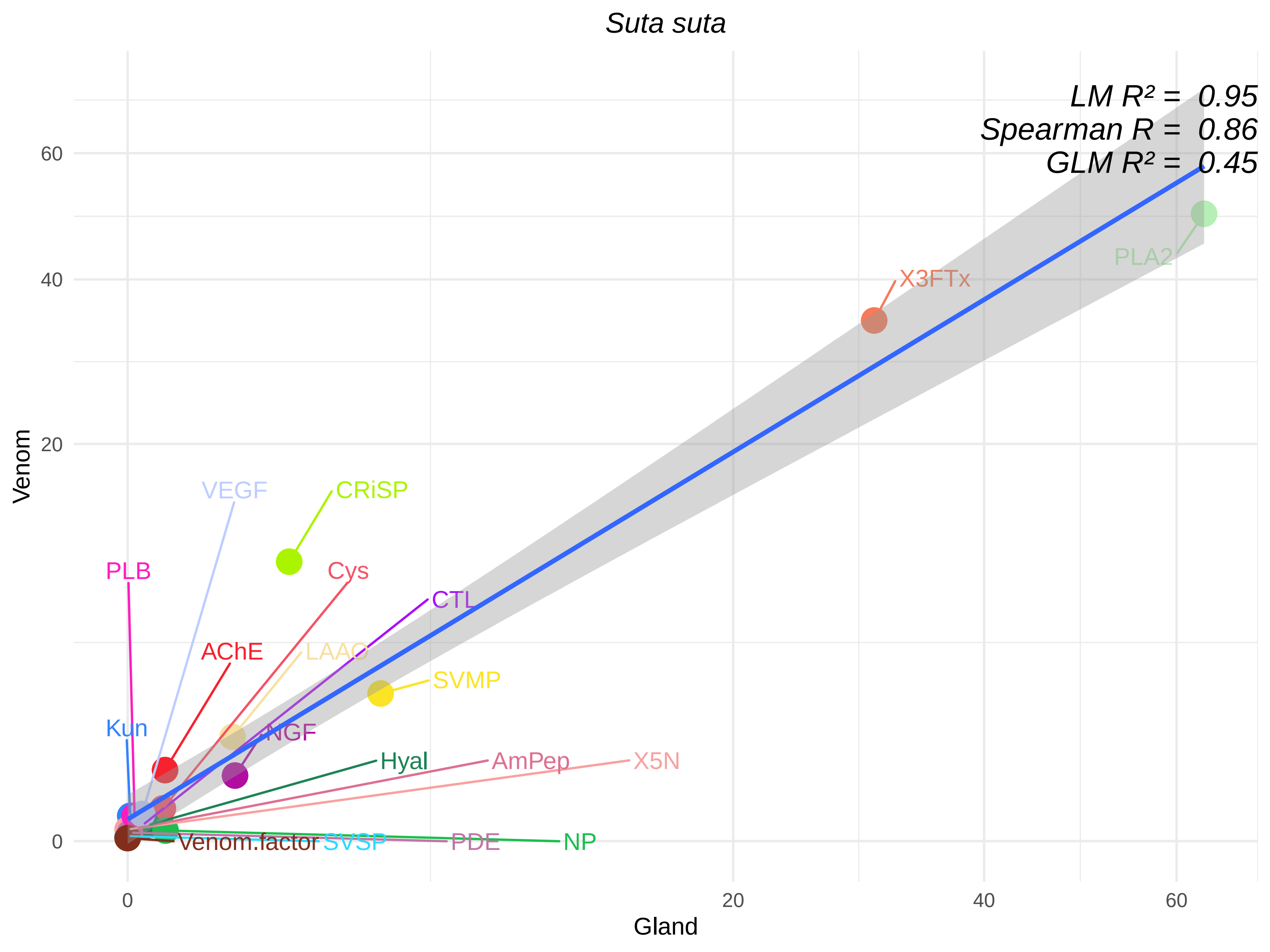
Suta suta
LM R² = 0.95
Spearman R = 0.86
GLM R² = 0.45
Venom
Gland
0
20
40
60
PLA2
X3FTx
VEGF
CRiSP
PLB
Cys
CTL
AChE
LAAO
SVMP
Kun
NGF
Hyal
AmPep
X5N
Venom.factor
SVSP
PDE
NP

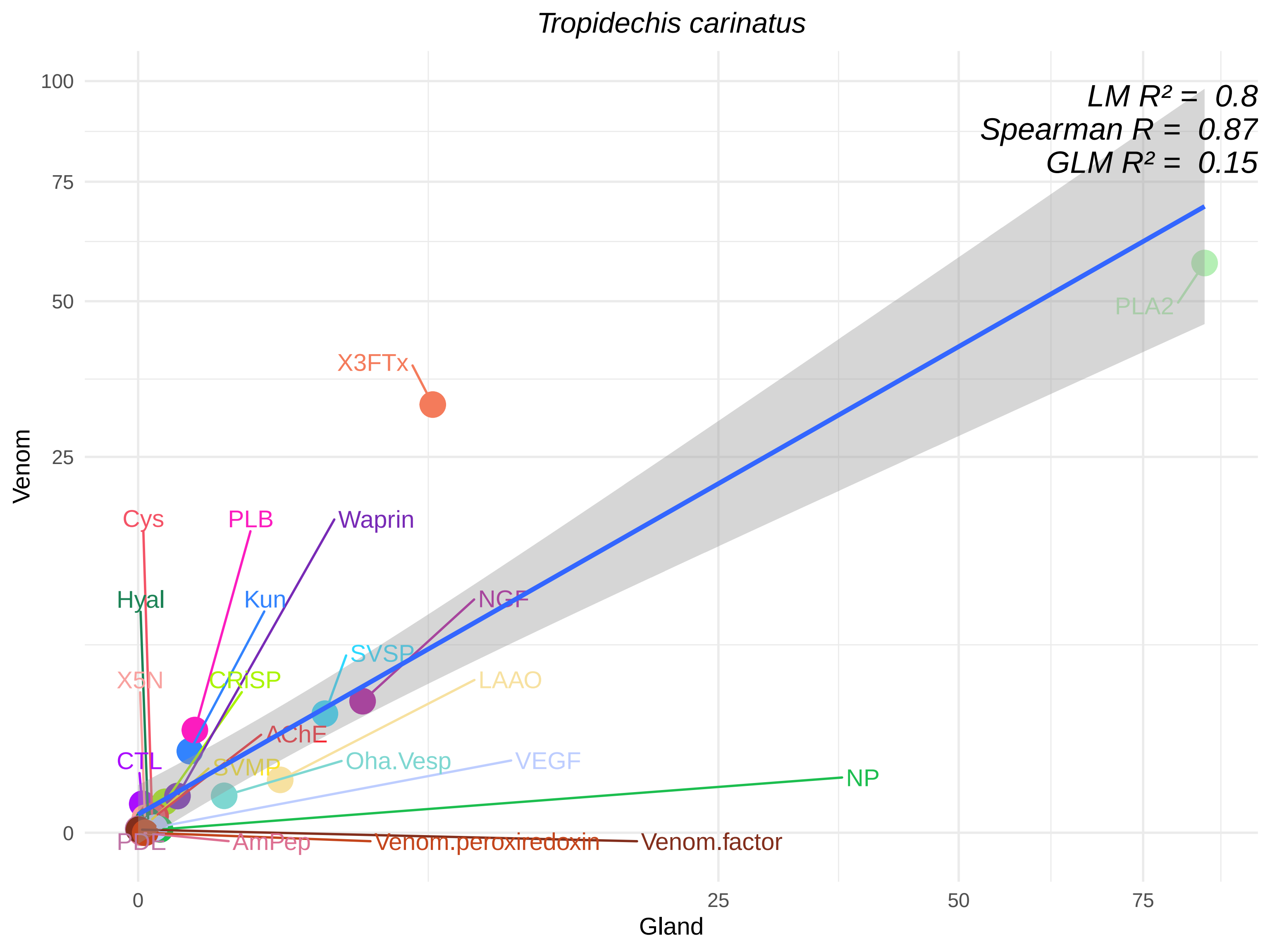

Tropidechis carinatus
LM R² = 0.8
Spearman R = 0.87
GLM R² = 0.15
Venom
Gland
100
75
50
25
0
0
25
50
75
PLA2
X3FTx
Cys
PLB
Waprin
Hyal
Kun
NGF
SVSP
X5N
CRiSP
LAAO
AChE
CTL
SVMP
Oha.Vesp
VEGF
NP
PDE
AmPep
Venom.peroxiredoxin
Venom.factor
